# Label-Free AI-Classification of Subcellular Organelles Based on Optical Photothermal Infrared Images

**DOI:** 10.64898/2026.06.02.729616

**Authors:** Michael J. Burke, Victor S. Batista, Caitlin M. Davis

## Abstract

Cells maintain homeostasis by dynamically reorganizing their organelles to tune metabolism in response to stress. Fluorescence microscopy maps organelle locations with subcellular resolution but provides limited information on their chemical composition. Infrared (IR) imaging offers a label-free alternative for probing intrinsic molecular vibrations that report on lipids, carbohydrates, and nucleic acids. However, its broader application to subcellular biology has been limited by spatial resolution and hyperspectral data complexity. Here, we combine submicron optical photothermal IR imaging with machine learning to classify subcellular structures in fixed U-2 OS cells. Using fluorescent-labeled organelles as ground truth, we trained and evaluated random forest (RF) classifiers and U-Net convolutional neural networks to identify organelles from IR spectra. The RF model converged rapidly, requiring fewer than 75 spectra per class per cell and fewer than 25 cells, indicating that models trained on small cellular regions can be extended to classify whole-cell images. The resulting classifiers accurately identified multiple organelles, including the endoplasmic reticulum, Golgi apparatus, mitochondria, nucleus, nucleolus, and stress granules. In contrast, classification was unsuccessful for nuclear speckles, actin, and microtubules, suggesting that some structures lack sufficiently distinct IR signatures under these conditions. Classifiers trained in U-2 OS cells generalized to HEK 293 cells, consistent with conserved organelle biochemical composition across cell types. However, the classifiers failed under cellular stress, indicating sensitivity to stress-induced changes in organelle state. Together, these results establish a scalable, label-free strategy for high-resolution mapping of organelle biochemical composition and provide a foundation for subcellular biomarker discovery and disease-state diagnostics.

## Introduction

Cells operate in a state of dynamic equilibrium, constantly adjusting their internal environment to maintain homeostasis. Disruptions can arise from imbalances of essential biological macromolecules (proteins, carbohydrates, lipids, and nucleic acids) or from their structural abnormalities, such as protein aggregation.^1,2^ Precise control over metabolite concentrations and their spatial distributions is essential not only for metabolic function but also for proper cell signaling.^3^ Organelles, the specialized compartments where metabolic chemistry occurs, depend on tightly regulated conditions, including pH, ionic balance, redox state, and membrane potential to function properly.^4^ However, these conditions are mostly missed by conventional fluorescence-based methods for visualizing subcellular structures. Such methods rely on stains of a specific chemical/physical property or label specific biomolecules that are used to report on the location of a target within the cell in a binary way. Additionally, fluorescent labels can be phototoxic to the cell or disrupt native molecular interactions.^5^

These limitations have driven interest in label-free imaging techniques that preserve the native state of the cell, particularly those that provide chemical contrast. A deeper understanding of the chemical composition of organelles is important due to the highly interconnected nature of metabolic pathways, where dysregulation in a pathway housed in one organelle can propagate across the cell, affect neighboring cells, and lead to tissue damage and disease. For example, impaired protein folding and lipid synthesis within the endoplasmic reticulum (ER) are linked to neurodegeneration, type 2 diabetes, and liver disease.^6–10^ Beyond the function of individual organelles, inter-organelle coordination, particularly through membrane contact sites, is essential for maintaining cellular homeostasis.^11^ For instance, mitochondria-lysosome crosstalk contributes to their co-dysfunction observed in Alzheimer’s and Parkinson’s diseases.^12–14^ Therefore, a comprehensive understanding of subcellular physiology and pathology of organelles requires concomitant characterization of organelle location, biomolecular composition and structure, and local environmental conditions.^4,15,16^

Vibrational Raman and infrared (IR) imaging offer label-free, nondestructive readouts of chemical composition and molecular structure.^17,18^ By probing intrinsic molecular vibrations, these methods enable identification of biochemical constituents, while band shifts report on changes in molecular conformation and local chemical environment.^19–24^ However, each vibrational imaging technique comes with trade-offs. Raman-based approaches have enabled subcellular classification, but their broader application is limited by long acquisition times and comparatively weak sensitivity to polar bonds (e.g. C-O, C-N, and N-H), which are abundant in biological systems.^25–28^ Conventional IR microscopy, in contrast, is poorly suited for subcellular analysis because mid-IR measurements are limited by strong water absorption and by optical diffraction, yielding spatial resolutions typically on the micrometer scale.^29^ Consequently, IR imaging has been used primarily for classification at the whole cell or tissue level.

Recent photothermal IR methods have begun to address these barriers. Optical photothermal infrared (OPTIR) imaging and photothermal atomic force microscopy-based IR (AFM-IR) now enable submicron IR measurements.^30^ AFM-IR can achieve spatial resolutions below 20 nm, but its slow acquisition rate limits its utility for imaging extended cellular regions. OPTIR, by contrast, can acquire sub-micron resolution images in real time in some configurations.^31^ Although OPTIR has already been used to classify cell types and detect disease biomarkers in large-scale tissue specimens, its use for organelle-level classification within individual cells has not yet been demonstrated.^32,33^

In this study, we combine OPTIR imaging with machine learning models to classify subcellular organelles in fixed U-2 OS cells. We developed two complementary models tailored to distinct acquisition modes: single-spectrum measurements and hyperspectral image collections. For single-spectrum data, we trained a random forest (RF) classifier to rapidly assign cellular location directly from IR spectra. This model enables real-time spectral identification and can guide subsequent OPTIR image acquisition. For hyperspectral datasets, which provide spatially resolved chemical maps of organelle states, we evaluate both the RF classifier and a U-Net convolutional neural network. Although the RF model can be applied to hyperspectral data, the U-Net improves prediction accuracy by incorporating spatial relationships among neighboring pixels. Because this spatial context is intrinsic to its architecture, the U-Net is restricted to hyperspectral maps. To identify the minimal set of IR spectral features required for organelle classification, we performed high-resolution spectral mapping across selected subcellular regions. While the collection of hyperspectral datasets on our OPTIR instrument exceeds the time constraints for live-cell studies, the use of a real-time OPTIR instrument and/or a reduced feature set may ultimately enable real-time organelle identification in living cells. We demonstrate that both the RF classifier and a U-Net convolutional neural network can reliably distinguish organelles, including the ER, Golgi apparatus, mitochondria, nucleus, nucleolus, and stress granules. These models also generalize to HEK 293 cells, demonstrating the potential for broader application across cell lines. Additionally, we observe changes in prediction accuracy based on cell dysfunction, which can have profound cellular and systemic consequences.^34–37^ Spectral signatures can be interpreted to understand compositional changes in the dysfunctional state. These findings highlight the importance of tools that can resolve organelle-specific changes with high precision, while providing valuable chemical information. Our machine learning-based approaches offer a promising strategy for investigating organelle function and detecting disease-associated biomarkers at the subcellular level.

## Results

### Training Dataset Acquisition

Accurate organelle classification depends on a representative training dataset that captures the spectral diversity of subcellular regions. This involves three key steps: (1) optimizing spectral acquisition conditions to achieve a signal-to-noise ratio (SNR) high enough to resolve organelle-level differences; (2) identifying the specific regions of the OPTIR spectrum that contain the most discriminative information for classification; and (3) determining the minimal dataset size required for effective model training.

Our OPTIR system, equipped with both fluorescence and IR capabilities, enables the collection and assignment of fluorescence and IR data at the same subcellular locations. The training dataset was generated from U-2 OS cells fixed with 4% paraformaldehyde (PFA), a standard fixative for fluorescence staining that has been reported to minimally affect IR spectra.^38^ Cells were stained with organelle-specific dyes or antibodies to generate ground truth maps for organelle identification (Fig. 1). These stains, applied at nanomolar (nM) concentrations, are not detectable by IR imaging (Fig. S1).^39^ Where possible, dye-based labels were used to minimize potential spectral interference from fluorescent proteins or antibody stains with native protein populations.

**Figure 1.**
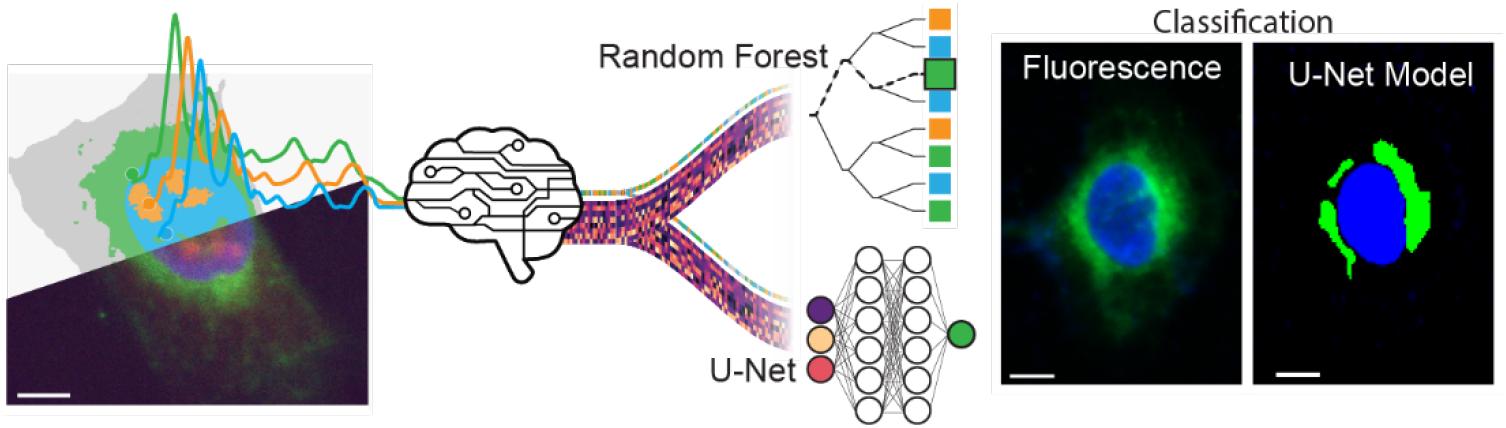
Machine learning-based classification of subcellular structures. U-2 OS cells were cultured on CaF_2_ coverslips, fixed with 4% PFA, and stained with organelle-specific fluorescent markers. Fluorescence labeled IR datasets were then collected using OPTIR. These datasets were systematically analyzed to identify the minimal set of unique parameters necessary to distinguish organelles within the training set, which comprises multiple cells. The extracted parameters were used for two models, including a Random Forest classifier and a U-Net convolutional neural network. Model accuracy was assessed by evaluating the ability of the algorithms to classify organelles in independent test data not included in the training set. An example of the classification output shows the model accurately identifying Golgi (green) and nucleus (blue) in comparison to the fluorescence.

The signal-to-noise ratio (SNR) in the dataset is governed by both sample composition and acquisition conditions. According to the Beer-Lambert law, signal strength depends on the optical path length, molar absorptivity, and concentration of cellular components at each frequency. As a result, signal strength is highest in thicker regions of the cell, particularly around the nucleus, and decreases toward the periphery. Noise in OPTIR measurements arises primarily from atmospheric water absorption and the instrument. To mitigate atmospheric interference, the sample chamber is purged with dry air. Our instrument uses a four-module tunable MIRcat QCL laser, which exhibits increased noise at module transitions, most significantly in the third module (1185 to 1280 cm^-1^), where laser power is lower (Fig. S2). Accordingly, spectral data from this range are less reliable and thus were not used in parameter selection.

To determine parameters for organelle classification, we compared IR spectral features across organelle classes and selected those with minimal inter-class overlap. To facilitate use in future studies, IR features were selected to reflect known biochemical distinctions between organelles (Table S1). To reduce redundancy, we computed pairwise correlations between parameters (Fig. S3) and excluded those that consistently correlated across classes. The final non-redundant parameter sets were used as inputs for our classification models. To demonstrate our approach, we first showcase its ability to distinguish the nucleus from the surrounding cytoplasm. We then extend the analysis to other cell lines and subcellular structures, including the nucleolus, ER, Golgi apparatus, mitochondria, and stress granules.

### Optimization of the AI Classification Approach: Distinguishing Nucleus from Cytoplasm

Since the nucleus and cytoplasm can be distinguished using IR spectra from diffraction-limited IR microscopes,^17^ we used these compartments to develop and validate our approach. We developed two models, a RF classifier for single-spectrum collection and a U-Net for hyperspectral images. The nuclear training dataset comprised of 7,031 OPTIR spectra sampled across 55 cells. Spectra were averaged, background corrected, and normalized to the amide I band. Seven IR features, representing characteristic absorptions from the four major classes of biological macromolecules, were selected as input parameters for machine learning (Fig. 2):

**Figure 2.**
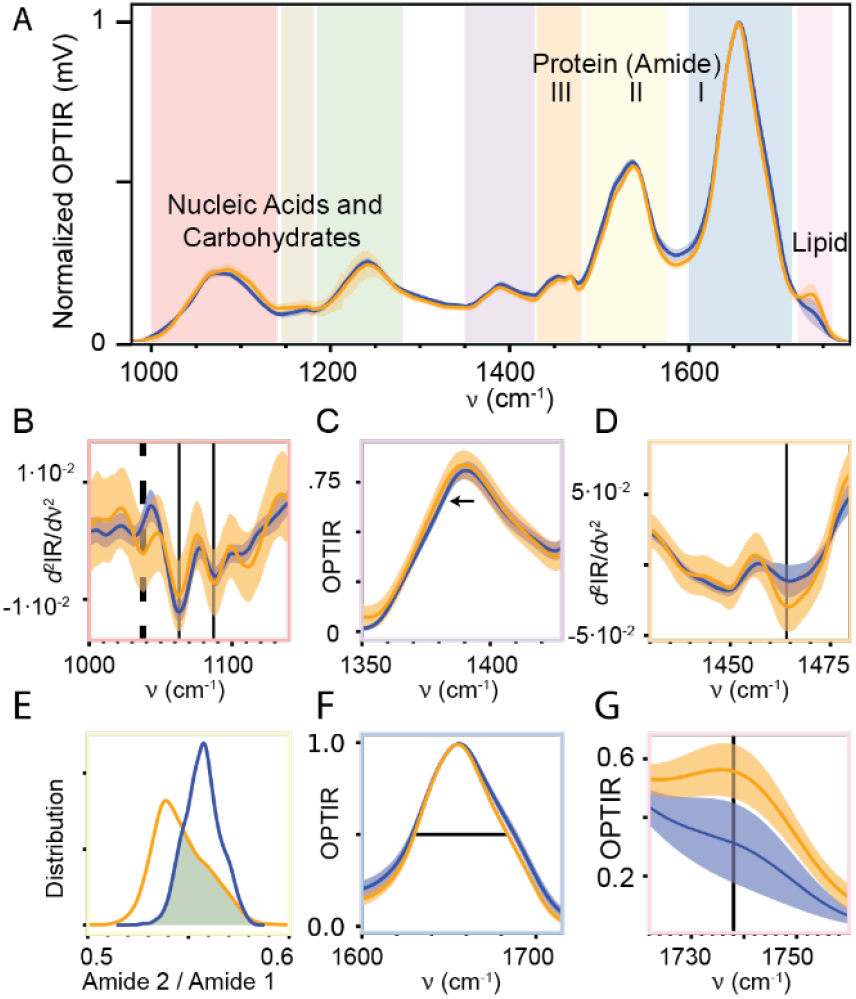
IR parameters for machine learning. **A. Spectral comparison between nucleus and cytoplasm**: The average IR spectrum of the nucleus (blue) is compared to that of the cytoplasm (orange) within a hyperspectral dataset for a single cell region with 7,040 pixels. The shaded areas represent the standard deviation of the spectra. The entire spectrum is divided into eight regions for detailed analysis. No features from the second (brown) or third (green) regions were selected as machine learning parameters for distinguishing the nucleus from the cytoplasm. **B. Discriminative parameters from the first region (red):** A second derivative analysis of the first spectral region (red) highlights two key parameters used to distinguish between the nucleus (blue) and cytoplasm (orange). First, cytoplasm-specific band at 1040 cm^-1^ (black dashed line) is present in cytoplasm but absent in the nucleus. Second, the ratio of absorption peaks 1063 cm^-1^/1087 cm^-1^ (black solid lines) is significantly different between the nucleus and cytoplasm. **C. Shift in the fourth region (purple):** A redshift at the 1390 cm^-1^ band is observed in the cytoplasm compared to nucleus. **D. Enhanced absorption in the fifth region (orange):** The cytoplasm exhibits an increased absorption at 1464 cm^-1^ relative to the nucleus. **E. Amide II/I ratio (sixth and seventh regions):** The ratio between Amide II (1450 cm^-1^, sixth yellow region) and Amide I (1650 cm^-1^, seventh blue region) differentiates the nucleus from the cytoplasm. **F. Amide I peak width variation (seventh region):** The FWHM of the Amide I peak is broader in the nucleus compared to the cytoplasm. **G. Lipid signal in the eighth region (pink):** The cytoplasm displays a higher lipid band at 1739 cm^-1^ compared to the nucleus.

#### Carbohydrates

The cytoplasm contains a high concentration and diversity of carbohydrates due to its central roles in carbohydrate storage and energy metabolism.^40^ In contrast, in the nucleus, carbohydrates are largely limited to the sugar-phosphate backbone of DNA.^41^ Carbohydrate-specific C-O stretching modes of carbohydrates (1000-1300 cm^-1^) appear within the IR fingerprint region. Second-derivative analysis of this region reveals clear spectral differences between the nucleus and cytoplasm (Fig. 2B). Notably, a distinct band at 1036 cm^−1^ is present in the cytoplasm but absent in the nucleus, likely corresponding to the C– O stretches of primary alcohol groups in sugars, particularly glucose.^42^

#### Nucleic acids

The overall nucleic acid content in the nucleus and cytoplasm is comparable. However, their composition differs: DNA is predominantly confined to the nucleus, whereas RNA is distributed between the nucleus and cytoplasm.^17,43^ As a result, both regions exhibit symmetric and asymmetric phosphate stretching bands at ∼1086 cm^-1^ and ∼1220 cm^-1^, respectively.^44^ Nevertheless, nucleus and cytoplasm can be differentiated because the IR band positions and intensities are also sensitive to the nucleic acid secondary structure. For example, the weak furanose ring stretch that appears as a shoulder near the symmetric phosphate band in B-form DNA becomes significantly enhanced in Z-form DNA.^45^ Indeed, in the second derivative OPTIR spectra, bands at 1063 and 1087 cm^−1^ are observed, with a notably stronger peak at 1063 cm^−1^ in the nucleus consistent with the presence of Z-form DNA (Fig. 2B).

#### Lipids

Lipid concentration is generally higher in the cytoplasm than in the nucleus,^46^ reflecting the increased lipid metabolism, storage, and the presence of membrane-bound organelles in the cytoplasm.^47^ The strongest lipid band detected by our OPTIR instrument corresponds to the C=O stretching of ester carbonyls in triglycerides and cholesterol esters, centered at 1740 cm^-1^.^48^ Although vibrational modes of other biomolecules overlap with strong lipid bands in the fingerprint region between 1425 and 1490 cm^-1^, the dominant contributions in this range arise from CH_3_ bending and CH_2_ scissoring modes originating from lipids.^49,50^ Reflecting the higher cytosolic lipid content, the signal intensity is higher at both 1462 cm^-1^ (Fig. 2D) and 1740 cm^-1^ (Fig. 2G).

#### Proteins

Because the hyperspectral data are normalized to the amide I region, the relative protein concentration in the nucleus and cytoplasm cannot be used as a distinguishing feature. Instead, the amide II/amide I ratio can serve as a proxy for differences in protein secondary structure, which affect the hydrogen bonding network in the amide II.^21^ This ratio is commonly used to differentiate between primary and malignant cells. We hypothesize that structural and compositional differences in proteins between the nucleus and cytoplasm would manifest as distinct amide II/amide I ratios. Consistent with this, the nucleus displays a higher amide II/amide I ratio compared to the cytoplasm (Fig. 2E). Additionally, secondary structure variations alter the shape of the amide I band,^51,52^ which we captured through measurements of its full-width at half maximum (FWHM) (Fig. 2F).

Carboxylate groups are found in all four major classes of biological macromolecules as well as in numerous metabolites involved in cell metabolism. The symmetric COO^-^ stretching vibration appears as a distinct band at 1390 cm^-1^.^53^ A red shift of this band in the cytoplasm relative to the nucleus reflects differences in the composition of carboxylate-containing molecules and variations in the local environment (Fig. 2C).

Fully establishing the molecular origins of spectral differences between organelles is challenging because there is significant overlap of absorptions within and across the major biomacromolecule classes. This is further convoluted by the environmental sensitivity of IR absorptions. Validation of any assignment is complicated, however, recently organelle assignments were precisely made by Raman imaging and the same approach could be used to validate IR parameters in the future.^54^

Our initial classification method leverages a RF model to distinguish between nucleus and cytoplasm based on the IR spectral features described above.^55^ The key advantage of this method is its ability to classify individual spectra independently, without relying on spatial information. Additionally, the model outputs confidence scores for each prediction, providing a built-in measure of classification certainty and enabling performance assessment.

Given the time-intensive nature of acquiring full hyperspectral maps for training, minimizing data collection while maintaining high classification accuracy is critical. We tested methods that reduce the number of pixels where data is collected while maintaining the number of datapoints within each IR spectrum, so that the same spectral background correction can be applied across all training sets. For intra-cell classification, sampling as few as 5 pixels each for the nucleus and cytoplasm yields 94 ± 1% accuracy for predictions within that cell (Fig. S4). However, inter-cell variability presents a greater challenge. A model trained on an equivalent amount of data, 5 pixels per class, but then tested on another cell reduces model accuracy to 81% (Fig. S5). To determine the optimal training dataset size necessary for robust, cross-cell classification, it is necessary to optimize both the required number of examples per cell and the total number of cells. To determine the required spectra per cell, we took a single hyperspectral map and randomly sampled an increasing amount of data per class for training a RF model. These models were then tested on nine different cells. To control for sampling bias, the process was repeated 250 times at each training set size. While accuracy and consistency improved with larger training sets, most of the benefit was seen with the first 25 training spectra of each class per cell before plateauing at 50–100 training spectra of each class per cell (Fig. S5).

To determine the required number of cellular examples in the training dataset, we used a similar strategy. Using a dataset of 55 cells, between 1 and 40 cells were randomly selected for training and tested on spectra from 5 randomly sampled cells that were not included in the training set (Fig. S6). By fitting a learning curve to this data, we estimated a theoretical maximum classification accuracy possible with the current method, which closely matched empirical performance (Table 1). Based on these results, we standardized training datasets for other organelles to include 50-150 spectra per cell (25-75 for each class) and 15-25 cells. During training we also determined that collecting negative class examples of both the cytoplasm and nucleoplasm is important because they represent distinct negative examples.

**Table 1.**
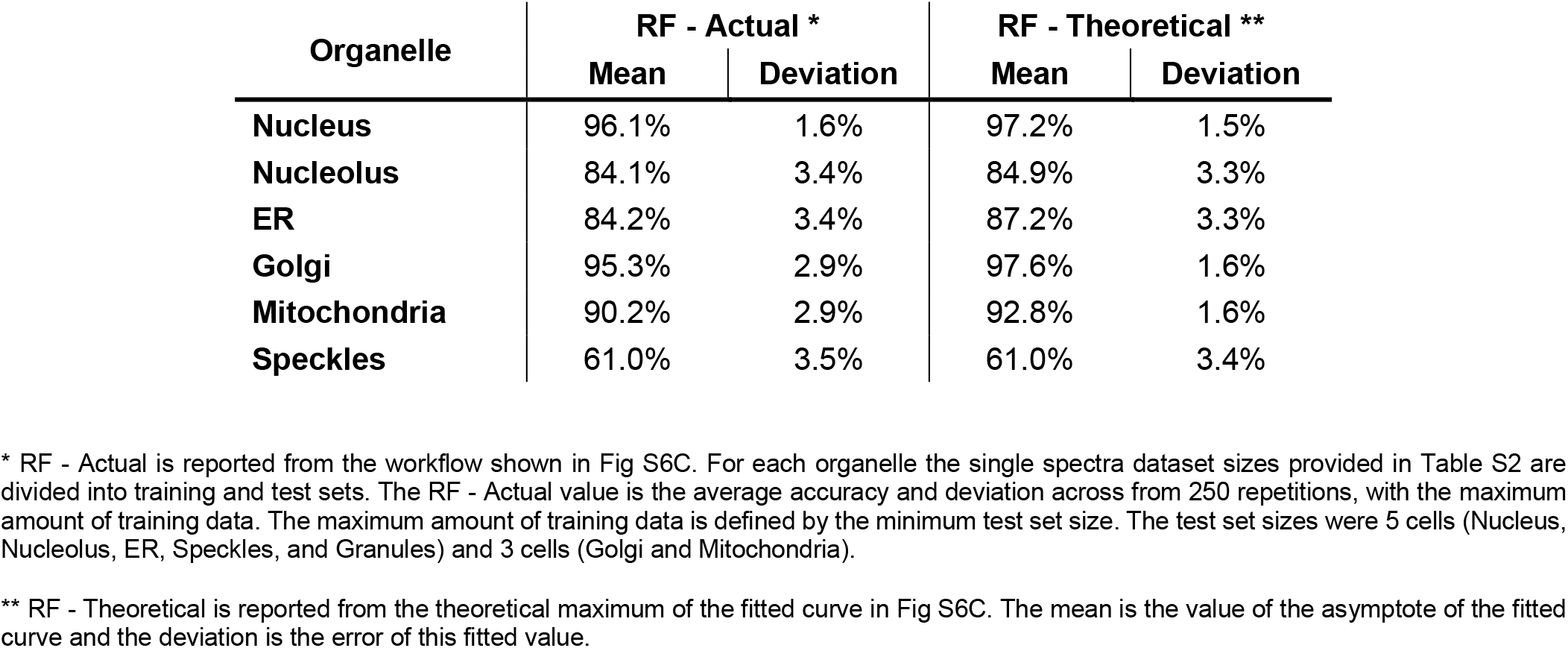
Random Forest Training Accuracies.

| Organelle | RF - Actual * |  | RF - Theoretical ** |  |
| --- | --- | --- | --- | --- |
|  | Mean | Deviation | Mean | Deviation |
| **Nucleus** | 96.1% | 1.6% | 97.2% | 1.5% |
| **Nucleolus** | 84.1% | 3.4% | 84.9% | 3.3% |
| **ER** | 84.2% | 3.4% | 87.2% | 3.3% |
| **Golgi** | 95.3% | 2.9% | 97.6% | 1.6% |
| **Mitochondria** | 90.2% | 2.9% | 92.8% | 1.6% |
| **Speckles** | 61.0% | 3.5% | 61.0% | 3.4% |

\* RF - Actual is reported from the workflow shown in Fig S6C. For each organelle the single spectra dataset sizes provided in Table S2 are divided into training and test sets. The RF - Actual value is the average accuracy and deviation across from 250 repetitions, with the maximum amount of training data. The maximum amount of training data is defined by the minimum test set size. The test set sizes were 5 cells (Nucleus, Nucleolus, ER, Speckles, and Granules) and 3 cells (Golgi and Mitochondria).
\*\* RF - Theoretical is reported from the theoretical maximum of the fitted curve in Fig S6C. The mean is the value of the asymptote of the fitted curve and the deviation is the error of this fitted value.

Because training and testing datasets were selected based on fluorescent markers for clear nuclear or cytoplasmic identity, the 96.1% classification accuracy overestimates model performance on full hyperspectral images. When applied across entire images, accuracy dropped to 90.2% (Table 2), with errors concentrated in regions of low SNR, e.g., distal cytoplasm and at the nucleus-cytoplasm interface where spectral overlap increases (Fig. 3). To address these limitations, we implemented a U-Net architecture, which incorporates both spectral and spatial information using convolutional and pooling layers for encoding and decoding.^60^ The U-Net architecture requires substantially more training data than the RF model, not just in terms of spectral diversity, but also in terms of the appearance (shape) and subcellular localization of each class. To meet this need, we generated synthetic cellular images using fluorescent images as templates, populated with spectra from the training set (see SI Methods, Fig. S7). This approach is effective when the required amount of training data exceeds the amount of data that can be reasonably collected.^56^

**Table 2.**
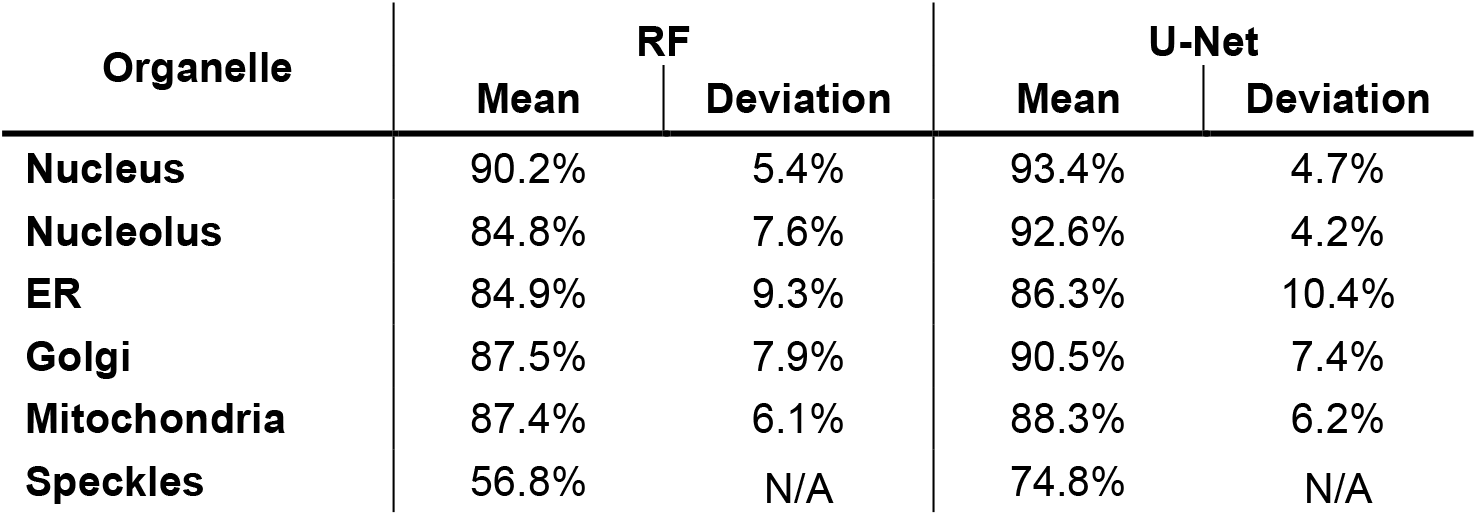
Average Classification Accuracies Across Partial and Full Cell Images.

**Figure 3.**
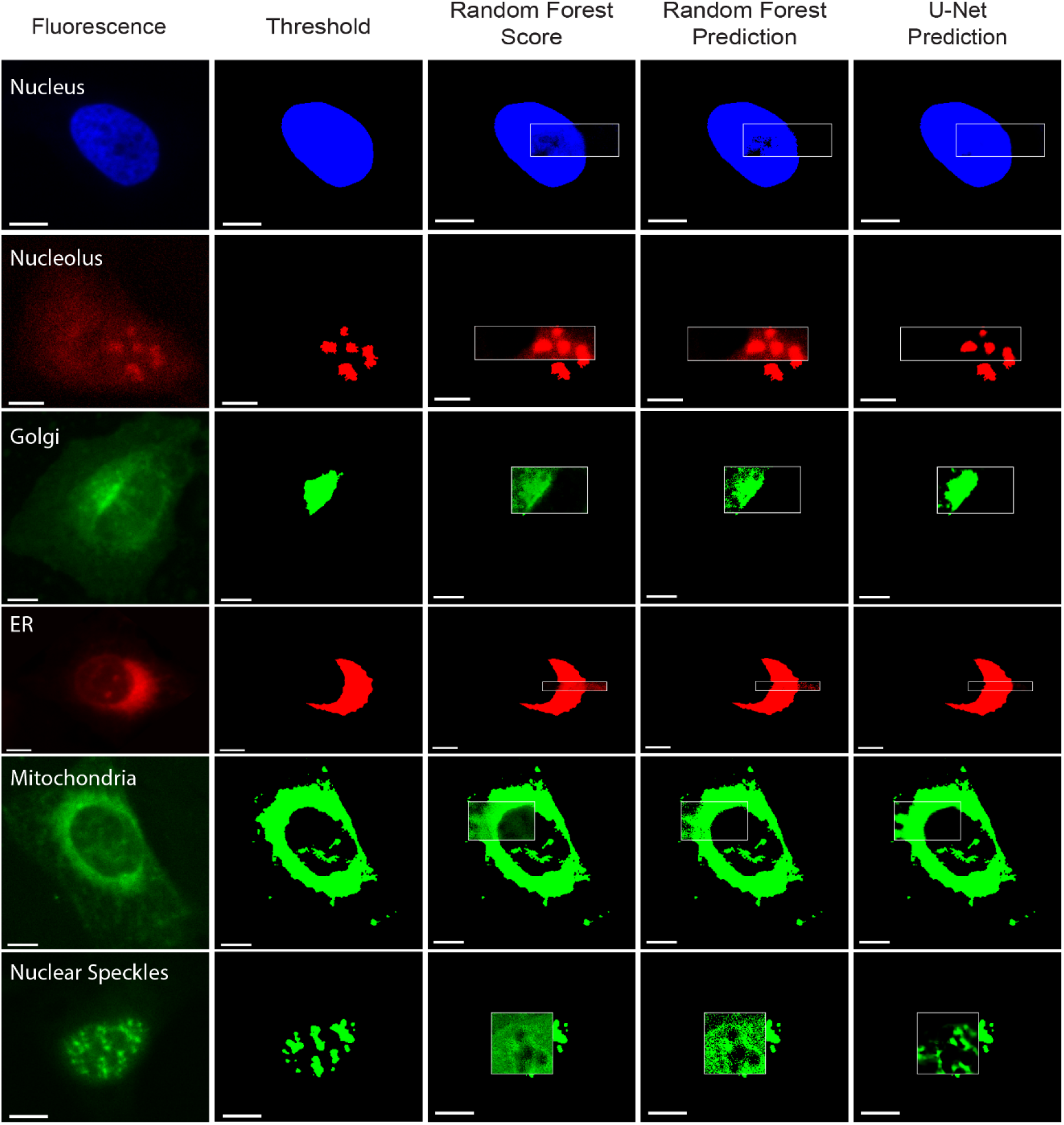
Random Forest and U-Net classification of organelles in cells not included in the training set. Representative classification of subcellular organelles using the random forest and U-Net machine learning approaches. The two leftmost columns display fluorescence images (scale bars: 10 μm) with organelle specific staining: nucleus (blue), nucleolus (red), nuclear speckles (green), ER (red), mitochondria (green), and Golgi apparatus (green). A white box is drawn around the region where OPTIR data was collected and where we show the model classification results. The third column shows the random forest confidence scores, which provide information on the certainty of the random forest model. The fourth column shows classification results generated by the random forest model. The final column presents predictions from the U-Net model, which was trained using a combination of random forest scores and IR features. Training dataset sizes are provided in Table S2.

We evaluated three types of input data for the U-Net: (1) the seven IR features alone, (2) RF confidence scores alone, and (3) a combination of both. All models were trained using the same set of pixels used to train the RF model. Among the three, the combined input of IR features and RF confidence scores achieved the highest accuracy and lowest variance across model replicates (Fig. S8). Qualitatively, this hybrid approach corrected many of the RF misclassifications and produced segmentations more closely aligned with the fluorescence-based ground truth (Fig. 3). Quantitatively, the U-Net improved nuclear classification accuracy by an average of 3.2% over the RF model (Table 2).

### Parameter Search for Label-Free Classification of Other Organelles

To extend the machine learning framework to classify multiple subcellular structures beyond the nucleus and cytoplasm, we collected hyperspectral data from subcellular regions containing each organelle of interest. RF predictions showed strong correlations with fluorescence images for the nucleolus, Golgi apparatus, ER, nuclear speckles, and mitochondria (Fig. 2). Since the accuracy is dependent on the fluorescent threshold, we have provided our fluorescent thresholds next to our predictions. Additionally, all hyperspectral images and the predictions for the organelle set (both the stained and unstained organelles) are provided in the supplementary materials (Fig. S17-S29). Attempts to classify actin and microtubules were not successful, likely due to their low concentration, making detection within a single voxel challenging (Fig. S9).

The U-Net model substantially improved classification performance by correcting errors present in RF predictions (Fig. 3). However, during training, U-Net models frequently became trapped in local minima. To address this, we employed a parallel training strategy: multiple models were trained simultaneously, with early termination for runs that failed to reach a predefined accuracy threshold within the first 10 epochs (see Methods). Final models were selected from the successful runs based on two criteria: (1) a low overall rate of false positives and negatives, and (2) a balanced distribution between false positive and negative rates (Fig. S10). This strategy yielded models capable of accurately classifying the nucleus, nucleolus, Golgi, ER, and mitochondria. However, nuclear speckles classification remains challenging; while the model has few false positives in cytoplasmic and nucleolus regions, it struggles to separate speckles from the surrounding nucleoplasm due to the high level of spectral similarity. In general, the most significant spectral differences between organelles were observed between nuclear and cytoplasmic structures, which are visually grouped into purple (nucleus) and green/blue (cytoplasmic) average (Fig. S11). This distinction is expected, as each voxel may contain mixed contributions from adjacent nuclear and cytoplasmic regions. Within each compartment, spectral distinctions between specific organelles are more subtle. Nonetheless, many organelles exhibit unique spectral signatures. For example, the nucleolus exhibits a pronounced peak at 1643 cm^-1^, suggesting a distinctive protein composition, potentially enriched in beta-sheet structures (Fig. S11G).

### Full Cell Organelle Classification

Next, we tested the ability of models optimized on cellular subsections to classify images of whole cells. Hyperspectral maps of three full cells were collected at 0.5 μm resolution. The spatial resolution was lowered from the 0.2 µm spatial resolution used in training to increase the imaging speed. In the full cell context, both RF and U-Net predictions show a strong correlation to the fluorescence image thresholds (Fig. 4, Fig. S27-29). The trend of the U-Net fixing errors present in the RF model continues, especially in the Golgi RF model which had a high tendency to misclassify cellular edges as falsely positive. On the other hand, the RF scores show a striking resemblance to the original fluorescent images. This suggests that the RF uncertainty score is tracking the amount of organelle present within each voxel. The RF prediction then makes a binary decision similar to our fluorescent thresholds. Alternatively, the U-Net can then act as an intelligent threshold that takes into account spatial context and corrects deeper errors within the parameter set. Average accuracies and the deviation across all tested hyperspectral images are provided in Table 2, with more detailed performance metrics for each cell provided in Fig. S27-29 Panel B.

**Figure 4.**
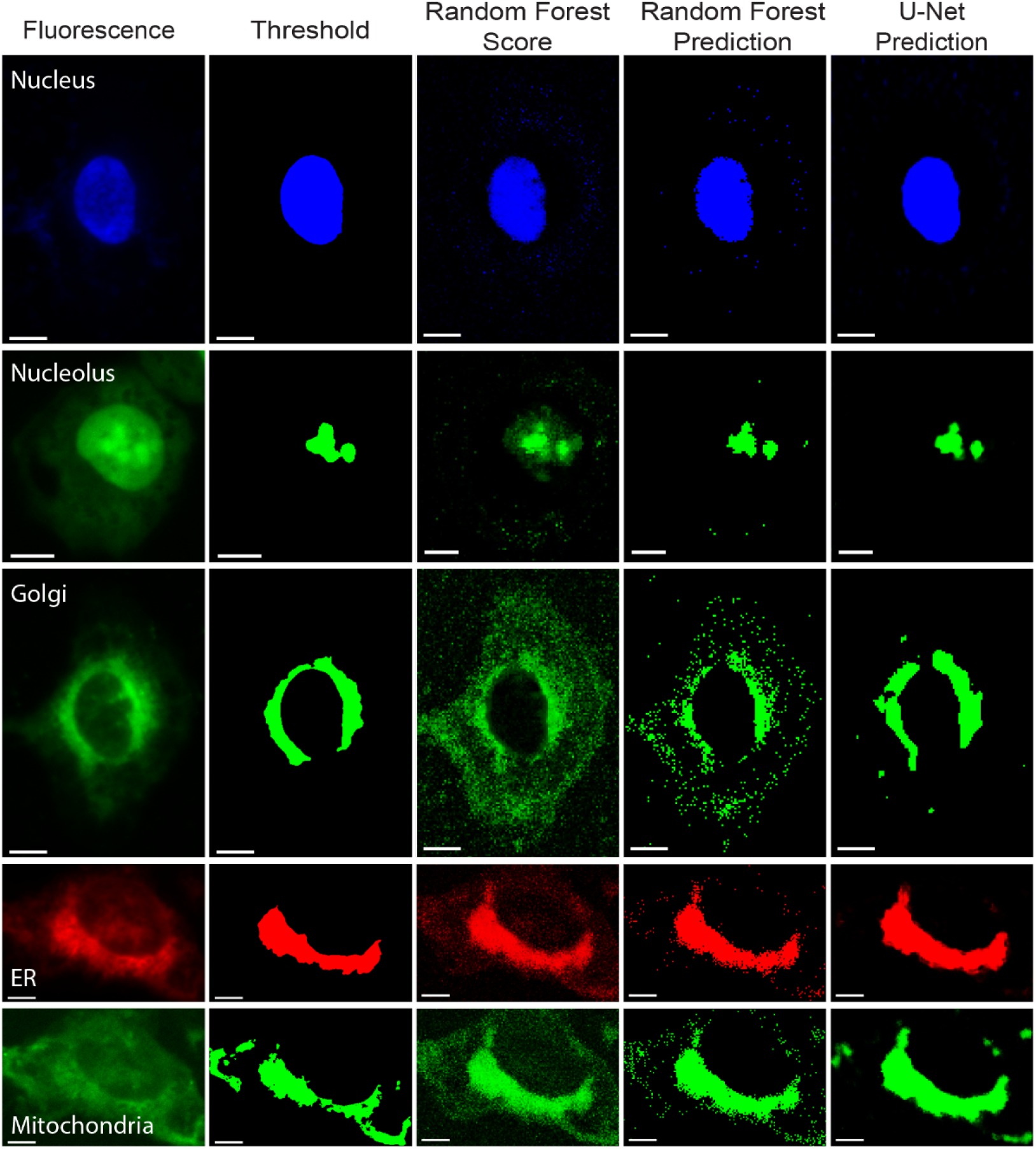
Random forest and U-Net classification of organelles in full cells. Representative classification of organelles in the context of full cells with the random forest and U-Net machine learning approaches. The two leftmost columns display fluorescence images and the fluorescent threshold (scale bars: 10 μm) for each organelle specific staining: nucleus (blue), nucleolus (green), Golgi apparatus (green), ER (red), mitochondria (green). The third column shows the random forest confidence scores, which provide information on the certainty of the random forest model. The fourth column shows classification results generated by the random forest model. The final column presents predictions from the U-Net model, which was trained using a combination of random forest scores and IR features. Training dataset sizes are provided in Table S2.

### Extension to Other Systems

To assess the generalizability of our classification method, we applied the RF and U-Net models trained on U-2 OS cells to hyperspectral data collected from HEK 293 cells. Despite biological and lineage differences between the bone cancer cell line (U-2 OS) and the immortalized kidney cell line (HEK 293), the models successfully identified the nucleus, nucleolus, and mitochondria in HEK 293 cells with average accuracies of 96.6%, 91.2%, and 88.9%, respectively (n=3) comparable to performance on U-2 OS cells (Fig. 5A-B). These results indicate that the models capture the characteristic biomolecular components of organelles rather than features specific to a particular cell line. This cross-cell line performance suggests that with appropriate training, the models could be broadly applicable for organelle classification across diverse cell types. Adaptability is a strength of the method, particularly for biomedical applications where deviations in biomolecular composition could be detectable through reduced model performance, such as those associated with disease or drug response. For example, a model trained on healthy cells could potentially detect health deviations based on systematic classification errors.

**Figure 5.**
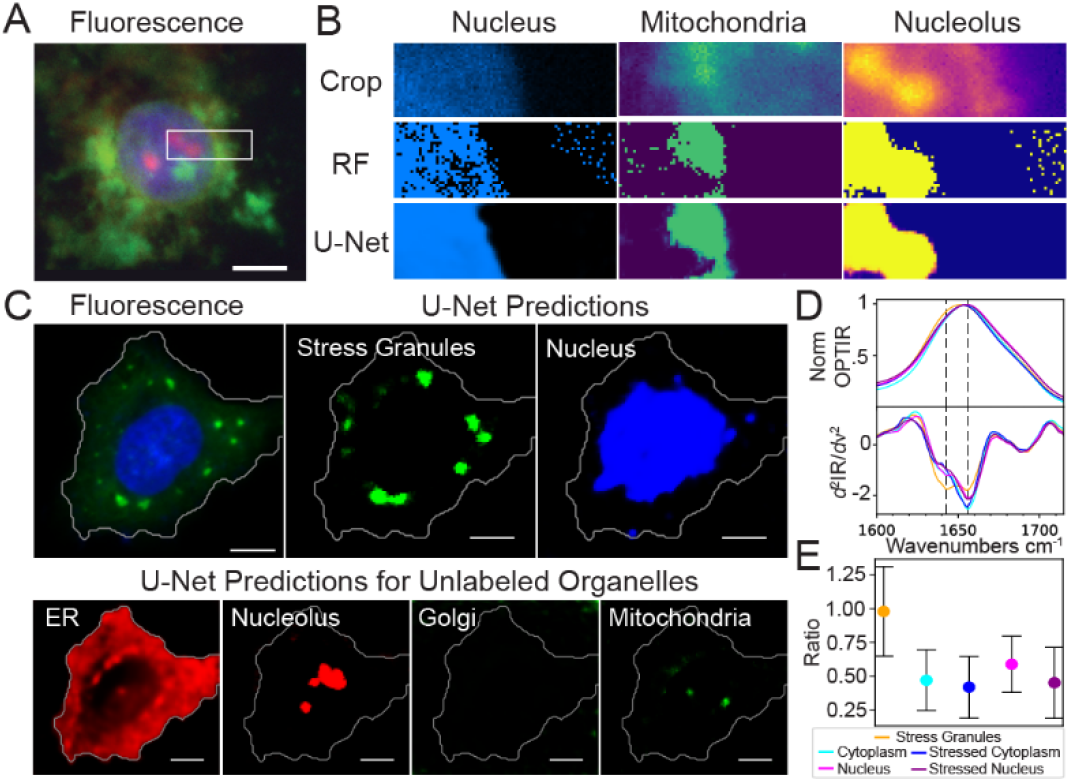
Classification of HEK 293 cells and stress granules. **A**. Representative fluorescence image of a HEK 293 cell with fluorescent labeling of the nucleus (blue), nucleolus (red), and mitochondria (green) (scale bar = 10 μm). The region of interest (ROI) is outlined in white. **B**. Classification of the ROI within HEK 293 cells using random forest (RF) and U-Net models trained on U-2 OS cells. The top panel shows a magnified fluorescent ROI, followed by random forest and U-Net classification results for the nucleus (left), mitochondria (middle), and nucleolus (right). **C**. Representative fluorescence image of stress granules (green) induced by oxidative stress within the cell (scale bar = 10 μm) with cellular region outlined in white. U-Net prediction images for each organelle are shown, giving poor predictions for all organelles except the nucleolus. **D**. Normalized OPTIR spectra (top) and second derivative (bottom) for stress granules and both the stressed and healthy cytoplasm and nucleus. Changes in protein secondary structure for beta sheet at 1643 cm^-1^ and alpha helices at 1656 cm^-1^ **E**. The ratio of the second derivative at 1643 cm^-1^ and 1656 cm^-1^ is compared. Stressed organelles show a decrease in this ratio compared to healthy spectra, which is paired with a large increase compared to the healthy cytoplasm.

To test this hypothesis, we placed cells under oxidative stress and evaluated classification accuracy. This also tested the ability of our workflow to classify transient biological phenomena formed under oxidative stress, stress granules. Despite being extensively studied, the structure, formation, and health implications of stress granules remain poorly understood. Spectral information on the cellular state after stress granule induction could provide valuable insights into their formation and function.^57^ Given that the models successfully identified another membraneless organelle, the nucleolus, we hypothesized that they could also be trained to identify transient membraneless compartments. Using the same workflow, we developed models that identify stress granules (Fig. 5C). Interestingly, despite their high accuracy for stress granules, our models for other organelles (besides the nucleolus) fail to generate reasonable predictions in stressed cells (Fig. 5C). Notably, the most prominent spectral difference in the amide I region between stress granules and the surrounding nucleus and cytoplasm is a peak at 1643 cm^-1^ (Fig. 5D), likely arising from beta-sheet rich RNA-binding domains of RNA-binding proteins densely packed within the granules.^58^ This is further highlighted in the ratio of the second derivative of the beta sheet (1643 cm^-1^) to the alpha helical (1656 cm^-1^) band of the different compartments (Fig. 5E). Spectral similarity between the cytoplasm and nucleus increased, suggesting that a breakdown in cellular organization leads to failure of models trained on healthy cell organelles.^59^ This opens new avenues for investigating the role of biomolecular condensation in cellular stress responses and disease pathogenesis.

## Discussion

In this study, we introduced a label-free, IR-based imaging workflow integrated with machine learning for high-resolution classification of subcellular structures, including the ER, Golgi apparatus, mitochondria, nucleus, nucleolus, and stress granules. With our two models, the RF and U-Net, we achieved accurate and robust classification across organelles of multiple cell types to support data collection in both single spectra and hyperspectral contexts. Key spectral features were selected based on distinct biochemical signatures, enabling precise organelle identification without fluorescent labels. The successful application of models trained on U-2 OS cells to HEK 293 cells demonstrates the generalizability of the approach to other cell lines. Together, these results position IR spectral imaging as a powerful, adaptable tool for quantitative, label-free organelle mapping and the investigation of cellular organization.

All microscopes require trade-offs among data quality, acquisition speed, ease of use, and cost. The optimal platform therefore depends on the biological question. In fluorescence microscopy, for example, widefield instruments are favored for applications that require speed, such as screening and live-cell imaging, whereas confocal microscopes provide improved spatial resolution for resolving fine structures. Fluorescent stains guide image acquisition and distinguish specific structures across these length scales. Analogously, biomolecular vibrational resonances provide intrinsic chemical contrasts for IR microscopy. Here, our models exploit the distinct chemical compositions of subcellular compartments to define vibrational ‘stains’ for organelle identification in IR images. This strategy should be generalizable to IR imaging at other spatial scales. Indeed, models trained on images acquired at 0.2 µm resolution successfully classified cell images collected at 0.5 µm resolution (Figure S27-29). As IR microscopy continues to advance across multiple length scales and acquisition speeds, such vibrational ‘stains’ provide a complementary framework for extracting biological information from this rapidly developing technology.

Additionally, accurate organelle classification does not require the full set of training parameters. To assess how parameter reduction affects performance, we systematically reduced the number of input parameters using the current training dataset (Fig. S12). Using the RF models, we iteratively removed the least important parameter for each organelle to establish a hierarchy of parameter importance and determine the minimum number of parameters needed for organelle detection. Nuclear classification was the most resilient to parameter reduction. With a single IR parameter, nuclei in half of the cells were classified with only a 12% decrease in accuracy relative to models trained with all seven IR parameters (Fig. S12A). By contrast, classification of most other organelles converged to approximately 50% accuracy with one IR parameter, consistent with near-random classification. Nevertheless, RF models for the Golgi apparatus and nucleolus also showed substantial resilience to parameter reduction, suggesting that comparable performance may be achievable with fewer spectral inputs (Fig. S12B).

We next tested how many parameters could be removed from U-Net models, using the nucleus and endoplasmic reticulum (ER) as representative examples (Figs. S13 and S14). Following the parameter-importance hierarchy defined in Fig. S12, U-Net models were trained with progressively fewer RF parameters until only a single parameter remained. As observed for the RF model, accurate U-Net prediction of nuclei was possible with one parameter: the full width at half maximum of the amide I band (Fig. S13C). The ER classification further highlighted the advantage of the U-Net architecture. Unlike the RF model, whose accuracy decreased steadily as parameters were removed, the U-Net model showed no loss in ER classification accuracy until only two parameters remained (Fig. S14B).

This analysis highlights the most critical parameters for classification that may be exploited in a workflow adapted for live-cell imaging. However, several technical challenges remain to be addressed. First, imaging times must be drastically reduced to maintain cell viability. As demonstrated above, this can be achieved by reducing the number of parameters. Additionally, recent advancements in OPTIR instrumentation have made video-rate imaging feasible through both wide-field and laser scanning methods.^60^ Second, high background absorption of water in the amide I region complicates the interpretation of protein-rich features in live cells. Although OPTIR suppresses the water background relative to traditional IR microscopy, spectral overlap remains significant. Emerging water-subtraction algorithms may help isolate biologically relevant features and enable clearer detection of subcellular structures in the hydrated environments of live cells.^61,62^ Interestingly, one of the most informative spectral regions for organelle classification was 1000-1145 cm^-1^ (Fig. S11A), which is distant from the prominent water band that overlaps with the amide I region. Furthermore, removing amide I parameters from the RF training dataset of the nucleus caused only an 8.8% drop in mean accuracy (Fig. S15), supporting the feasibility of water-tolerant classification schemes.

A key challenge of our approach lies in the reliance on fluorescence-based segmentation to generate training data and validation labels. This dependency is reflected in the discrepancy between the RF prediction accuracy for the training dataset (Table 1) and that of the hyperspectral images (Table 2). Fluorescent markers may not consistently label all regions of the target organelle and can exhibit non-specific binding, resulting in incomplete or ambiguous reference masks. Additionally, small changes in threshold selection can alter the binarization boundary, introducing bias in the validation sets. The edge of where an organelle ends and begins is often ambiguous across the cell. To ensure high-confidence labeling for training, the training dataset is composed of clear examples from each organelle class. While this strategy enhances labeling accuracy, it also makes the training dataset easier to classify, potentially overestimating the performance of the RF model. In contrast, when applied to real hyperspectral images, the model must classify spectra from boundary regions with lower labeling confidence, leading to reduced accuracy. Developing datasets that simultaneously label multiple organelles and train on different combinations of the labels could improve accuracy in regions where organelles overlap. However, the use of fluorescent labels inherently limits the total number of organelles that can be labeled simultaneously.

Promisingly, the U-Net model proved valuable for assessing organelle health and function. Under oxidative stress, not only did stress granules form in the cytosol (Fig. 5C-E) with a unique chemical composition, but also the chemical composition of other organelles changed. The spectral profile of the nucleus of stressed cells is distinct from the nucleus in healthy training sets (Fig. S16). This indicates that the classification models not only identify organelles but also specifically recognize healthy ones. In the future, models could be trained to identify the organelle state(s) of interest. Importantly, these compositional changes are undetectable using traditional immunofluorescent stains, which do not discriminate between healthy and diseased states. In contrast, IR spectra reveal compartment-specific composition changes. This capability offers a compelling foundation for developing diagnostic tools that operate at the subcellular level, enabling early detection of disease or stress responses via biochemical fingerprinting.

## Materials and Methods

### Cell Culture Methods

U-2 OS cells (ATCC HTB-96) were cultured in DMEM containing L-Glutamine, 4.5 g/L glucose, and sodium pyruvate (Corning 10013CM), supplemented with penicillin-streptomycin (ThermoFisher Scientific 15140122) at 100 U/mL and 10% fetal bovine serum (Gibco™ 10438026). Cell lines were maintained by splitting every three days at 80% confluency using trypsin-EDTA (Gibco™ 25300054). For imaging, U-2 OS cells between passages 5 and 30 were grown overnight on glass or calcium fluoride coverslips.

### Stress Granule Induction

Stress granules were induced by treating U-2 OS cells with DMEM (as described above) supplemented with 500 μM sodium arsenite (Sigma Aldrich S7400) and incubated for 40 minutes, before fixation with 4% PFA for 10 minutes at room temperature.^63^

### Dye Staining

Cells were fixed in 4% paraformaldehyde and stained according to the manufacturer’s instructions. The dyes used included: Golgi ID Green Assay Kit (Enzo Life Science ENZ-51028), ER-ID Red Assay Kit (Enzo Life Science ENZ-51026), Hoechst 33342 (Enzo Life Science from ENZ-51026), Nucleolus Bright Red (Dojindo N512), MitoView Green (Biotium 70054). After staining, cells were washed three times with PBS and three times with deionized water before imaging to prevent salt crystals.

### Immunofluorescence Staining

Cells were fixed in 4% paraformaldehyde for 10 minutes, permeabilized with 0.5% Tween-20 PBS for 15 minutes, and blocked with 1% BSA, 0.1% Tween-20 PBS solution for 45 minutes, with three PBS washes between each step. Primary antibody incubation was performed overnight at 4°C. Stress granules were stained with a rabbit polyclonal G3BP1 Antibody (Invitrogen PA5-29455) at a 1:100 dilution. Nuclear speckles were stained with a rabbit polyclonal SRRM2 antibody (Invitrogen #PA5-66827) at a 1:200 dilution. Secondary antibody incubation for both was with donkey anti-rabbit IgG Alexa Fluor™ Plus 488 (Invitrogen #A32790) at a 1:2000 dilution for 45 minutes. Cells were washed three times with PBS and three times with deionized water before imaging to prevent salt crystals.

### OPTIR Imaging

IR spectra were acquired using a mIRage-LS IR microscope (Photothermal Spectroscopy Corporation, Santa Barbara, CA) equipped with a four-module-pulsed quantum cascade laser (QCL) system (Daylight Solutions, San Diego, CA). All spectra were collected from 932 cm^-1^ to 1816 cm^-1^ in reflective mode with 2 cm^-1^ spectral resolution, averaging three scans per sample. The IR laser operated at 20% power with a 100 kHz pulse rate, while the probe laser (532 nm) operated at 11% power with a 2x or 5x gain. Scans were conducted at a speed of 200 cm^-1^/s with hyperspectral images captured at 0.2 μm resolution for cell sections and 0.5 μm resolution for whole cells. Predictive models also demonstrated accuracy at both resolutions. Imaging utilized a 40X, 0.78 NA reflective Schwarzschild objective (PIKE Technologies, Fitchburg, WI) and a 50X MPlanFl, 0.80 NA objective lens (Olympus, Japan). OPTIR imaging used the 40x objective, while fluorescence images were acquired using both objectives. This provides our OPTIR images with a spatial resolution of around 350 nm and an axial resolution of around 1 μm.^64^

### OPTIR Data Preparation

Hyperspectral datasets were exported from PTIR Studio (Photothermal Spectroscopy Corporation, Santa Barbara, CA) and analyzed in Python (Python 3.10.9). Training datasets were labeled by cross-referencing the fluorescent pixel positions to the corresponding IR spectra locations. Hyperspectral datasets were segmented using OTSU thresholding based on fluorescent labeling to compare with model predictions.

To account for cell-to-cell variability, IR Spectra were linearly background-subtracted using average signals from low-noise regions (978 – 988 cm^-1^ and 1770 to 1780 cm^-1^). Spectra within the 978 cm^-1^ to 1780 cm^-1^ range were normalized to the amide I signal. Second-derivative spectra were calculated using a seven-point (14 cm^-1^) Savitzky–Golay filter, followed by curve fitting with SciPy’s UnivariateSpline function.^19^ Training set sizes for each organelle are provided in Table S3.

### Random Forest Model

A random forest model was implemented using the scikit-learn Python package, comprising 100 decision trees.^65^ For accuracy assessments shown in Fig. S6, the model was trained on a subset of labeled data and validated on the remaining data. To generate the random forest scores utilized in the U-Net model and to predict hyperspectral maps, the model was trained on the entire dataset. To evaluate potential overfitting, the performance of the model for the test set was compared against its training dataset.

### U-Net Model

Using the Keras Python package, our U-Net model used a fairly standard U-Net architecture (for full layer list see SI materials and method).^66^ Training was performed using a batch size of 64 images over 50 epochs, with 10 steps per epoch. Model performance was monitored after every 10 epochs to ensure convergence and to detect potential trapping in local minima. Training was halted if a model did not meet predefined accuracy thresholds (60% for nucleus classification and 70% for other organelles), designed to test for prediction improvement over random assignment. Backpropagation was performed using the Adaptive Moment Estimation (Adam) optimizer and evaluated with a binary cross-entropy loss function. Additional details can be found in the SI on Page S2-S7.

### Model Evaluation

The models were evaluated against several key performance metrics (Page S8). All metrics are evaluated within [0,1], and a higher value means better prediction. The metrics are presented alongside the predictions in Figure S17-S29.

## Author Contributions

C.M.D. conceived of the idea. C.M.D. and V.S.B. obtained funding and supervised the experiments and computation. M.J.B. performed the experiments, computations, and training. All authors discussed the results and contributed to the final manuscript.

## Acknowledgments

This work was supported by a Beckman Young Investigator Award from the Arnold and Mabel Beckman Foundation (https://dx.doi.org/10.13039/100000997).

## References

(1) Cuanalo-Contreras, K.; Schulz, J.; Mukherjee, A.; Park, K.-W.; Armijo, E.; Soto, C. Extensive Accumulation of Misfolded Protein Aggregates during Natural Aging and Senescence. Front Aging Neurosci 2023, 14, 1090109. 10.3389/fnagi.2022.1090109.

(2) Agmon, E.; Stockwell, B. R. Lipid Homeostasis and Regulated Cell Death. Current Opinion in Chemical Biology 2017, 39, 83–89. 10.1016/j.cbpa.2017.06.002.

(3) Baker, S. A.; Rutter, J. Metabolites as Signalling Molecules. Nat Rev Mol Cell Biol 2023, 24 (5), 355–374. 10.1038/s41580-022-00572-w.

(4) Casey, J. R.; Grinstein, S.; Orlowski, J. Sensors and Regulators of Intracellular pH. Nat Rev Mol Cell Biol 2010, 11 (1), 50–61. 10.1038/nrm2820.

(5) Day, R. N.; Davidson, M. W. The Fluorescent Protein Palette: Tools for Cellular Imaging. Chem. Soc. Rev. 2009, 38 (10), 2887. 10.1039/b901966a.

(6) Ozcan, L.; Tabas, I. Role of Endoplasmic Reticulum Stress in Metabolic Disease and Other Disorders. Annu Rev Med 2012, 63, 317–328. 10.1146/annurev-med-043010-144749.

(7) Katayama, T.; Imaizumi, K.; Sato, N.; Miyoshi, K.; Kudo, T.; Hitomi, J.; Morihara, T.; Yoneda, T.; Gomi, F.; Mori, Y.; Nakano, Y.; Takeda, J.; Tsuda, T.; Itoyama, Y.; Murayama, O.; Takashima, A.; St George-Hyslop, P.; Takeda, M.; Tohyama, M. Presenilin-1 Mutations Downregulate the Signalling Pathway of the Unfolded-Protein Response. Nat Cell Biol 1999, 1 (8), 479–485. 10.1038/70265.

(8) Laybutt, D. R.; Preston, A. M.; Akerfeldt, M. C.; Kench, J. G.; Busch, A. K.; Biankin, A. V.; Biden, T. J. Endoplasmic Reticulum Stress Contributes to Beta Cell Apoptosis in Type 2 Diabetes. Diabetologia 2007, 50 (4), 752–763. 10.1007/s00125-006-0590-z.

(9) Kammoun, H. L.; Chabanon, H.; Hainault, I.; Luquet, S.; Magnan, C.; Koike, T.; Ferré, P.; Foufelle, F. GRP78 Expression Inhibits Insulin and ER Stress-Induced SREBP-1c Activation and Reduces Hepatic Steatosis in Mice. J Clin Invest 2009, 119 (5), 1201–1215. 10.1172/JCI37007.

(10) Roussel, B. D.; Kruppa, A. J.; Miranda, E.; Crowther, D. C.; Lomas, D. A.; Marciniak, S. J. Endoplasmic Reticulum Dysfunction in Neurological Disease. Lancet Neurol 2013, 12 (1), 105–118. 10.1016/S1474-4422(12)70238-7.

(11) Scorrano, L.; De Matteis, M. A.; Emr, S.; Giordano, F.; Hajnóczky, G.; Kornmann, B.; Lackner, L. L.; Levine, T. P.; Pellegrini, L.; Reinisch, K.; Rizzuto, R.; Simmen, T.; Stenmark, H.; Ungermann, C.; Schuldiner, M. Coming Together to Define Membrane Contact Sites. Nat Commun 2019, 10 (1), 1287. 10.1038/s41467-019-09253-3.

(12) Audano, M.; Schneider, A.; Mitro, N. Mitochondria, Lysosomes, and Dysfunction: Their Meaning in Neurodegeneration. Journal of Neurochemistry 2018, 147 (3), 291–309. 10.1111/jnc.14471.

(13) Wong, Y. C.; Kim, S.; Peng, W.; Krainc, D. Regulation and Function of Mitochondria–Lysosome Membrane Contact Sites in Cellular Homeostasis. Trends in Cell Biology 2019, 29 (6), 500–513. 10.1016/j.tcb.2019.02.004.

(14) Kim, S.; Wong, Y. C.; Gao, F.; Krainc, D. Dysregulation of Mitochondria-Lysosome Contacts by GBA1 Dysfunction in Dopaminergic Neuronal Models of Parkinson’s Disease. Nat Commun 2021, 12 (1), 1807. 10.1038/s41467-021-22113-3.

(15) Grinstein, S.; Dixon, S. J. Ion Transport, Membrane Potential, and Cytoplasmic pH in Lymphocytes: Changes during Activation. Physiological Reviews 1989, 69 (2), 417–481. 10.1152/physrev.1989.69.2.417.

(16) Jones, D. P.; Go, Y.-M. Redox Compartmentalization and Cellular Stress. Diabetes Obesity Metabolism 2010, 12 (s2), 116–125. 10.1111/j.1463-1326.2010.01266.x.

(17) Matthäus, C.; Bird, B.; Miljković, M.; Chernenko, T.; Romeo, M.; Diem, M. Chapter 10 Infrared and Raman Microscopy in Cell Biology. In Methods in Cell Biology; Biophysical Tools for Biologists, Volume Two: In Vivo Techniques; Academic Press, 2008; Vol. 89, pp 275–308. 10.1016/S0091-679X(08)00610-9.

(18) Zhang, D.; Li, C.; Zhang, C.; Slipchenko, M. N.; Eakins, G.; Cheng, J.-X. Depth-Resolved Mid-Infrared Photothermal Imaging of Living Cells and Organisms with Submicrometer Spatial Resolution. Sci. Adv. 2016, 2 (9), e1600521. 10.1126/sciadv.1600521.

(19) Yang, H.; Yang, S.; Kong, J.; Dong, A.; Yu, S. Obtaining Information about Protein Secondary Structures in Aqueous Solution Using Fourier Transform IR Spectroscopy. Nat Protoc 2015, 10 (3), 382–396. 10.1038/nprot.2015.024.

(20) DeFlores, L. P.; Ganim, Z.; Nicodemus, R. A.; Tokmakoff, A. Amide I′−II′ 2D IR Spectroscopy Provides Enhanced Protein Secondary Structural Sensitivity. J. Am. Chem. Soc. 2009, 131 (9), 3385–3391. 10.1021/ja8094922.

(21) Barth, A.; Zscherp, C. What Vibrations Tell about Proteins. Quart. Rev. Biophys. 2002, 35 (4), 369–430. 10.1017/S0033583502003815.

(22) Ali, S. M.; Bonnier, F.; Lambkin, H.; Flynn, K.; McDonagh, V.; Healy, C.; Lee, T. C.; Lyng, F. M.; Byrne, H. J. A Comparison of Raman, FTIR and ATR-FTIR Micro Spectroscopy for Imaging Human Skin Tissue Sections. Anal. Methods 2013, 5 (9), 2281–2291. 10.1039/C3AY40185E.

(23) Movasaghi, Z.; Rehman, S.; Ur Rehman, Dr. I. Fourier Transform Infrared (FTIR) Spectroscopy of Biological Tissues. Applied Spectroscopy Reviews 2008, 43 (2), 134–179. 10.1080/05704920701829043.

(24) Talari, A.; Garcia Martinez, M.; Movasaghi, Z.; Rehman, S.; Rehman, I. Advances in Fourier Transform Infrared (FTIR) Spectroscopy of Biological Tissues. Applied Spectroscopy Reviews 2016, 52, 00–00. 10.1080/05704928.2016.1230863.

(25) Baliyan, A.; Imai, H. Machine Learning Based Analytical Framework for Automatic Hyperspectral Raman Analysis of Lithium-Ion Battery Electrodes. Sci Rep 2019, 9 (1), 18241. 10.1038/s41598-019-54770-2.

(26) Klein, K.; Gigler, A. M.; Aschenbrenner, T.; Monetti, R.; Bunk, W.; Jamitzky, F.; Morfill, G.; Stark, R. W.; Schlegel, J. Label-Free Live-Cell Imaging with Confocal Raman Microscopy. Biophysical Journal 2012, 102 (2), 360–368. 10.1016/j.bpj.2011.12.027.

(27) Li, Y.; Heo, J.; Lim, C.-K.; Pliss, A.; Kachynski, A. V.; Kuzmin, A. N.; Kim, S.; Prasad, P. N. Organelle Specific Imaging in Live Cells and Immuno-Labeling Using Resonance Raman Probe. Biomaterials 2015, 53, 25–31. 10.1016/j.biomaterials.2015.02.056.

(28) Kuhar, N.; Sil, S.; Verma, T.; Umapathy, S. Challenges in Application of Raman Spectroscopy to Biology and Materials. RSC Adv. 2018, 8 (46), 25888–25908. 10.1039/C8RA04491K.

(29) Wrobel, T. P.; Bhargava, R. Infrared Spectroscopic Imaging Advances as an Analytical Technology for Biomedical Sciences. Anal. Chem. 2018, 90 (3), 1444–1463. 10.1021/acs.analchem.7b05330.

(30) Greaves, G. E.; Kiryushko, D.; Auner, H. W.; Porter, A. E.; Phillips, C. C. Label-Free Nanoscale Mapping of Intracellular Organelle Chemistry. Commun Biol 2023, 6 (1), 583. 10.1038/s42003-023-04943-7.

(31) Prater, C. B.; Kjoller, K. J.; Stuart, A. P. D.; Grigg, D. A.; ‘Limurn, R.; Gough, K. M. Widefield Super-Resolution Infrared Spectroscopy and Imaging of Autofluorescent Biological Materials and Photosynthetic Microorganisms Using Fluorescence Detected Photothermal Infrared (FL-PTIR). Appl Spectrosc 2024, 78 (11), 1208–1219. 10.1177/00037028241256978.

(32) Gajjela, C. C.; Brun, M.; Mankar, R.; Corvigno, S.; Kennedy, N.; Zhong, Y.; Liu, J.; Sood, A. K.; Mayerich, D.; Berisha, S.; Reddy, R. Leveraging Mid-Infrared Spectroscopic Imaging and Deep Learning for Tissue Subtype Classification in Ovarian Cancer. Analyst 2023, 148 (12), 2699–2708. 10.1039/D2AN01035F.

(33) Reihanisaransari, R.; Gajjela, C. C.; Wu, X.; Ishrak, R.; Zhong, Y.; Mayerich, D.; Berisha, S.; Reddy, R. Cervical Cancer Tissue Analysis Using Photothermal Midinfrared Spectroscopic Imaging. Chemical & Biomedical Imaging 2024, 2 (9), 651–658. 10.1021/cbmi.4c00031.

(34) Weeks, S. E.; Metge, B. J.; Samant, R. S. The Nucleolus: A Central Response Hub for the Stressors That Drive Cancer Progression. Cell. Mol. Life Sci. 2019, 76 (22), 4511–4524. 10.1007/s00018-019-03231-0.

(35) Galganski, L.; Urbanek, M. O.; Krzyzosiak, W. J. Nuclear Speckles: Molecular Organization, Biological Function and Role in Disease. Nucleic Acids Research 2017, 45 (18), 10350–10368. 10.1093/nar/gkx759.

(36) Mohan, A. G.; Calenic, B.; Ghiurau, N. A.; Duncea-Borca, R.-M.; Constantinescu, A.-E.; Constantinescu, I. The Golgi Apparatus: A Voyage through Time, Structure, Function and Implication in Neurodegenerative Disorders. Cells 2023, 12 (15), 1972. 10.3390/cells12151972.

(37) Liu, J.; Huang, Y.; Li, T.; Jiang, Z.; Zeng, L.; Hu, Z. The Role of the Golgi Apparatus in Disease (Review). International Journal of Molecular Medicine 2021, 47 (4), 1–1. 10.3892/ijmm.2021.4871.

(38) Whelan, D. R.; Bell, T. D. M. Correlative Synchrotron Fourier Transform Infrared Spectroscopy and Single Molecule Super Resolution Microscopy for the Detection of Composition and Ultrastructure Alterations in Single Cells. ACS Chem. Biol. 2015, 10 (12), 2874–2883. 10.1021/acschembio.5b00754.

(39) Petersen, T. W.; Ibrahim, S. F.; Diercks, A. H.; Van Den Engh, G. Chromatic Shifts in the Fluorescence Emitted by Murine Thymocytes Stained with Hoechst 33342. Cytometry Pt A 2004, 60A (2), 173–181. 10.1002/cyto.a.20058.

(40) Chandel, N. S. Carbohydrate Metabolism. Cold Spring Harb Perspect Biol 2021, 13 (1), a040568. 10.1101/cshperspect.a040568.

(41) Alberts, B.; Johnson, A.; Lewis, J. The Structure and Function of DNA. In Molecular Biology of the Cell; Garland Science: New York, 2000.

(42) Coblentz Society; Stein, S. E. D-Glucose Anhydrous in Evaluated Infrared Reference Spectra. In NIST Chemistry WebBook, NIST Standard Reference Database 69; National Institute of Standards and Technology; Vol. 20899.

(43) Zetterberg, A. Nuclear and Cytoplasmic Nucleic Acid Content and Cytoplasmic Protein Synthesis during Interphase in Mouse Fibroblasts in Vitro. Experimental Cell Research 1966, 43 (3), 517–525. 10.1016/0014-4827(66)90022-X.

(44) Banyay, M.; Sarkar, M.; Gräslund, A. A Library of IR Bands of Nucleic Acids in Solution. Biophysical Chemistry 2003, 104 (2), 477–488. 10.1016/S0301-4622(03)00035-8.

(45) Taillandier, E.; Peticolas, W. L.; Adam, S.; Huynh-Dinh, T.; Igolen, J. Polymorphism of the d(CCCGCGGG)2 Double Helix Studied by FT-i.r. Spectroscopy. Spectrochimica Acta Part A: Molecular Spectroscopy 1990, 46 (1), 107–112. 10.1016/0584-8539(93)80018-6.

(46) Ledeen, R.; Wu, G. Nuclear Lipids and Their Metabolic and Signaling Properties. In Handbook of Neurochemistry and Molecular Neurobiology; Lajtha, A., Tettamanti, G., Goracci, G., Eds.; Springer US: Boston, MA, 2009; pp 173–198. 10.1007/978-0-387-30378-9_7.

(47) Walther, T. C.; Farese, R. V. Lipid Droplets and Cellular Lipid Metabolism. Annu Rev Biochem 2012, 81, 687–714. 10.1146/annurev-biochem-061009-102430.

(48) Derenne, A.; Claessens, T.; Conus, C.; Goormaghtigh, E. Infrared Spectroscopy of Membrane Lipids. In Encyclopedia of Biophysics; Roberts, G. C. K., Ed.; Springer Berlin Heidelberg: Berlin, Heidelberg, 2013; pp 1074–1081. 10.1007/978-3-642-16712-6_558.

(49) Liu, Y.; Lunter, D. J. Selective and Sensitive Spectral Signals on Confocal Raman Spectroscopy for Detection of Ex Vivo Skin Lipid Properties. Transl Biophotonics 2020, 2 (3), e202000003. 10.1002/tbio.202000003.

(50) Rohman, A.; Windarsih, A.; Lukitaningsih, E.; Rafi, M.; Betania, K.; Fadzillah, N. A. The Use of FTIR and Raman Spectroscopy in Combination with Chemometrics for Analysis of Biomolecules in Biomedical Fluids: A Review. BSI 2020, 8 (3–4), 55–71. 10.3233/BSI-200189.

(51) Dovbeshko, G. FTIR Spectroscopy Studies of Nucleic Acid Damage. Talanta 2000, 53 (1), 233–246. 10.1016/S0039-9140(00)00462-8.

(52) Chiriboga, L.; Xie, P.; Yee, H.; Vigorita, V.; Zarou, D.; Zakim, D.; Diem, M. Infrared Spectroscopy of Human Tissue. I. Differentiation and Maturation of Epithelial Cells in the Human Cervix. Biospectroscopy 1998, 4 (1), 47–53. 10.1002/(SICI)1520-6343(1998)4:1%253C47::AID-BSPY5%253E3.0.CO;2-P.

(53) Max, J.-J.; Chapados, C. Infrared Spectroscopy of Aqueous Carboxylic Acids: Comparison between Different Acids and Their Salts. J. Phys. Chem. A 2004, 108 (16), 3324–3337. 10.1021/jp036401t.

(54) Nowakowska, A. M.; Pieczara, A.; Korona, W.; Borek-Dorosz, A.; Adamczyk, A.; Dawiec, P.; Leszczenko, P.; Orleanska, J.; Machalska, E.; Orzechowska, B.; Orzechowska, S.; Brzozowski, K.; Krynicka, W.; Majzner, K.; Malek, K.; Baranska, M. The Raman Map of the Human Cell. Anal. Chem. 2025, 97 (30), 16374–16382. 10.1021/acs.analchem.5c02035.

(55) Breiman, L. Random Forests. Machine Learning 2001, 45 (1), 5–32. 10.1023/A:1010933404324.

(56) Al-Mualem, Z. A.; Baiz, C. R. Generative Adversarial Neural Networks for Denoising Coherent Multidimensional Spectra. J. Phys. Chem. A 2022, 126 (23), 3816–3825. 10.1021/acs.jpca.2c02605.

(57) Millar, S. R.; Huang, J. Q.; Schreiber, K. J.; Tsai, Y.-C.; Won, J.; Zhang, J.; Moses, A. M.; Youn, J.-Y. A New Phase of Networking: The Molecular Composition and Regulatory Dynamics of Mammalian Stress Granules. Chem. Rev. 2023, 123 (14), 9036–9064. 10.1021/acs.chemrev.2c00608.

(58) Besnard-Guérin, C. Cytoplasmic Localization of Amyotrophic Lateral Sclerosis-related TDP-43 Proteins Modulates Stress Granule Formation. Eur J of Neuroscience 2020, 52 (8), 3995–4008. 10.1111/ejn.14762.

(59) Zhang, K.; Daigle, J. G.; Cunningham, K. M.; Coyne, A. N.; Ruan, K.; Grima, J. C.; Bowen, K. E.; Wadhwa, H.; Yang, P.; Rigo, F.; Taylor, J. P.; Gitler, A. D.; Rothstein, J. D.; Lloyd, T. E. Stress Granule Assembly Disrupts Nucleocytoplasmic Transport. Cell 2018, 173 (4), 958–971.e17. 10.1016/j.cell.2018.03.025.

(60) Adak, S.; Raveendran, A. T.; Popp, J.; Krafft, C. High Resolution and Fast Chemical Imaging Using Widefield Optical Photothermal Infrared Microscope. In Biomedical Spectroscopy, Microscopy, and Imaging III; Popp, J., Gergely, C., Eds.; SPIE: Strasbourg, France, 2024; p 78. 10.1117/12.3016833.

(61) Trevisan, J.; Angelov, P. P.; Carmichael, P. L.; Scott, A. D.; Martin, F. L. Extracting Biological Information with Computational Analysis of Fourier-Transform Infrared (FTIR) Biospectroscopy Datasets: Current Practices to Future Perspectives. Analyst 2012, 137 (14), 3202. 10.1039/c2an16300d.

(62) Zhao, J.; Cui, J.-K.; Chen, R.-X.; Tang, Z.-Z.; Tan, Z.-L.; Jiang, L.-Y.; Liu, F. Real-Time in-Situ Quantification of Protein Secondary Structures in Aqueous Solution Based on ATR-FTIR Subtraction Spectrum. Biochemical Engineering Journal 2021, 176, 108225. 10.1016/j.bej.2021.108225.

(63) Lee, A. K.; Potts, P. R. Protocol for the Determination of Intracellular Phase Separation Thresholds. STAR Protocols 2021, 2 (1), 100308. 10.1016/j.xpro.2021.100308.

(64) Prater, C. B.; Kansiz, M.; Cheng, J.-X. A Tutorial on Optical Photothermal Infrared (O-PTIR) Microscopy. APL Photonics 2024, 9 (9), 091101. 10.1063/5.0219983.

(65) Pedregosa, F.; Varoquaux, G.; Gramfort, A.; Michel, V.; Thirion, B.; Grisel, O.; Blondel, M.; Prettenhofer, P.; Weiss, R.; Dubourg, V.; Vanderplas, J.; Passos, A.; Cournapeau, D.; Brucher, M.; Perrot, M.; Duchesnay, É. Scikit-Learn: Machine Learning in Python. Journal of Machine Learning Research 2011, 12 (85), 2825–2830.

(66) Ronneberger, O.; Fischer, P.; Brox, T. U-Net: Convolutional Networks for Biomedical Image Segmentation. In Medical Image Computing and Computer-Assisted Intervention – MICCAI 2015; Navab, N., Hornegger, J., Wells, W. M., Frangi, A. F., Eds.; Springer International Publishing: Cham, 2015; pp 234–241. 10.1007/978-3-319-24574-4_28.

