## Supplementary Information for "Label-Free AI-Classification of Subcellular Organelles Based on Optical Photothermal Infrared Images"

This PDF file includes:

- Additional Materials and Methods
- Figures S1-S29
- Table S1-S2
- SI References

### Materials and Methods

#### Scheme S1. U-Net Model Training

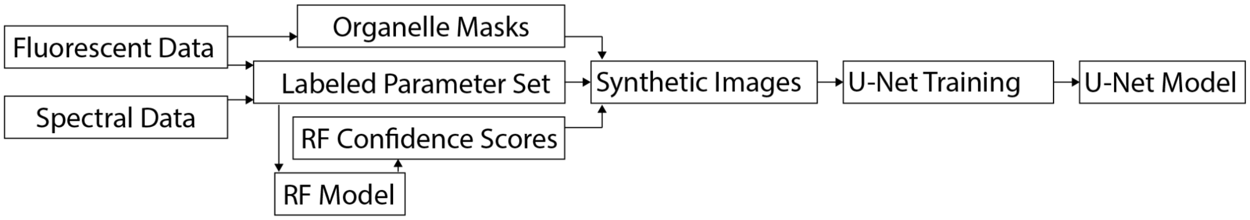


The section below provides details on how the U-Net and RF models were trained. Scheme S1 shows the workflow used to train each model. More details about each section of the scheme are provided below, with information on the collection of fluorescent and spectral data present in the main text.

#### Labeled Parameter Set

***Parameter Sources.*** The IR features relevant to each organelle are summarized in Table S1. Parameters for machine learning were derived from three types of data: 1. Single IR absorption values; 2. Comparisons between two IR absorption values; or 3. Values obtained through additional analytical processing.

- Parameters derived from a single IR absorption value
  - From OPTIR Signal: 2, 3, 6, 16
  - From Second Derivative: 1, 5, 9, 12, 13, 15
- Parameters derived from the comparison of two IR absorption values
  - From OPTIR Signal: 8 (Ratio)
  - From Second Derivative: 4 (Difference), 14 (Ratio)
- Parameters derived from additional analyses
  - From OPTIR Signal
    - 7: Peak location near 1390 $cm^{-1}$
    - 10: Ratio of the highest signal in the amide II region to the highest signal in the amide I region
    - 11: Full width at half maximum of the Amide I.

#### Organelle Masks for U-Net Training

U-Net training requires many example images to train the model to identify possible spatial configurations. However, random forest training demonstrated that high accuracies could be reached with a relatively low number of infrared spectra. From this we concluded that it was not necessary that the example images for U-Net training contain unique spectral data. Therefore, we created synthetic data sets by combining geometries from real fluorescent data with real spectral data. This section details how, before training the U-Net model, we generate a set of organelle masks from real fluorescent images.

Masking of the fluorescent data is tailored to each organelle to account for the signal-to-noise ratio of the stain and different organelle morphologies. In general, each method uses Otsu thresholding. Often because the background, cytoplasm, and nucleoplasm differ more than the organelles within them, Otsu thresholding is used twice. The first application of Otsu thresholding separates the compartment where the organelle is housed (cytoplasm or nucleus). The second application of Otsu thresholding is applied just to the compartment where the organelle resides. Because our goal is to generate a varied set of possible organelle morphologies and stain efficiencies, we further adjust the masks with either erosion or dilation steps. These parameters are adjusted for the different organelle to balance increasing variation with the retention of reasonable organelle morphologies.

*Cytoplasmic Organelle Mask Generation (Golgi, ER, and Mitochondria)*

A two-step thresholding was used to capture both a loose and a strict threshold of the fluorescence. The broad threshold with Otsu thresholding over the entire cell and background. A more stringent mask was then applied to either the mask from step one or on the area just within the cell. To increase variability while addressing thresholding artifacts, the masks underwent some morphological processing. This was done through either iterative erosion or iterative dilation, with each iteration saved as a potential mask. In order to generate masks with reasonable organelle morphologies, the process was adjusted based off the initial organelle coverage. These masks were combined with a nucleus mask in order for the assignment of the appropriate spectral data.

- Golgi:
    - > 35%: Eroded three times, no dilation.
    - 35 – 30%: Eroded twice, dilated once.
    - 30 – 25%: Eroded once, dilated twice.
    - < 25%: No Erosion, dilated three times.
- ER and Mitochondria
    - > 50%: Eroded three times, no dilation.
    - 50 – 40%: Eroded twice, dilated once.
    - 40 – 35%: Eroded once, dilated twice.
    - < 25%: No Erosion, dilated three times.

*Nucleus Mask Generation*

To introduce variability in nuclear masks, Gaussian smoothing was applied with three different standard deviations (σ = 1, 3, 5), followed by binary thresholding and dilation to recover size variations. This process generated four versions of the nucleus: the initial threshold mask and three smoothed versions. The cytoplasm and nucleoplasm were distinct negative classes when training, so the objective was to increase variability at the interface between the organelle of interest and the nucleus.

Each of these nucleus variants was then combined with each organelle variant, resulting in 32 possible masks per cell. When the regions of overlap between the nucleus and the organelle were small, the overlapping areas were designated as nucleus. However, if the overlap was significant, an additional 32 masks were generated where the overlapping regions were assigned to the organelle.

*Nucleolus Mask Generation*

Nucleolus masks were generated using a single fluorescent threshold applied exclusively within the nuclear region. To increase variability, the initial mask threshold separately underwent five iterative erosions (σ = 0.5) or two iterative dilations (σ = 1), yielding eight distinct mask variants. Combining these eight nucleolus variants with four previously generated nucleus variants (calculated as specified previously) resulted in 32 total nucleolus masks per cell.

*Nuclear Speckles Mask Generation*

To create the nuclear speckles training mask set, we first independently thresholded the speckles, nucleolus, and nucleus. To introduce morphological diversity, each thresholded mask was processed using Gaussian smoothing dilations (σ = 1, 3, 5), generating three additional variants per structure for a total of four variants each. Combinations of these masks yielded 64 total masks per cell. This comprehensive approach was necessary because the nucleolus and nuclear speckles share several spectral parameters but can differ significantly from the surrounding nucleoplasm. Including both structures in the U-net training dataset ensures that the model learns to accurately distinguish between them.

*Stress Granule Mask Generation*

Stress granule masks were generated by first thresholding the granule-specific fluorescence signal. To further enhance variability, four dilation steps were applied, followed by three Gaussian smoothing conditions ($\sigma$ = 1, 2, 3). This process generated eight mask variants per cell. Each of these stress granule variants was then combined with four nucleus variants, producing 32 total masks per cell. This approach ensured that the training data captured the variability in stress granule morphology.

**Synthetic Images**

The goal of this process was to use existing fluorescent images and data to represent hyperspectral data. To do this we would create masks based on fluorescent value thresholds and populate it with real data. To effectively sample those masks and data distributions the following approach was used.

*1. Mask Selection:* A cell was first randomly selected from the organelle mask set. Each cell had either 32 or 64 masks generated from the fluorescent data using different thresholding conditions (described below). Due to the spectral differences of the cytoplasm and nucleoplasm, negative data class data had to be separated into negative(nucleoplasm) and negative(cytoplasm). To allow for this, the masks also had to be labeled with the region corresponding to the organelle of interest, the nucleoplasm, and cytoplasm. Each organelle required a slightly different approach for generating the mask set, which is described below.

*2. Mask Variation:* To increase the variation further the selected mask was rotated (full 360$^{o}$ rotation allowed) and independently scaled in both the width and height (±20%) to allow for skewing the image. After a 64x64 pixel region was selected from the cell image to populate with data for training. The location was randomly selected with a bias for images that contain regions with organelle of interest. This was done to ensure the model had enough data to find the organelle of interest, but also could show regions with no organelle of interest. To ensure this we included extra images at the end of each dataset that was entirely composed of each of the classes within the mask. For example, when training the Golgi, four additional masks would be provided. These images would be entirely calcium fluoride, cytoplasm without the Golgi, the Golgi, and the nucleoplasm.

*3. Data Population:* From the spectral training dataset, a random cell was selected. Before U-net training each cell has the distributions of values for each parameter for the organelle, cytoplasm, and nucleoplasm was calculated. For each pixel in the mask, this distribution was randomly sampled to populate the image with spectral feature data. Once the image was populated with these sampled values, each pixel was then passed through the previously trained random forest (RF) model. The model class probabilities than were used as a feature for the model. The features that did not come from the random forest were then normalized from 0 to 1. The random forest class probabilities were not normalized because they are already scaled from 0 to 1 and the preservation of these values is important for assigning class values in cases where the imaged area is homogenous.

*4. Batch Assembly*: The entire process (mask variation, data population, model scoring, and normalization) was repeated for each image in the batch. A new synthetic batch was generated for every training epoch, ensuring that the model was trained on new data throughout training. As previously mentioned, the final images in each batch were homogenous examples, generated using masks containing only calcium fluoride, nucleoplasm, cytoplasm, and organelle of interest.

**U-Net Layer List:**

- Input: (64 x 64 x Number of Features + 1 For Random Forest Scores)
- Encoding Arm – 3 Blocks:
    - Block 1
        - 2D Convolution, Kernel Size = 3, Activation = "Relu"
            - Output: 64 x 64 x 64
        - 2D Convolution, Kernel Size = 3, Activation = "Relu"
            - Output: 64 x 64 x 64
        - 2D Max Pooling, Pool Size = (2, 2)
            - Output: 32 x 32 x 64
    - Block 2
        - 2D Convolution, Kernel Size = 3, Activation = "Relu"
            - Output: 32 x 32 x 128
        - 2D Convolution, Kernel Size = 3, Activation = "Relu"
            - Output: 32 x 32 x 128
        - 2D Max Pooling, Pool Size = (2, 2)
            - Output: 16 x 16 x 128
    - Block 3
        - 2D Convolution, Kernel Size = 3, Activation = "Relu"
            - Output: 16 x 16 x 256
        - 2D Convolution, Kernel Size = 3, Activation = "Relu"
            - Output: 16 x 16 x 256
        - 2D Max Pooling, Pool Size = (2, 2)
            - Output: 8 x 8 x 256
        - Dropout Layer (Frequency = 0.1)
            - Output: 8 x 8 x 256
- Bottom
    - 2D Convolution, Kernel Size = 3, Activation = "Relu"
        - Output: 8 x 8 x 512
    - 2D Convolution, Kernel Size = 3, Activation = "Relu"
        - Output: 8 x 8 x 512
    - Dropout Layer (Frequency = 0.1)
        - Output: 8 x 8 x 512
- Decoding Arm
    - Block 1
        - 2D Up sampling Layer
            - Output: 16 x 16 x 512
        - 2D Convolution Layer
            - Output: 16 x 16 x 256
        - Concatenate with Encoding Block 3, 2D Convolution Layer
            - Output: 16 x 16 x 512
        - 2D Convolution, Kernel Size = 3, Activation = "Relu"
            - Output: 16 x 16 x 256
        - 2D Convolution, Kernel Size = 3, Activation = "Relu"
            - Output: 16 x 16 x 256
    - Block 2
        - 2D Up sampling Layer
            - Output: 32 x 32 x 256

                - 2D Convolution Layer
                    - Output: 32 x 32 x 128
                - Concatenate with Encoding Block 2, 2D Convolution Layer
                    - Output: 32 x 32 x 256
                - 2D Convolution, Kernel Size = 3, Activation = “Relu”
                    - Output: 32 x 32 x 128
                - 2D Convolution, Kernel Size = 3, Activation = “Relu”
                    - Output: 32 x 32 x 128
        - Block 3
            - 2D Up sampling Layer
                - Output: 64 x 64 x 128
            - 2D Convolution Layer
                - Output: 64 x 64 x 64
            - Concatenate with Encoding Block 1, 2D Convolution Layer
                - Output: 64 x 64 x 128
            - 2D Convolution, Kernel Size = 3, Activation = “Relu”
                - Output: 64 x 64 x 64
            - 2D Convolution, Kernel Size = 3, Activation = “Relu”
                - Output: 64 x 64 x 64
- Output Layers
    - 2D Convolution, Kernel Size = 3, Activation = “Relu”
        - Output: 64 x 64 x 2
    - 2D Convolution, Kernel Size = 3, Activation = “Sigmoid”
        - Output: 64 x 64 x 1

**Generation of Ground Truth Thresholds**

Unlike the generation of masks for the U-Net training set, the goal of ground truth thresholds is to generate the most accurate representation of the organelle morphology based on the data. Ground truth masks were generated using Otsu thresholding, but the implementation was sometimes adjusted to better match the fluorescence data. This is a drawback of fluorescent methods, where sometimes the boundary regions can be ambiguous or where staining is not uniform. For example, a slightly different approach was needed to properly threshold the nucleus in the two fluorescence images in Figure S30. The types of adjustments to the protocol are detailed below.

Fluorescent Ground Truth Adjustments from Simple Otsu Thresholding

- Removal of extracellular regions: 1. Otsu thresholding to remove the glass background. 2. Otsu thresholding on the cell identified in 1.
- Nuclear Thresholding: For speckles and nucleolus, the fluorescence was better reflected in the ground truth when the nucleus region was first defined using the nuclear stain and Otsu thresholding was applied to the speckle or nucleolus stain of that region.
- Cytosol Thresholding: Some cytoplasmic stains were better reflected in the ground truth when the nucleus region was first defined with the nuclear stain and then removed from the image. The Otsu thresholding was applied to the cytosol only.
- Uneven Nucleus and Cytoplasm Background: For a few cells the nonspecific background staining of the nucleus and cytoplasm were different. This was problematic for Otsu thresholding of organelle that spanned the nucleus and cytoplasm. Here, Otsu thresholding was applied to the nucleus and cytosol separately and then combined.

The choices made during these steps can be subjective. To help alleviate this concern we provide all of the fluorescent ground truth thresholds that were used to calculate the accuracy of our models next to the raw fluorescent images and predictions in Figure 3, Figure 4, and Figure S17 – S29.

**Model Evaluation**

Proper quantification of accuracy is challenging in images with high class imbalance, such as for organelles that occupy a small region within the cell. Metrics such as Intersection over Union (IoU) and F1, which try to better account for class imbalance can still fail to properly quantitatively represent the classification accuracy in cases similar to organelle prediction.[1] To account for the different utility in different accuracy metrics we have included several alongside the predictions in Figure S17-S29.

To minimize class imbalance during training, RF training sets contained an approximately equal amount of positive and negative labels for each class. The organelle classes and the size of the training data sets are provided in Table S2. To evaluate the performance of our RF models the data was sampled 250 times, with equal amounts of positive and negative labels for each class to account for potential sampling error and to stabilize the results (Fig. S6). The predicted label of each spectrum was compared to the real label. True positive (TP) assignments correctly predict the positive class. True negative (TN) assignments correctly predict the negative class. False positive (FP) assignments incorrectly predict the positive class. False negative (FN) assignments incorrectly predict the negative class. All metrics are evaluated within [0,1], and a higher value means better prediction. Values for the prediction of individual spectra with the RF model are reported in Table 1. Performance metrics for the hyperspectral maps for both the RF and U-Net model are reported in Fig. S17-S29, with the average accuracy from each map reported in Table 2.

The **Precision** of the model predictions were calculated with Equation S1:

$$Precision = \frac{\sum TP}{\sum(TP+FP)} \qquad \text{(S1)},$$

where TP is the number of true positives and FP is the number of false positives.

The **Recall** of the model predictions were calculated with Equation S2:

$$Recall = \frac{\sum TP}{\sum(TP+FN)} \quad \text{(S2)},$$

where TP is the number of true positives and FN is the number of false negatives.

The **F1 score** of the model predictions were calculated with Equation S3:

$$F1 = \frac{2xPrecisionxRecall}{Precision+Recall} \text{ (S3)},$$

using the Precision (ES1) and Recall (ES2) calculated above.

The **Accuracy** of the model predictions were calculated with Equation S4:

$$Accuracy = \frac{\sum(TP+TN)}{\sum(TP+FP+FN+TN)} \quad \text{(S4)},$$

where TP is the number of true positive assignments, TN is the number of true negative assignments, FP is the number of false positive assignments, and FN is the number of false positive assignments.

The **Specificity** of the model predictions were calculated with Equation S5:

$$Specificity = \frac{\sum TN}{\sum(TN+FP)} \qquad \text{(S5)},$$

where TN is the number of true negatives, TN is the number of true negatives, and FP is the number of false positives.

The **Intersection over Union (IoU)** of the model predictions were calculated with Equation S6:

$$IoU = \frac{\sum TP}{\sum(TP+FP+FN)} \qquad \text{(S6)},$$

where TP is the number of true positives, FP is the number of false positives, and FN is the number of false negatives.

The **Area Under the Curve (AuC)** is the area under the ROC curve and is calculated with the scikit-learn AuC function. The equation for this is presented below as Equation S7:

$$AUC = \sum_{i=1}^{n} \frac{(FPR_i - FPR_{i-1})(TPR_i - TPR_{i-1})}{2} \qquad \text{(S7)},$$

Where:

FPR is the false positive rate = FP / (FP + TN)

where FP is the false positives and TN is the true negatives.

TPR is the true positive rate = TP / (TP + FN)

where TP is the number of true positives and FN is false negatives.

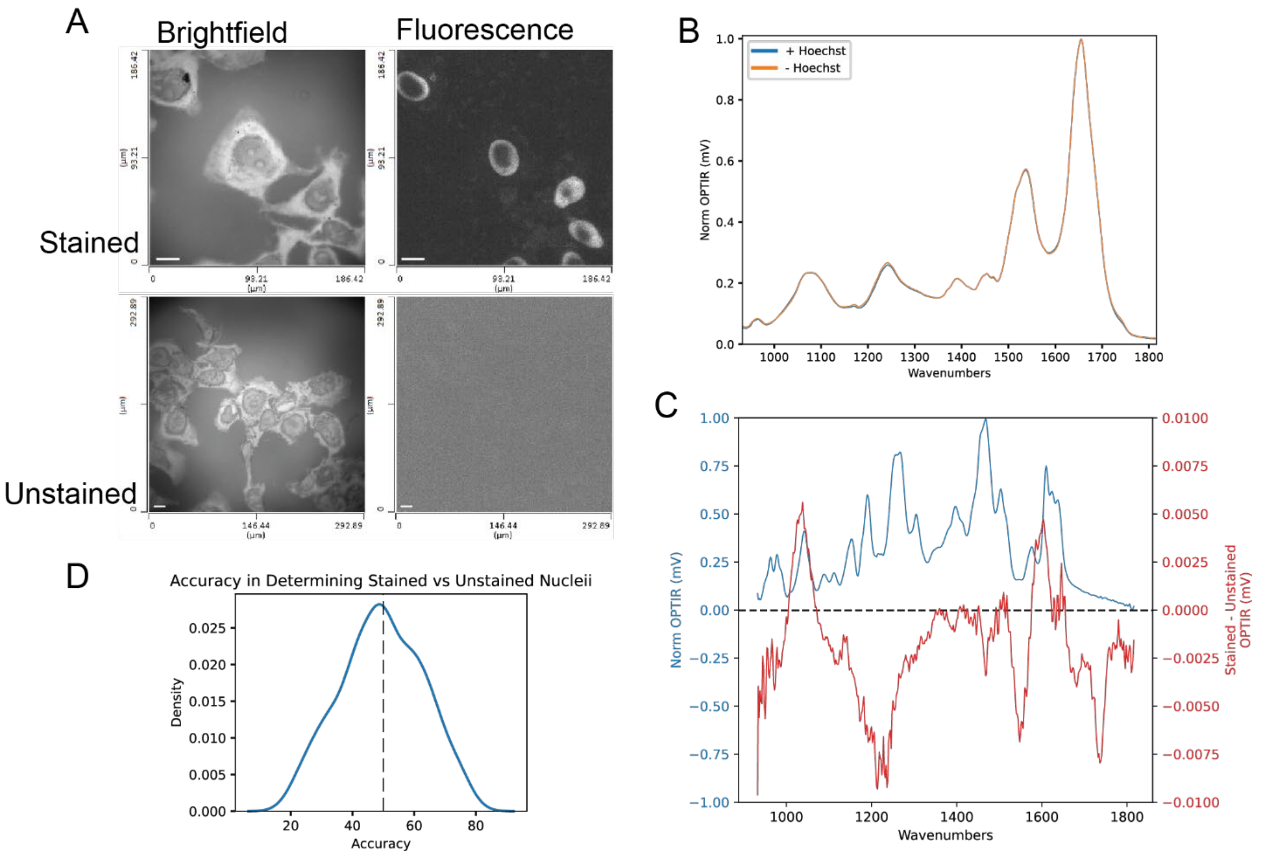


**Figure S1. Stain in not detected by IR**. A. Representative images of fixed U-2 OS cells stained with Hoechst (top) and unstained (bottom) (scale bar = 10 μm). B. The average of 550 OPTIR spectra from 11 Hoechst stained (blue) and 11 unstained cells (orange), with 50 spectra from each cell. C. The difference between the stained and unstained average spectra from panel B (red) is shown in comparison to the Hoescht OPTIR spectra (blue). D. To test whether the model is trained to identify the stain, six cells (3 stained and 3 unstained) were randomly selected and used to train a random forest model. The model was tested on the remaining dataset and the accuracy was recorded. This process was repeated 100 times. The distribution of the accuracies of the 100 trials is centered at 49.5% (vertical dashed line marks 50%).

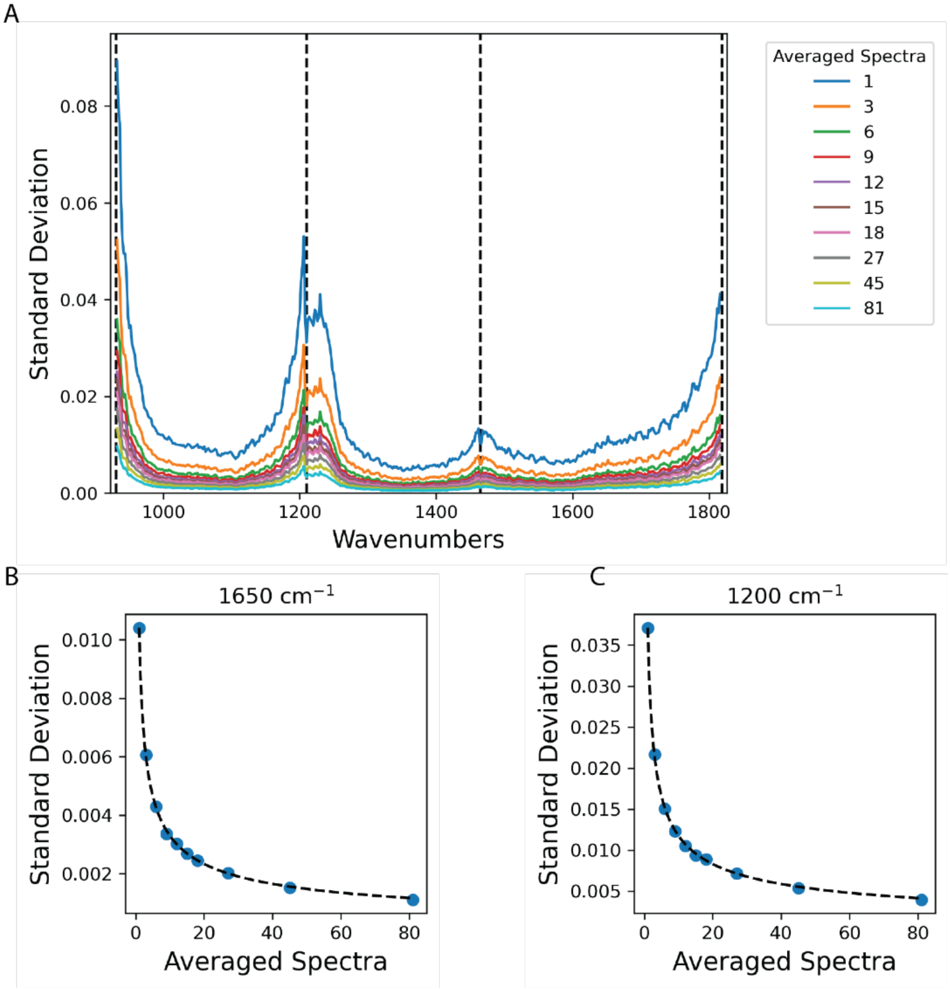


**Figure S2. Noise in the OPTIR spectra.** A. Average IR spectrum of a $CaF_2$ coverslip arising from between 1 and 81 measurements, drawn from a set of 250 calcium fluoride spectra. Vertical black dashed lines shown at 931, 1211, 1819, 1465 $cm^{-1}$ correspond to chip changes. Increasing the number of acquisitions lowers this noise for both low (B) and high (C) noise regions according to the square root of the number of samples.

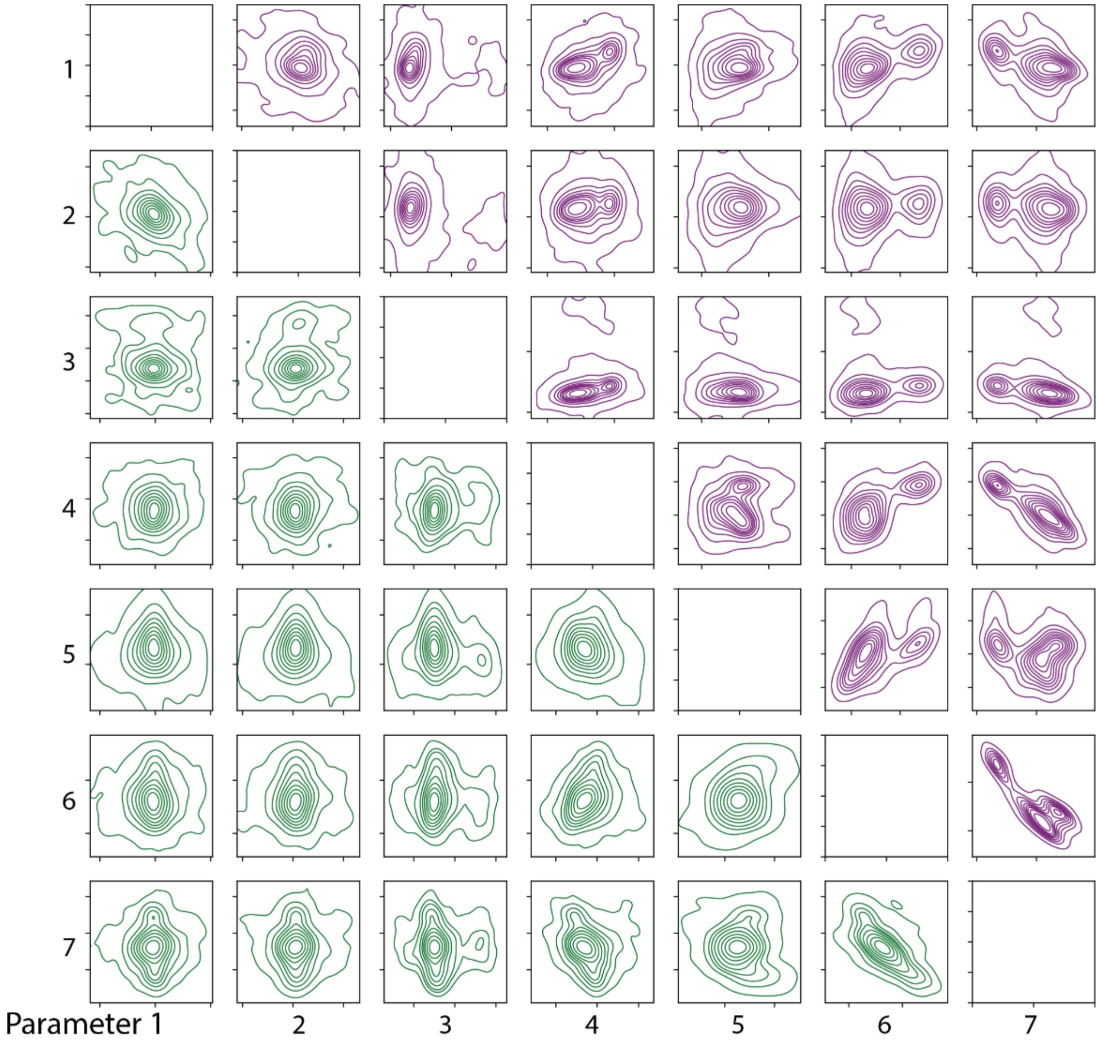


| # | Parameters | Biomolecule |
|---|---|---|
| 1 | Second Derivative at 1040 | Carbohydrate |
| 2 | Difference Second Derivative at 1063 and 1087 | Nucleic Acid |
| 3 | Peak Location at 1390 | Lipid |
| 4 | Signal Ratio at 1450 and 1464 | Lipid |
| 5 | Amide II : Amide I Ratio | Protein |
| 6 | Amide I FHWM | Protein |
| 7 | Signal at 1740 | Lipid |

**Figure S3. Correlation of Parameters of Nucleus Data**. To ensure that the selected parameters for the nucleus contain unique information the distributions for each parameter were plotted against the distribution for every other parameter. Each of the seven parameters (infrared feature in table) is compared with all of the other parameters and shown here with a representative cell with 100 spectra, equally divided between the cytoplasm and nucleus. The diagonal column is left blank as each distribution would perfectly correlate with itself. To facilitate viewing the cytoplasmic (green) and nuclear (purple) parameter distributions are separated, but all of the same comparisons are present in the dataset, across the diagonal. Displayed is representative correlation of parameters for the nucleus (purple) and the cytoplasm (green) for a cell from the training dataset.

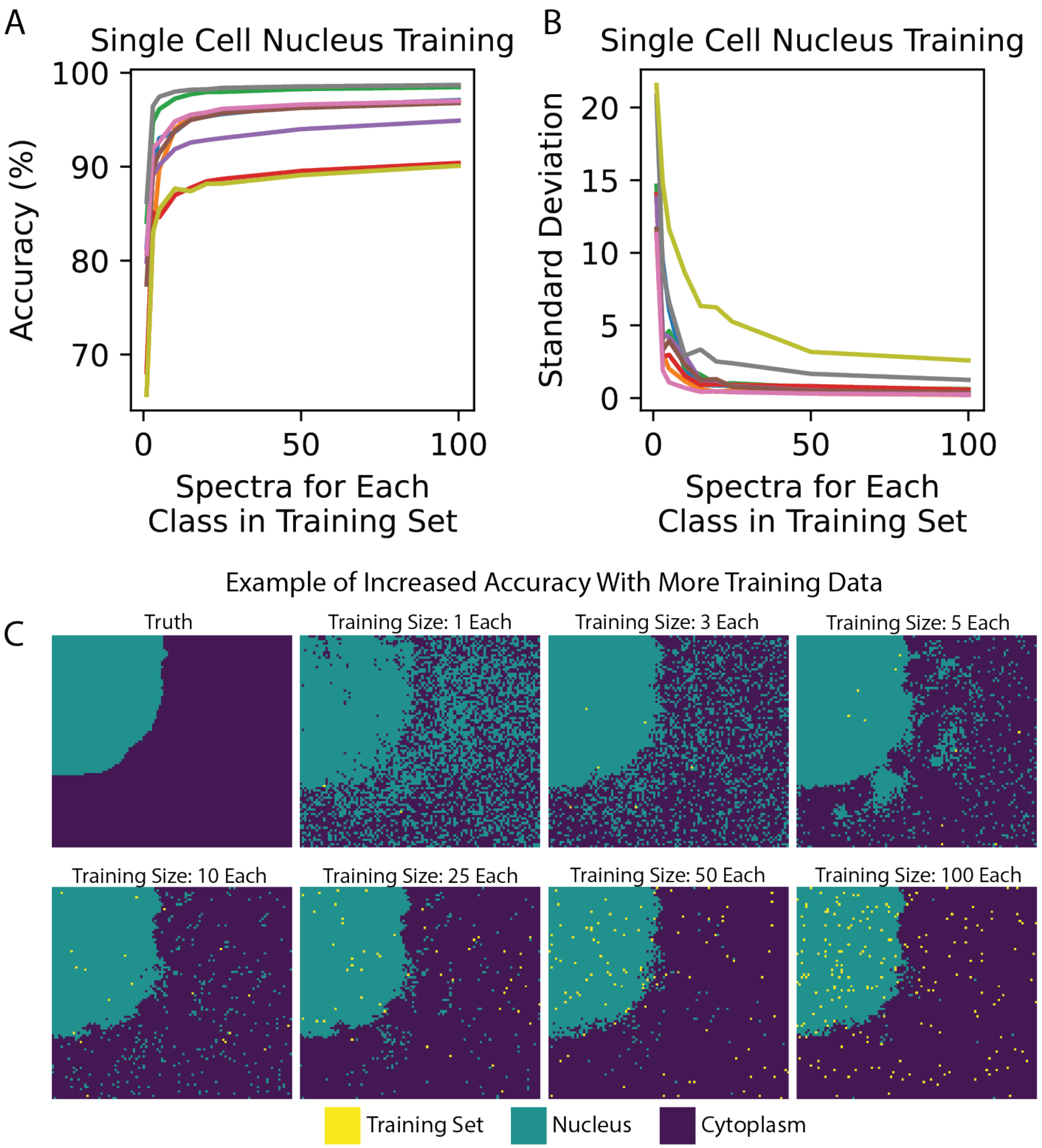


**Figure S4. Training on a Single Cell.** Fluorescently labeled hyperspectral maps of 9 cells were collected. The random forest model was trained on between 1 and 100 spectra randomly selected for each class (cytoplasm and nucleus) from a single cell and used to predict the class of the other spectra from the same cell. This process was repeated 250 times for each cell to evaluate (A) the average accuracy of the random forest models and (B) the standard deviation of the accuracy of the models. Each color is a different cell. Deviation is plotted separately to provide information about the potential effect of sampling bias during training dataset collection. C. The fluorescence data is thresholded to provide a ground truth for comparison to representative random forest predictions for a single cell trained on the specified number of spectra for each class (cytoplasm and nucleus). The yellow-colored pixels are used to train and predict the nucleus (green) or cytoplasm (blue) class of the rest of the cell image.

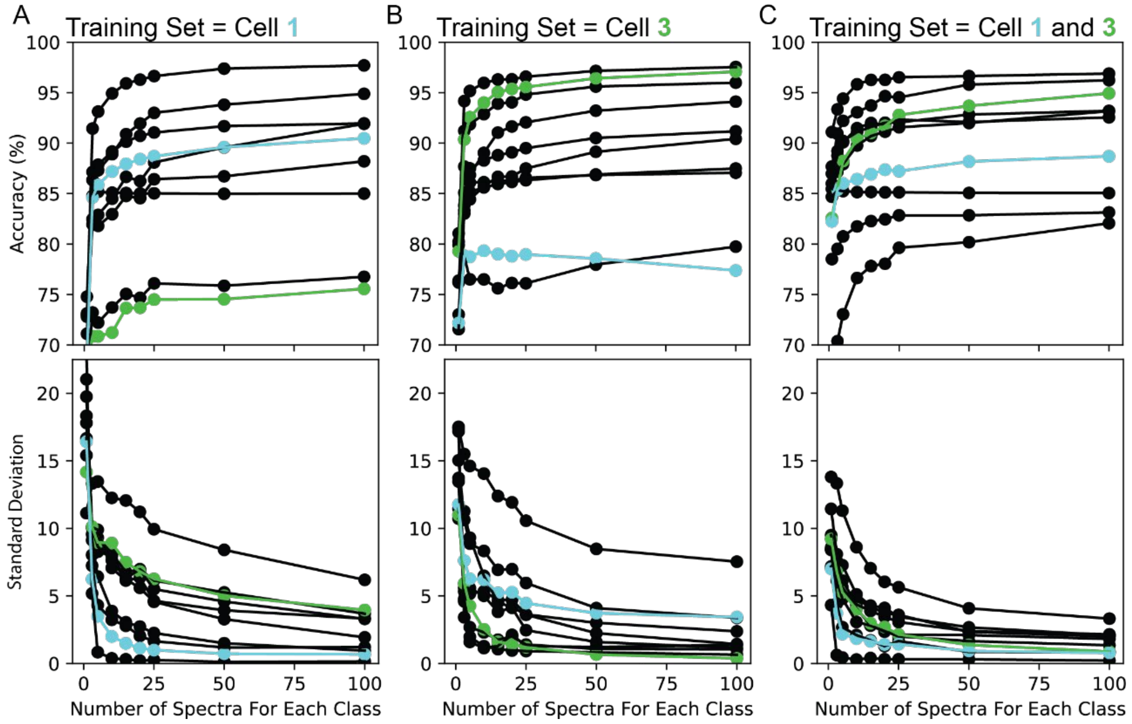


**Figure S5. Training on Multiple Cells.** A. Random Forest models were trained on between 1 and 100 spectra randomly selected from each class (nucleus and cytoplasm) of cell 1 (cyan). The model is used to predict the class of spectra from nine other cells. Displayed are the accuracy and the standard deviation of the prediction for a given number of training spectra. B. Random Forest models trained on between 1 and 100 spectra randomly selected from each class (nucleus and cytoplasm) of cell 3 (green). The model is used to predict the class of spectra from all nine cells. Displayed are the accuracy and the standard deviation of the prediction for a given number of training spectra. C. Random Forest model trained on between 1 and 100 spectra randomly selected from each class (nucleus and cytoplasm) of the combined datasets of cell 1 (blue) and cell 3 (green). The model is used to predict the class of spectra from all nine cells. Displayed are the accuracy and the standard deviation of the prediction for a given number of training spectra.

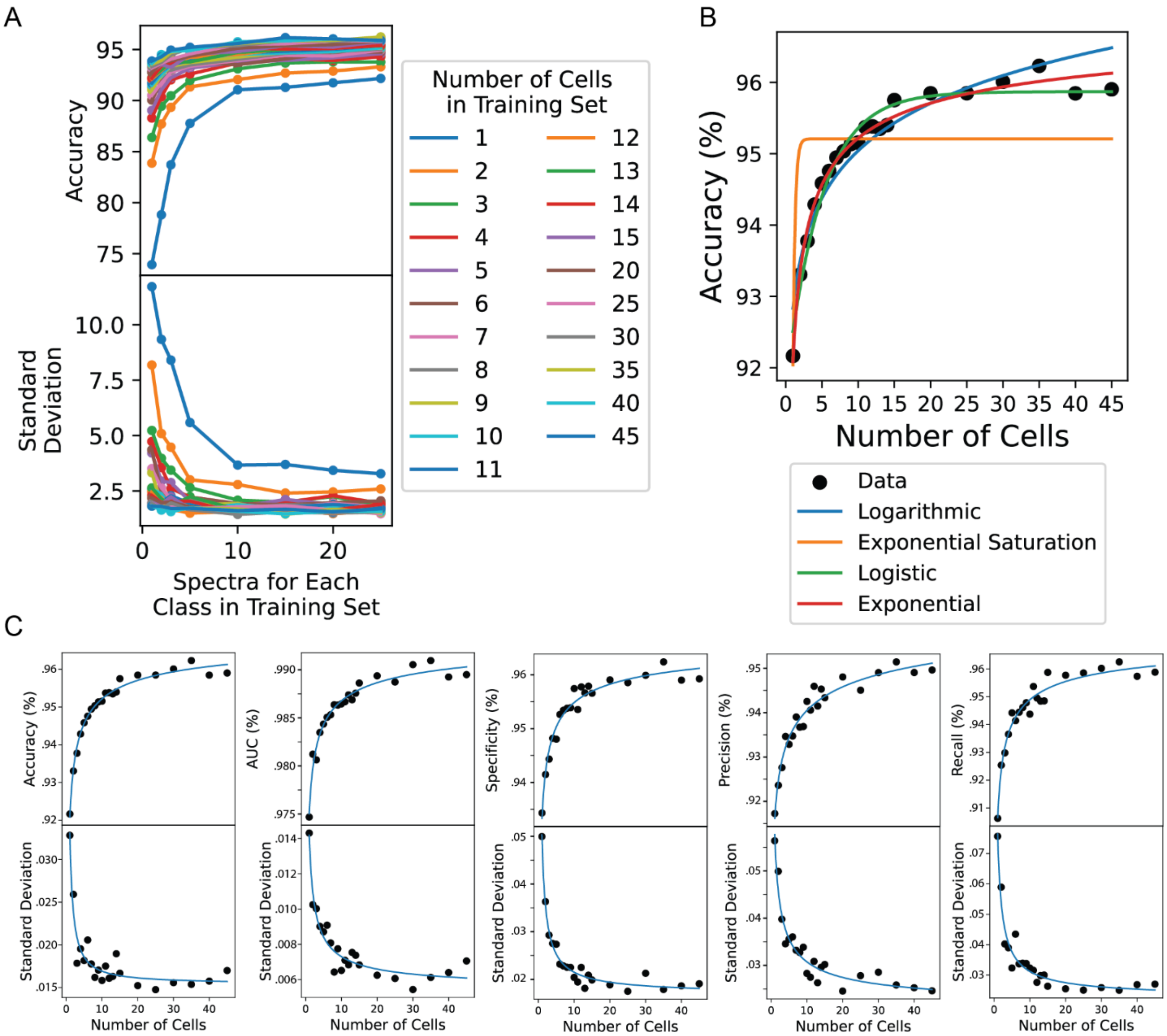


**Figure S6. Dependence of Training Accuracy on Quantity of Data.** Fluorescently labeled spectra for the nucleus and cytoplasm of 55 cells were collected. The 55 cells were randomly divided into training sets of 1 to 45 cells. The test set was 5 randomly selected cells not included in the training set. From the cells in the training set, a range of 1 to 25 randomly selected spectra for each class (cytoplasm and nucleus) was selected from each cell and used to train a random forest model. This process was repeated 250 times for each condition. A. Accuracy (top) and standard deviation (bottom) of spectra from between 1 and 45 cells in a training set predicting classification of 5 different cells in a test set. B. Different learning functions (logarithmic, exponential saturation, logistic, and exponential) are fit to the accuracy versus number of cells for 25 randomly selected spectra for each class to estimate the maximum accuracy of our approach with larger amounts of data. C. The model behavior is classified by (left to right) accuracy, area under the receiver operating curve (AUC), specificity, precision, and recall. All closely track the accuracy.

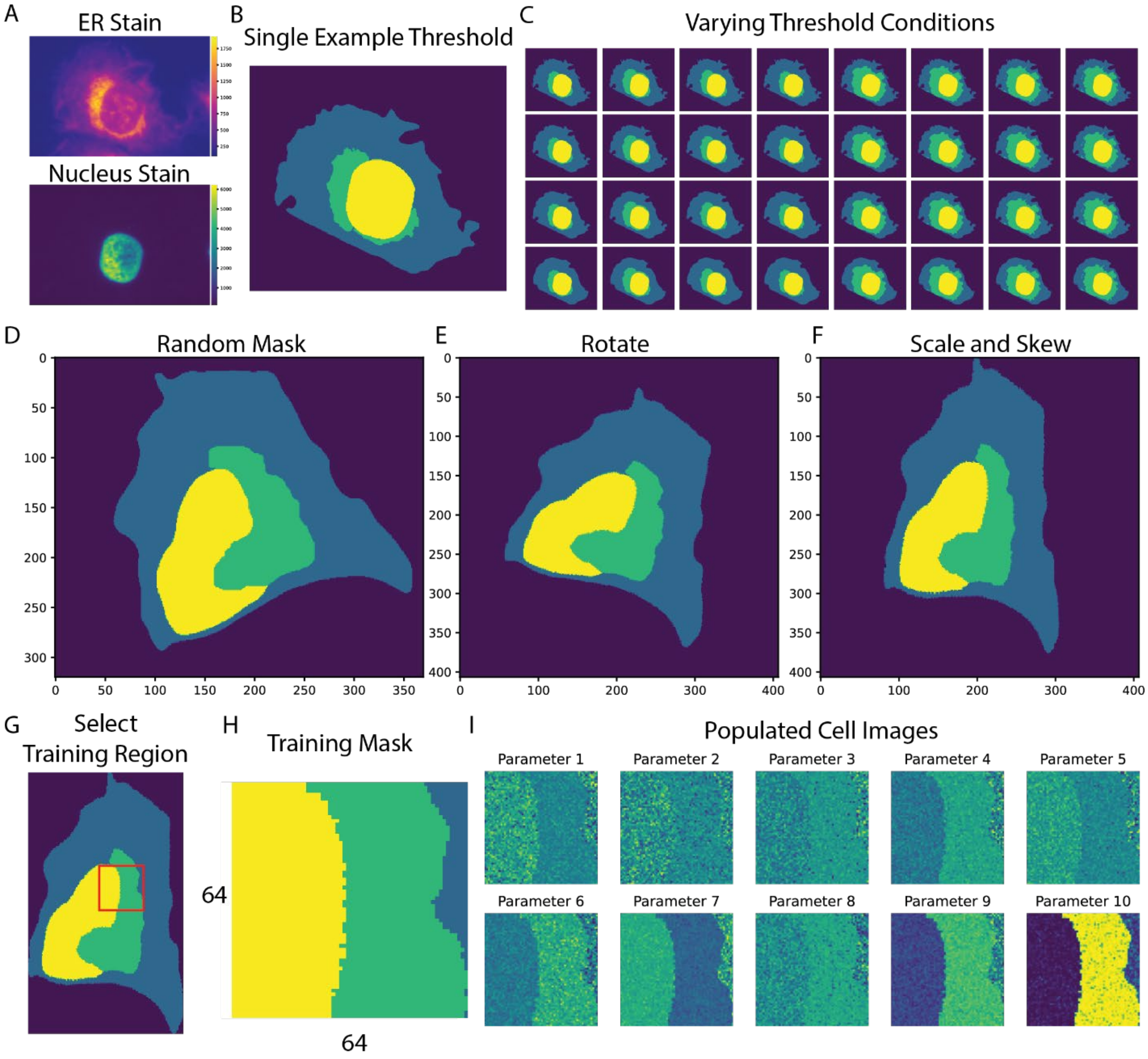


**Figure S7. Mask Generation.** A. Representative fluorescence image of cell stained with the different dyes [ER (red) and Nuclei (blue-green)]. B. The fluorescence image (A) is masked using Otsu's method to segment the organelle of interest ER (green), nucleus (yellow), and the cell boundary (blue). Here the cytoplasm is the ER (green) + the cell boundary (blue) C. To control for errors in thresholding and increase the variation of the masks the masks generated by Otsu's method were expanded and eroded. To account for overlap of organelles, duplicate masks with different mask ordering were generated. An example of this is given in D-G, which show the ER overlayed over the nucleus, whereas B-C show the nucleus over the ER. To generate additional synthetic masks for training, one mask was (D) chosen and then (E) rotated, scaled, and (F) skewed. From the synthetic masks, (G) a random 64x64 region was selected as (H) a training mask. I. The synthetic mask was then populated with data from the hyperspectral training dataset.

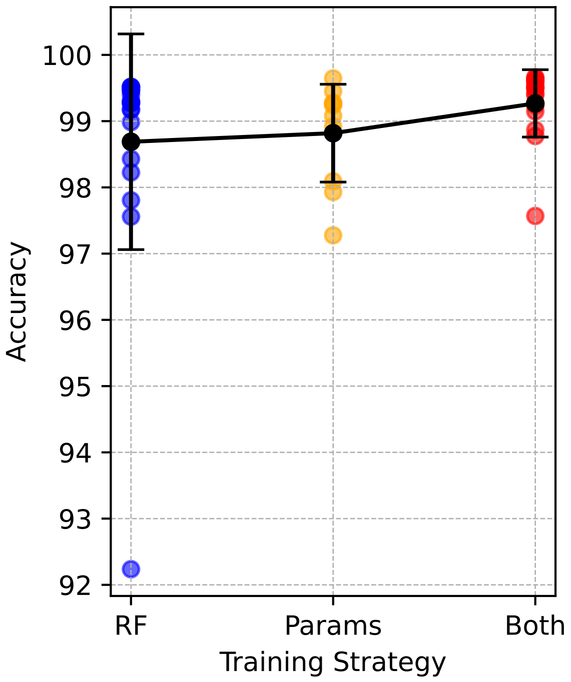


**Figure S8. Classification Accuracy of Supervised Learning Algorithms**. A CNN was trained with 3 different types of data: random forest (RF) scores, infrared parameters, or a combination of both for cytoplasmic/nucleus identification. The model was trained for 20 epochs, and the accuracy of converged models was compared. These models were trained with the dataset sizes detailed in Table S2.

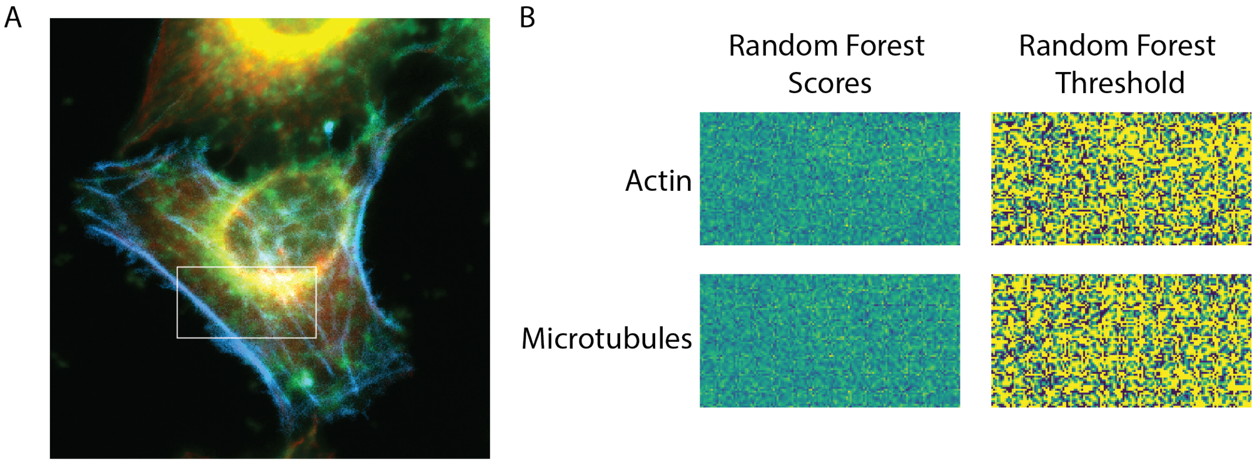


**Figure S9 Random Forest Prediction of Actin and Microtubules**. A. Fluorescent image of a U2-OS cell stained for actin (blue), microtubules (red), and mitochondria (green). B. Attempts at random forest classification result in accuracies of 39% for microtubules and 26% for actin. The actin training set size was 14 cells with 700 positive and 700 negative examples. The microtubule training set size was 20 cells with 857 positive examples and 750 negative examples.

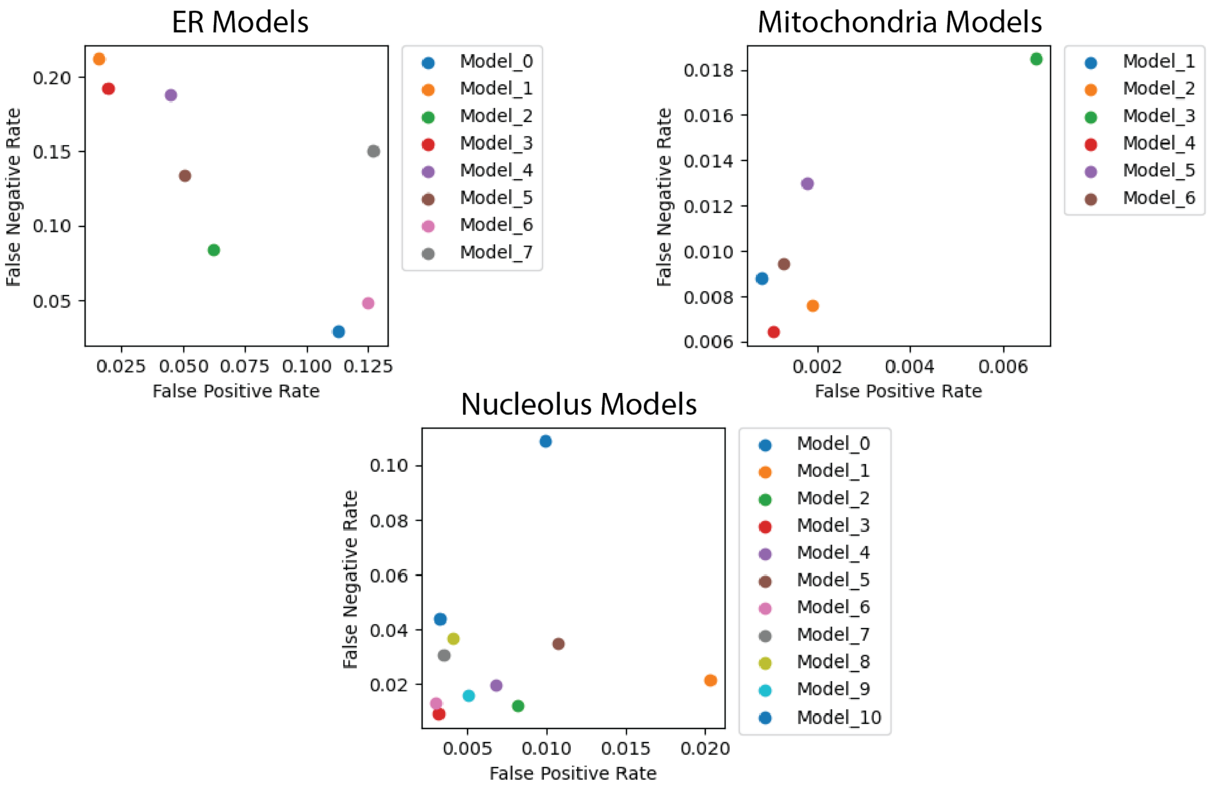


**Figure S10. CNN Model Selection.** Multiple CNN models were trained in parallel for the ER, mitochondria, and nucleolus. Displayed are the false negative versus false positive rate for each model. The final CNN models chosen are 2 for the ER, 4 for the mitochondria, and 3 for the nucleolus.

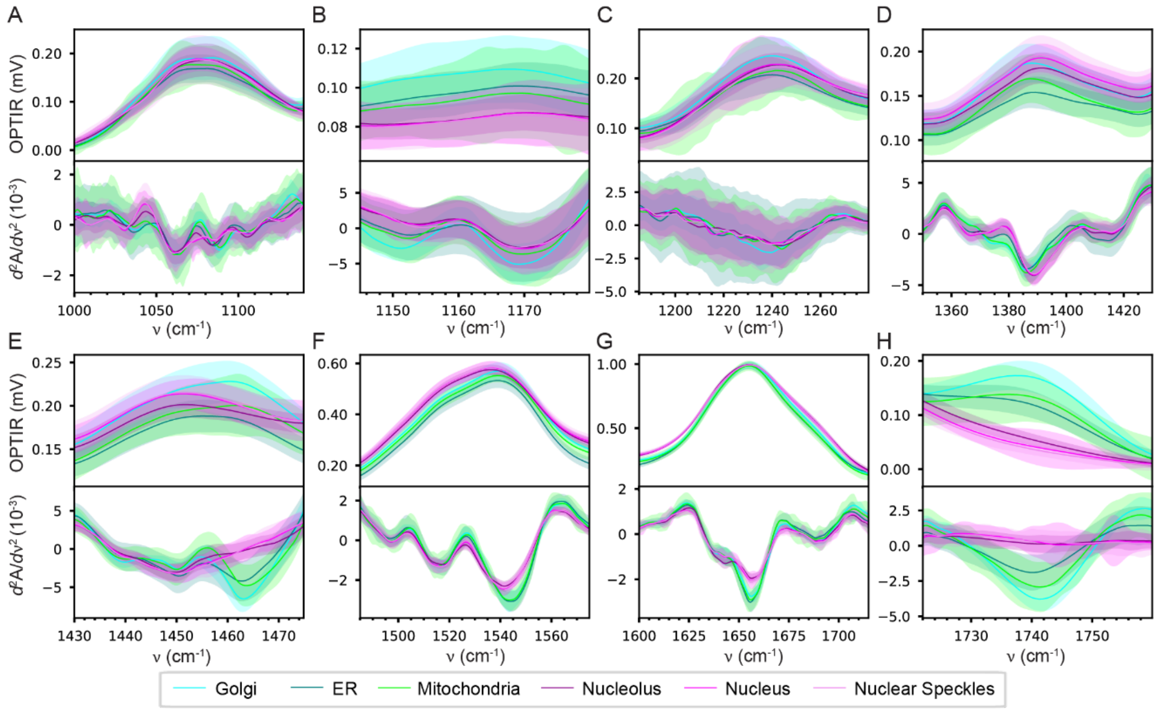


**Figure S11. OPTIR spectra of subcellular organelles. A.–H.** Averaged OPTIR spectra (top panels) and corresponding second derivative spectra (bottom panels) for pixels classified as Golgi (cyan), ER (blue), mitochondria (green), nucleolus (purple), nucleus (magenta), and nuclear speckles (lavender). Shaded areas indicate the standard deviation of the spectra. The number of pixels per organelle are detailed in Table S2.

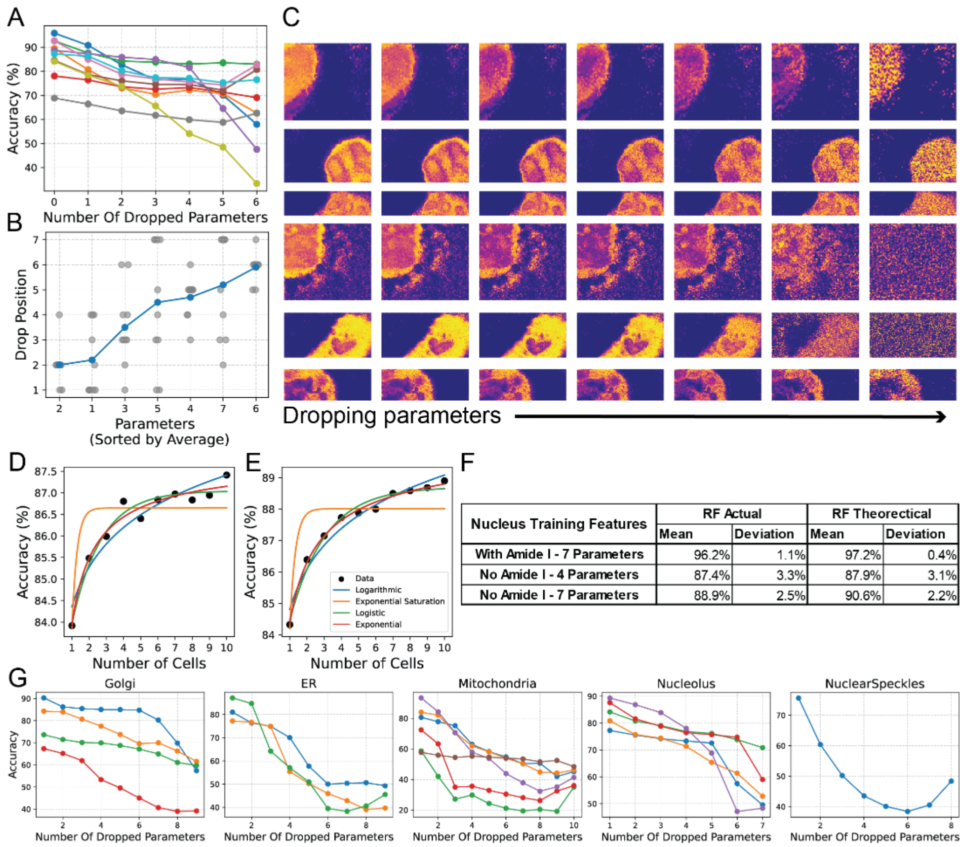


| Nucleus Training Features | RF Actual | | RF Theorectical | |
|---|---|---|---|---|
| | Mean | Deviation | Mean | Deviation |
| With Amide I - 7 Parameters | 96.2% | 1.1% | 97.2% | 0.4% |
| No Amide I - 4 Parameters | 87.4% | 3.3% | 87.9% | 3.1% |
| No Amide I - 7 Parameters | 88.9% | 2.5% | 90.6% | 2.2% |

**Figure S12. Parameter Drop Test.** A. Fluorescently labeled hyperspectral maps of 9 cells were collected. Random forest training using all of the spectra from the training dataset (nucleus and cytoplasm) was performed with all the parameters and tested on itself to establish a baseline accuracy. The training was then repeated by dropping each parameter, training the model with one less parameter. The accuracy for each of these was compared and the parameter that caused the smallest accuracy loss was then permanently dropped and the process was repeated until only one parameter was left. B. Plot of the order that parameters were dropped versus the infrared parameter that was dropped. C. Representative cell visualizing the effect of the loss of accuracy in the random forest scores as the parameters are dropped. D. This test was repeated on each of the other organelles. Each color indicates a different cell.

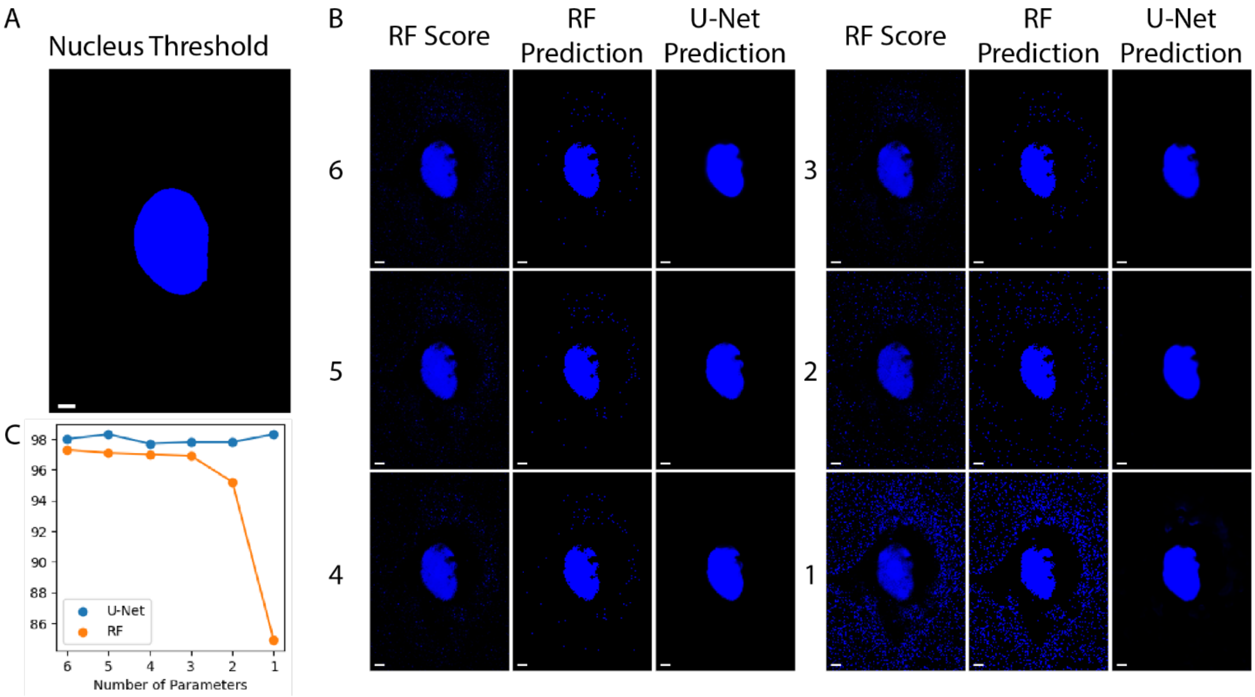


**Figure S13. Nucleus Parameter Drop Test.** Using the order of importance of parameters from Figure S12, parameters were dropped from training, with the least important parameters dropped first. A. The target example of the nucleus ground truth for comparison to the predictions. B. The RF score, RF prediction, and U-Net prediction, for fewer parameters from 6 to 1 are shown. The number next to the images tracks how many parameters are remaining from the original 7 parameter set. C. The tracked accuracy for each of the U-Net and RF predictions across the presented results.

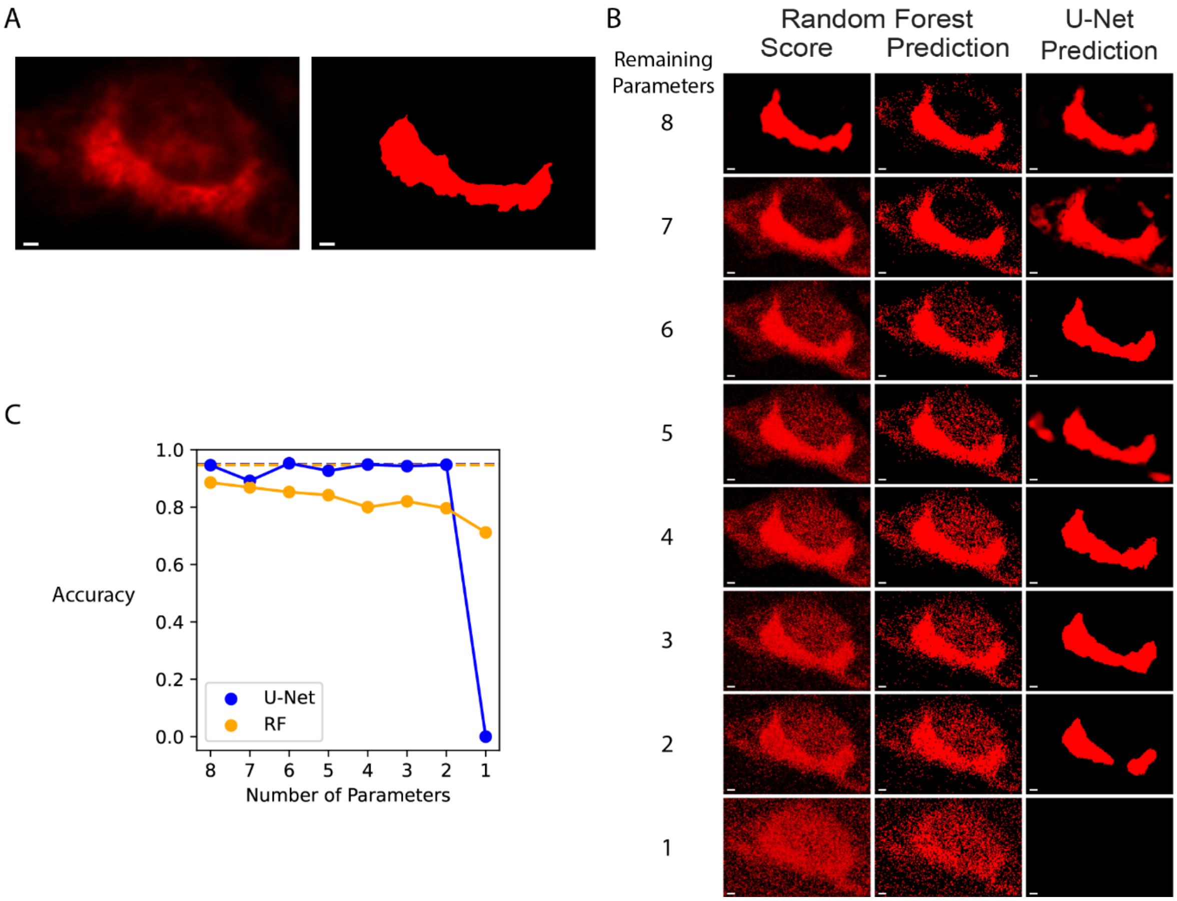


**Figure S14. ER Parameter Drop Test.** The number of parameters needed for the ER prediction was tested. A. The fluorescence (*left*) and ground truth (*right*) (scale bar = 10 μm) are shown for the ER to serve as a target for the predictions. B. Using the order of importance of parameters from Figure S12, parameters were dropped with the least important parameters dropped first. The ER was classified with reasonable accuracy until only 1 parameter remained. The singular parameter case generated extremely noisy RF classifications and did not give U-Net prediction for this cell. C. The tracked accuracy for each of the U-Net (blue) and RF (orange) predictions in B.

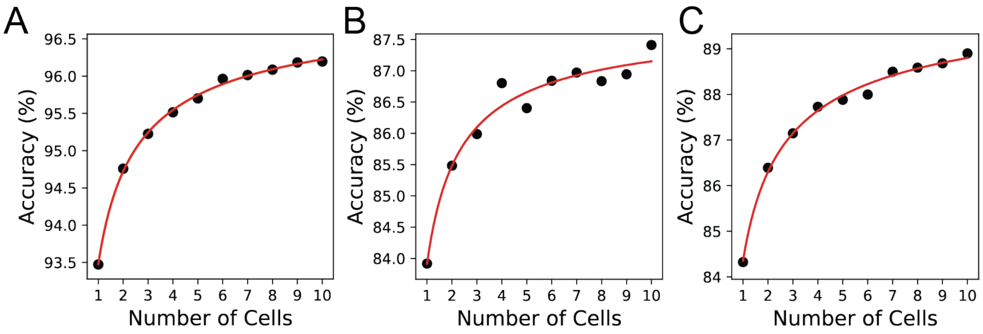


D

| Nucleus Training Features | RF Actual | | RF Theorectical | |
|---|---|---|---|---|
| | Mean | Deviation | Mean | Deviation |
| A With Amide I - 7 Parameters | 96.2% | 1.1% | 97.2% | 0.4% |
| B No Amide I - 4 Parameters | 87.4% | 3.3% | 87.9% | 3.1% |
| C No Amide I - 7 Parameters | 88.9% | 2.5% | 90.6% | 2.2% |

**Figure S15. Amide I Parameters for Nucleus Classification.** A. Accuracy versus number of cells for 25 randomly selected spectra for each class (cytoplasm and nucleus) trained on a random forest model with seven infrared features. (Data and best fit from Fig. S6E) B. Random Forest performance after removing infrared features that overlap with the strong water absorption in the mid-IR. Parameters removed from A include the amide II: amide I ratio, FWHM of the amide I, and lipid signal at 1740 $cm^{-1}$. C. Random Forest performance after adding three additional parameters to B. Additional parameters that were added include the second derivative at 1010 $cm^{-1}$, the second derivative at 1030 $cm^{-1}$, and the signal at 1170 $cm^{-1}$. Data in A-C fit to an exponential. D. Summary of the observed and theoretical values for the random forest accuracy and standard deviation of the models in A-C.

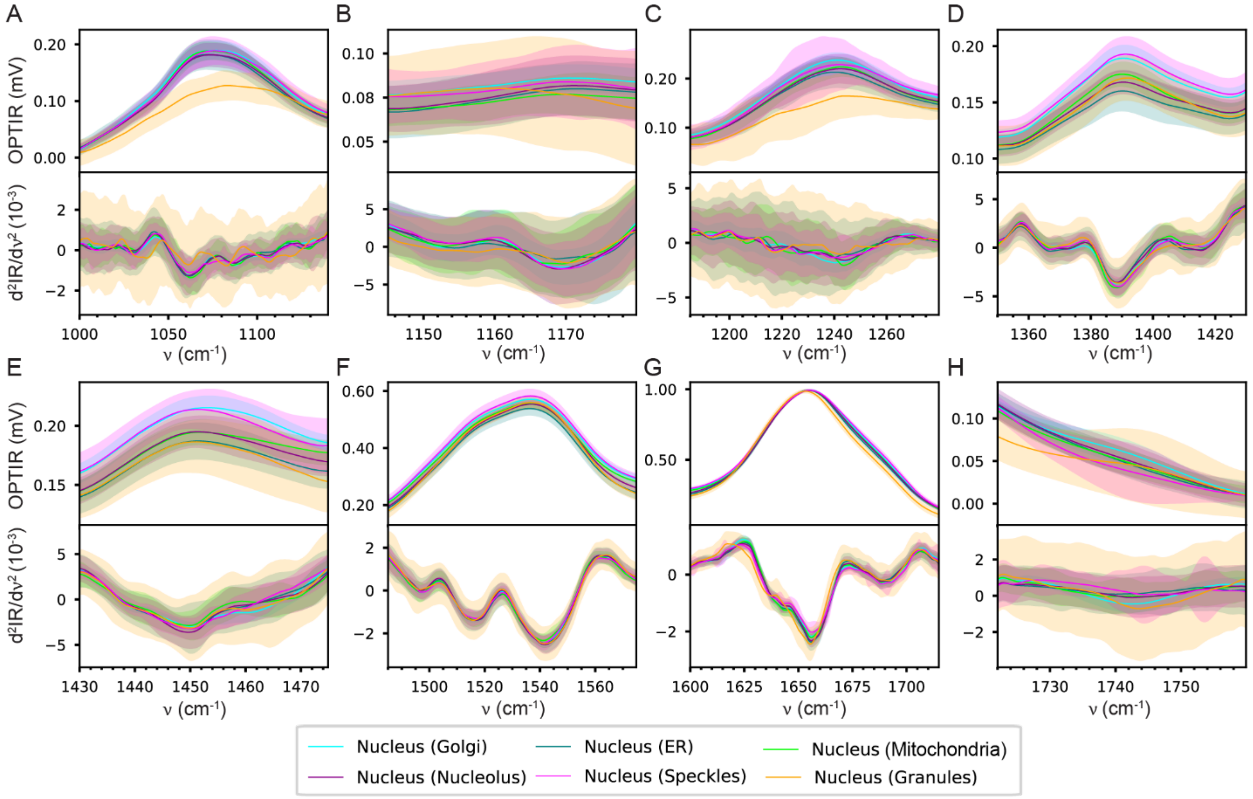


**Figure S16. Effect of Oxidative Stress on Nuclear Signals.** (top) Average OPTIR spectra and (bottom) second derivative of the average OPTIR spectra over the defined spectral region. Nuclei in cells that have undergone oxidative stress (orange) show significant spectral differences to the nuclei from all other collected training datasets. Shaded regions represent the standard deviation of the classified spectra. Dataset sizes for each are given in Table S2.

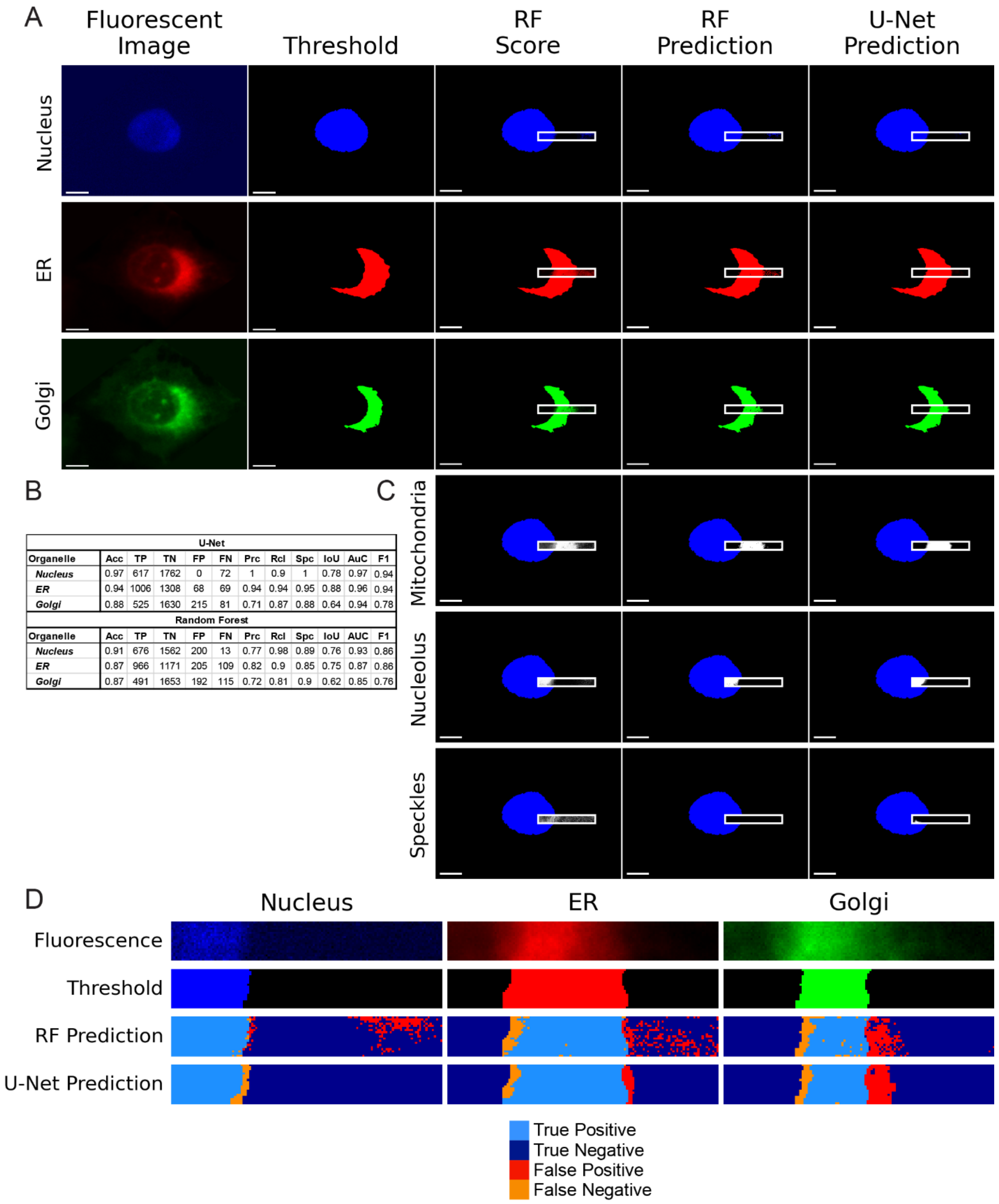


| U-Net | | | | | | | | | | | |
|---|---|---|---|---|---|---|---|---|---|---|---|
| Organelle | Acc | TP | TN | FP | FN | Prc | Rcl | Spc | IoU | AuC | F1 |
| *Nucleus* | 0.97 | 617 | 1762 | 0 | 72 | 1 | 0.9 | 1 | 0.78 | 0.97 | 0.94 |
| *ER* | 0.94 | 1006 | 1308 | 68 | 69 | 0.94 | 0.94 | 0.95 | 0.88 | 0.96 | 0.94 |
| *Golgi* | 0.88 | 525 | 1630 | 215 | 81 | 0.71 | 0.87 | 0.88 | 0.64 | 0.94 | 0.78 |
| **Random Forest** | | | | | | | | | | | |
| Organelle | Acc | TP | TN | FP | FN | Prc | Rcl | Spc | IoU | AUC | F1 |
| *Nucleus* | 0.91 | 676 | 1562 | 200 | 13 | 0.77 | 0.98 | 0.89 | 0.76 | 0.93 | 0.86 |
| *ER* | 0.87 | 966 | 1171 | 205 | 109 | 0.82 | 0.9 | 0.85 | 0.75 | 0.87 | 0.86 |
| *Golgi* | 0.87 | 491 | 1653 | 192 | 115 | 0.72 | 0.81 | 0.9 | 0.62 | 0.85 | 0.76 |

**Figure S17. Partial Cell Hyperspectral Map 1.** A. RF and U-Net predictions for stained organelles. The two leftmost columns display fluorescence images and the fluorescent threshold (scale bars: 10 μm). In columns three-five, a white box is drawn around the region where OPTIR data was collected. The third column shows the random forest confidence scores, which provide information on the certainty of the random forest model. The fourth column shows classification results generated by the random forest model. The final column presents predictions from the U-Net model, which was trained using a combination of random forest scores and IR features. Training dataset sizes are provided in Table S2. Each row shows a different organelle. Row 1: Nucleus (Blue), Row 2: ER (Red), and Row 3: Golgi (green). B. Table of accuracies of the RF and U-Net predictions for stained organelles in A. Accuracy (Acc), True Positive (TP), True Negative (TN), False Positive (FP), and False Negative (FN) are presented first. This is followed by derived performance metrics Precision (Prc), Recall (Rcl), Specificity (Spc), Intersection over Union (AoU), Area under the Curve (AuC), and F1 scores (F1). C. RF and U-Net predictions for unstained organelles in the region OPTIR data was collected. To give spatial context to the location within the cell, the nucleus fluorescence threshold is shown in blue. In line with the columns in A, the third column shows the RF scores, the fourth column shows the RF prediction, and the fifth column shows the U-Net prediction. The predictions are displayed in grayscale to distinguish them from organelles with fluorescent comparison. Row 1: Mitochondria, Row 2: Nucleolus, Row 3: Nuclear Speckles. D. Mapping classification errors for the stained organelles. To show where errors in each model occur, the cellular region where OPTIR data was collected is enlarged (the regions outlined in white in panel A). The top two rows display the fluorescence and fluorescence threshold from the region. The bottom two rows show the RF and U-Net predictions, colored according to the accuracy of the prediction. Pixels correctly predicted are shown in blue, with light blue for true positives and dark blue for true negatives. Pixels predicted incorrectly are shown in red for false positives and orange for false negatives. The organelles shown here by column are the same organelles as A. Column 1: Nucleus (Blue), Column 2: ER (Red), and Column 3: Golgi (green).

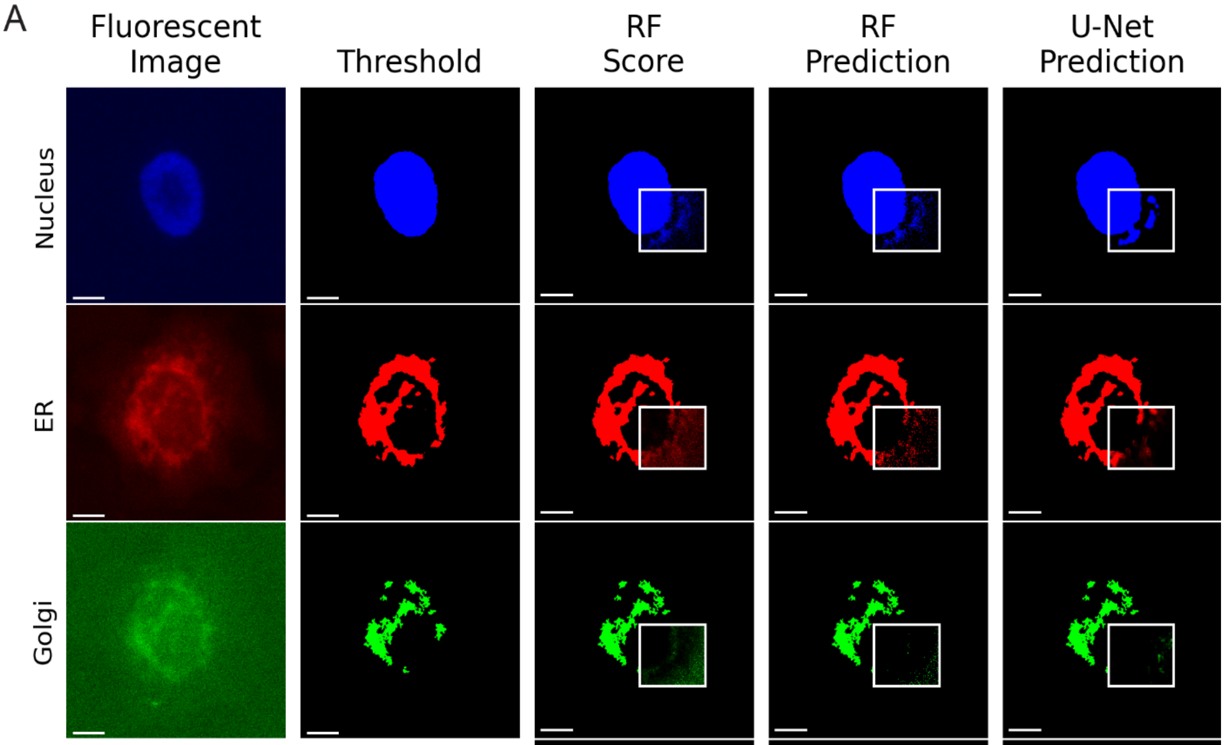


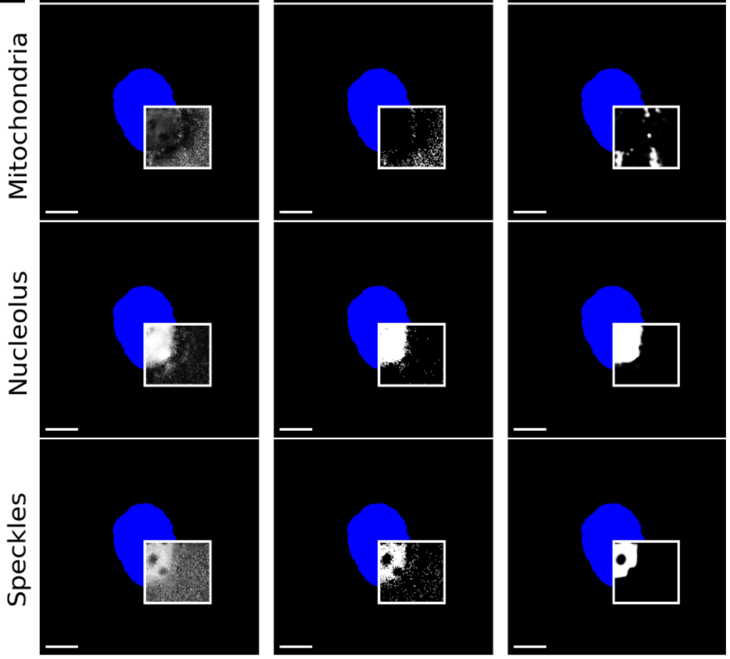


B

| U-Net | | | | | | | | | | | |
|---|---|---|---|---|---|---|---|---|---|---|---|
| Organelle | Acc | TP | TN | FP | FN | Prc | Rcl | Spc | IoU | AUC | F1 |
| ***Nucleus*** | 0.89 | 2875 | 5467 | 912 | 119 | 0.76 | 0.96 | 0.86 | 0.74 | 0.94 | 0.85 |
| ***ER*** | 0.81 | 657 | 6962 | 1386 | 368 | 0.32 | 0.64 | 0.83 | 0.27 | 0.82 | 0.43 |
| ***Golgi*** | 0.97 | 0 | 9138 | 0 | 235 | N/A | 0 | 1 | 0.00 | 0.55 | 0.00 |
| **Random Forest** | | | | | | | | | | | |
| **Organelle** | **Acc** | **TP** | **TN** | **FP** | **FN** | **Prc** | **Rcl** | **Spc** | **IoU** | **AUC** | **F1** |
| ***Nucleus*** | 0.86 | 2858 | 5223 | 1156 | 136 | 0.71 | 0.95 | 0.82 | 0.69 | 0.89 | 0.82 |
| ***ER*** | 0.81 | 380 | 7175 | 1173 | 645 | 0.24 | 0.37 | 0.86 | 0.17 | 0.62 | 0.29 |
| ***Golgi*** | 0.96 | 0 | 8994 | 144 | 235 | N/A | 0 | 0.98 | 0.00 | 0.49 | 0.00 |

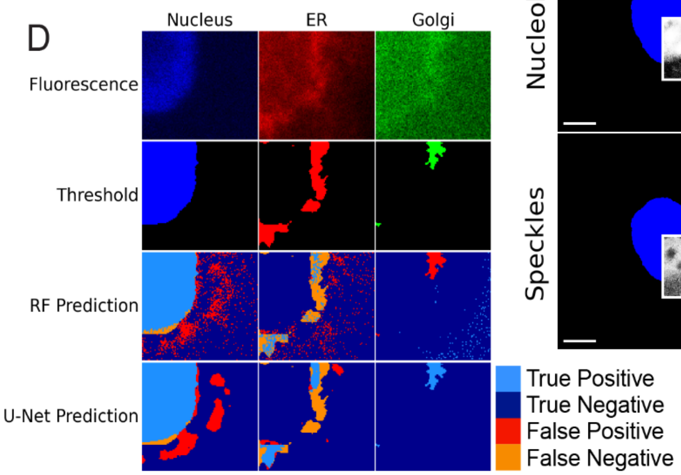


**Figure S18. Partial Cell Hyperspectral Map 2.** A. RF and U-Net predictions for stained organelles. The two leftmost columns display fluorescence images and the fluorescent threshold (scale bars: 10 μm). In columns three-five, a white box is drawn around the region where OPTIR data was collected. The third column shows the random forest confidence scores, which provide information on the certainty of the random forest model. The fourth column shows classification results generated by the random forest model. The final column presents predictions from the U-Net model, which was trained using a combination of random forest scores and IR features. Training dataset sizes are provided in Table S2. Each row shows a different organelle. Row 1: Nucleus (Blue), Row 2: ER (Red), and Row 3: Golgi (green). B. Table of accuracies of the RF and U-Net predictions for stained organelles in A. Accuracy (Acc), True Positive (TP), True Negative (TN), False Positive (FP), and False Negative (FN) are presented first. This is followed by derived performance metrics Precision (Prc), Recall (Rcl), Specificity (Spc), Intersection over Union (IoU), Area under the Curve (AuC), and F1 scores (F1). C. RF and U-Net predictions for unstained organelles in the region OPTIR data was collected. To give spatial context to the location within the cell, the nucleus fluorescence threshold is shown in blue. In line with the columns in A, the third column shows the RF scores, the fourth column shows the RF prediction, and the fifth column shows the U-Net prediction. The predictions are shown in grayscale to distinguish them from organelles with fluorescent comparison. Row 1: Mitochondria, Row 2: Nucleolus, Row 3: Nuclear Speckles. D. Mapping classification errors for the stained organelles. To show where errors in each model occur, the cellular region where OPTIR was collected is enlarged (the regions outlined in white in panel A). The top two rows display the fluorescence and fluorescence threshold from the region. The bottom two rows show the RF and U-Net predictions, colored according to the accuracy of the prediction. Pixels correctly predicted are shown in blue, with light blue for true positives and dark blue for true negatives. Pixels predicted incorrectly are shown in red for false positives and orange for false negatives. The organelles shown here by column are the same organelles as A. Column 1: Nucleus (Blue), Column 2: ER (Red), and Column 3: Golgi (green).

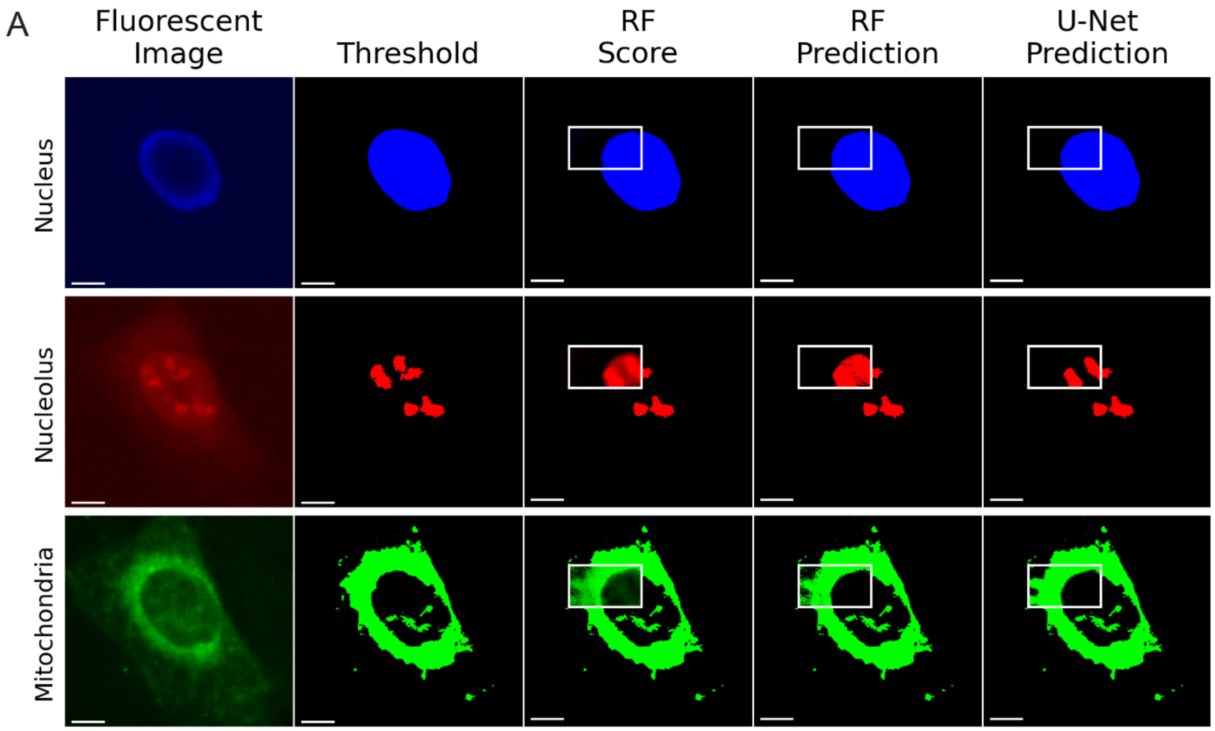


B

| U-Net | | | | | | | | | | | |
|---|---|---|---|---|---|---|---|---|---|---|---|
| Organelle | Acc | TP | TN | FP | FN | Prc | Rcl | Spc | IoU | AUC | F1 |
| *Nucleus* | 0.93 | 3022 | 3514 | 0 | 504 | 1 | 0.86 | 1 | 0.86 | 0.95 | 0.92 |
| *Nucleolus* | 0.91 | 893 | 5538 | 232 | 377 | 0.79 | 0.7 | 0.96 | 0.59 | 0.91 | 0.75 |
| *Mitochondria* | 0.91 | 3018 | 3388 | 362 | 272 | 0.89 | 0.92 | 0.9 | 0.83 | 0.98 | 0.90 |

| Random Forest | | | | | | | | | | | |
|---|---|---|---|---|---|---|---|---|---|---|---|
| Organelle | Acc | TP | TN | FP | FN | Prc | Rcl | Spc | IoU | AUC | F1 |
| *Nucleus* | 0.92 | 2961 | 3501 | 13 | 565 | 1 | 0.84 | 1 | 0.84 | 0.92 | 0.91 |
| *Nucleolus* | 0.82 | 1142 | 4607 | 1163 | 128 | 0.5 | 0.9 | 0.8 | 0.47 | 0.85 | 0.64 |
| *Mitochondria* | 0.89 | 2831 | 3452 | 298 | 459 | 0.9 | 0.86 | 0.92 | 0.79 | 0.89 | 0.88 |

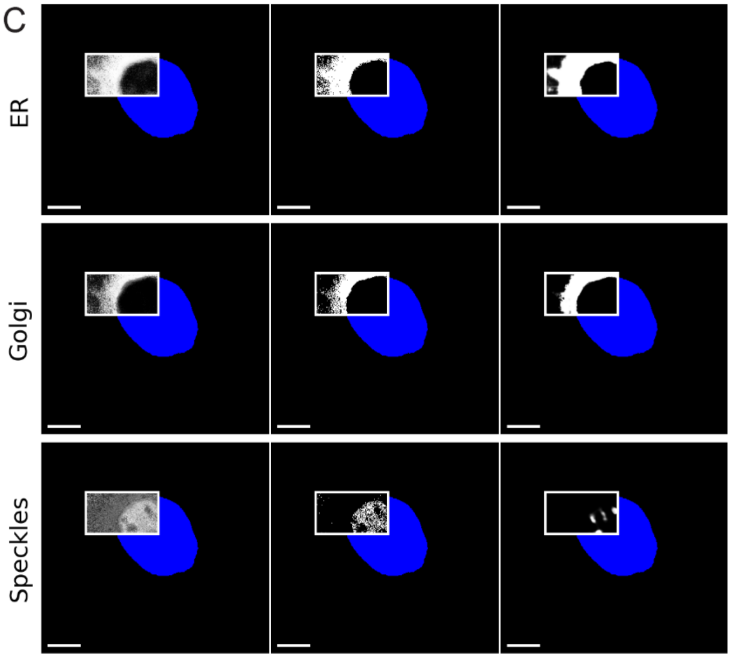


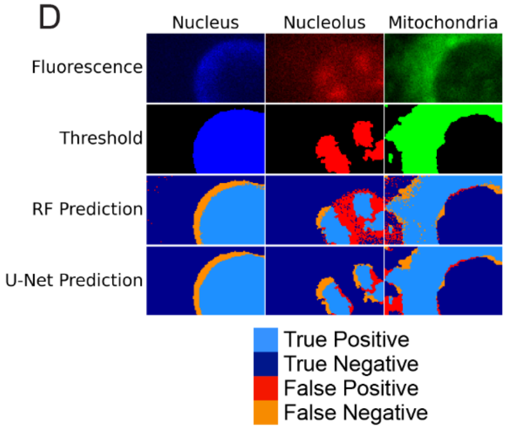


**Figure S19. Partial Cell Hyperspectral Map 3.** A. RF and U-Net predictions for stained organelles. The two leftmost columns display fluorescence images and the fluorescent threshold (scale bars: 10 μm). In columns three-five, a white box is drawn around the region where OPTIR data was collected. The third column shows the random forest confidence scores, which provide information on the certainty of the random forest model. The fourth column shows classification results generated by the random forest model. The final column presents predictions from the U-Net model, which was trained using a combination of random forest scores and IR features. Training dataset sizes are provided in Table S2. Each row shows a different organelle. Row 1: Nucleus (Blue), Row 2: Nucleolus (Red), and Row 3: Mitochondria (green). B. Table of accuracies of the RF and U-Net predictions for stained organelles in A. Accuracy (Acc), True Positive (TP), True Negative (TN), False Positive (FP), and False Negative (FN) are presented first. This is followed by derived performance metrics, Precision (Prc), Recall (Rcl), Specificity (Spc), Intersection over Union (IoU), Area under the Curve (AuC), and F1 scores (F1). C. RF and U-Net predictions for unstained organelles in the region OPTIR data was collected. To give spatial context to the location within the cell, the nucleus fluorescence threshold is shown in blue. In line with the columns in A, the third column shows the RF scores, the fourth column shows the RF prediction, and the fifth column shows the U-Net prediction. The predictions are shown in grayscale to distinguish them from organelles with fluorescent comparison. Row 1: ER, Row 2: Golgi, Row 3: Nuclear Speckles. D. Mapping classification errors for the stained organelles. To show where errors in each model occur, the cellular region where OPTIR data was collected is enlarged (the regions outlined in white in panel A). The top two rows display the fluorescence and fluorescence threshold from the region. The bottom two rows show the RF and U-Net predictions, colored according to the accuracy of the prediction. Pixels correctly predicted are shown in blue, with light blue for true positives and dark blue for true negatives. Pixels predicted incorrectly are shown in red for false positives and orange for false negatives. The organelles shown here by column are the same organelles as A. Column 1: Nucleus (Blue), Column 2: Nucleolus (Red), and Column 3: Mitochondria (green).

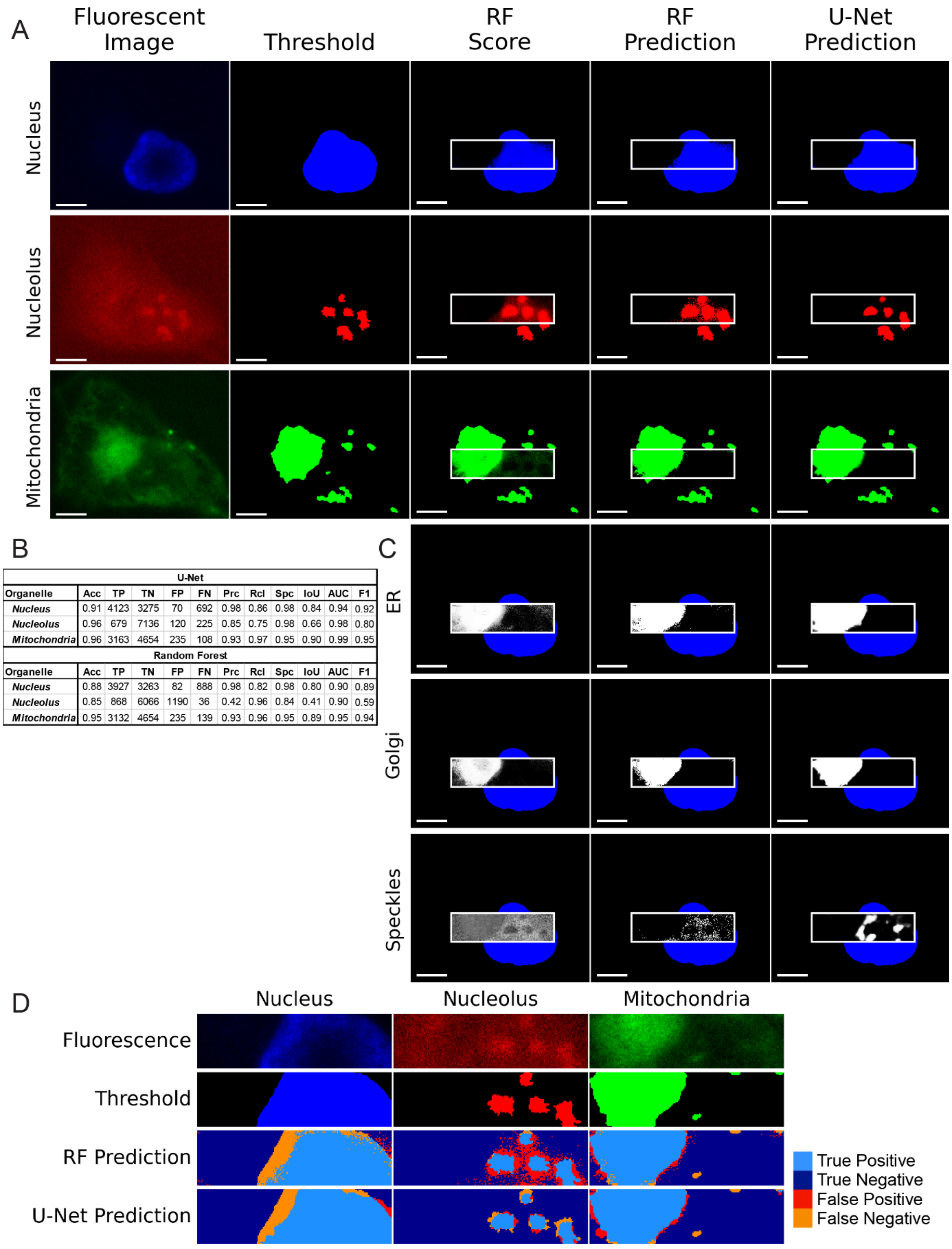


| U-Net | | | | | | | | | | | |
|---|---|---|---|---|---|---|---|---|---|---|---|
| **Organelle** | **Acc** | **TP** | **TN** | **FP** | **FN** | **Prc** | **Rcl** | **Spc** | **IoU** | **AUC** | **F1** |
| ***Nucleus*** | 0.91 | 4123 | 3275 | 70 | 692 | 0.98 | 0.86 | 0.98 | 0.84 | 0.94 | 0.92 |
| ***Nucleolus*** | 0.96 | 679 | 7136 | 120 | 225 | 0.85 | 0.75 | 0.98 | 0.66 | 0.98 | 0.80 |
| ***Mitochondria*** | 0.96 | 3163 | 4654 | 235 | 108 | 0.93 | 0.97 | 0.95 | 0.90 | 0.99 | 0.95 |
| **Random Forest** | | | | | | | | | | | |
| **Organelle** | **Acc** | **TP** | **TN** | **FP** | **FN** | **Prc** | **Rcl** | **Spc** | **IoU** | **AUC** | **F1** |
| ***Nucleus*** | 0.88 | 3927 | 3263 | 82 | 888 | 0.98 | 0.82 | 0.98 | 0.80 | 0.90 | 0.89 |
| ***Nucleolus*** | 0.85 | 868 | 6066 | 1190 | 36 | 0.42 | 0.96 | 0.84 | 0.41 | 0.90 | 0.59 |
| ***Mitochondria*** | 0.95 | 3132 | 4654 | 235 | 139 | 0.93 | 0.96 | 0.95 | 0.89 | 0.95 | 0.94 |

**Figure S20. Partial Cell Hyperspectral Map 4.** A. RF and U-Net predictions for stained organelles. The two leftmost columns display fluorescence images and the fluorescent threshold (scale bars: 10 μm). In columns three-five, a white box is drawn around the region where OPTIR data was collected. The third column shows the random forest confidence scores, which provide information on the certainty of the random forest model. The fourth column shows classification results generated by the random forest model. The final column presents predictions from the U-Net model, which was trained using a combination of random forest scores and IR features. Training dataset sizes are provided in Table S2. Each row shows a different organelle. Row 1: Nucleus (Blue), Row 2: Nucleolus (Red), and Row 3: Mitochondria (green). B. Table of accuracies of the RF and U-Net predictions for stained organelles in A. Accuracy (Acc), True Positive (TP), True Negative (TN), False Positive (FP), and False Negative (FN) are presented first. This is followed by derived performance metrics Precision (Prc), Recall (Rcl), Specificity (Spc), Intersection over Union (IoU), Area under the Curve (AuC), and F1 scores (F1). C. RF and U-Net predictions for unstained organelles in the region OPTIR data was collected. To give spatial context to the location within the cell, the nucleus fluorescence threshold is shown in blue. In line with the columns in A, the third column shows the RF scores, the fourth column shows the RF prediction, and the fifth column shows the U-Net prediction. The predictions are shown in grayscale to distinguish them from organelles with fluorescent comparison. Row 1: ER, Row 2: Golgi, Row 3: Nuclear Speckles**.** D. Mapping classification errors for the stained organelles. To show where errors in each model occur, the cellular region where OPTIR data was collected is enlarged (the regions outlined in white in panel A). The top two rows display the fluorescence and fluorescence threshold from the region. The bottom two rows show the RF and U-Net predictions, colored according to the accuracy of the prediction. Pixels correctly predicted are shown in blue, with light blue for true positives and dark blue for true negatives. Pixels predicted incorrectly are shown in red for false positives and orange for false negatives. The organelles shown here by column are the same organelles as A. Column 1: Nucleus (Blue), Column 2: Nucleolus (Red), and Column 3: Mitochondria (green).

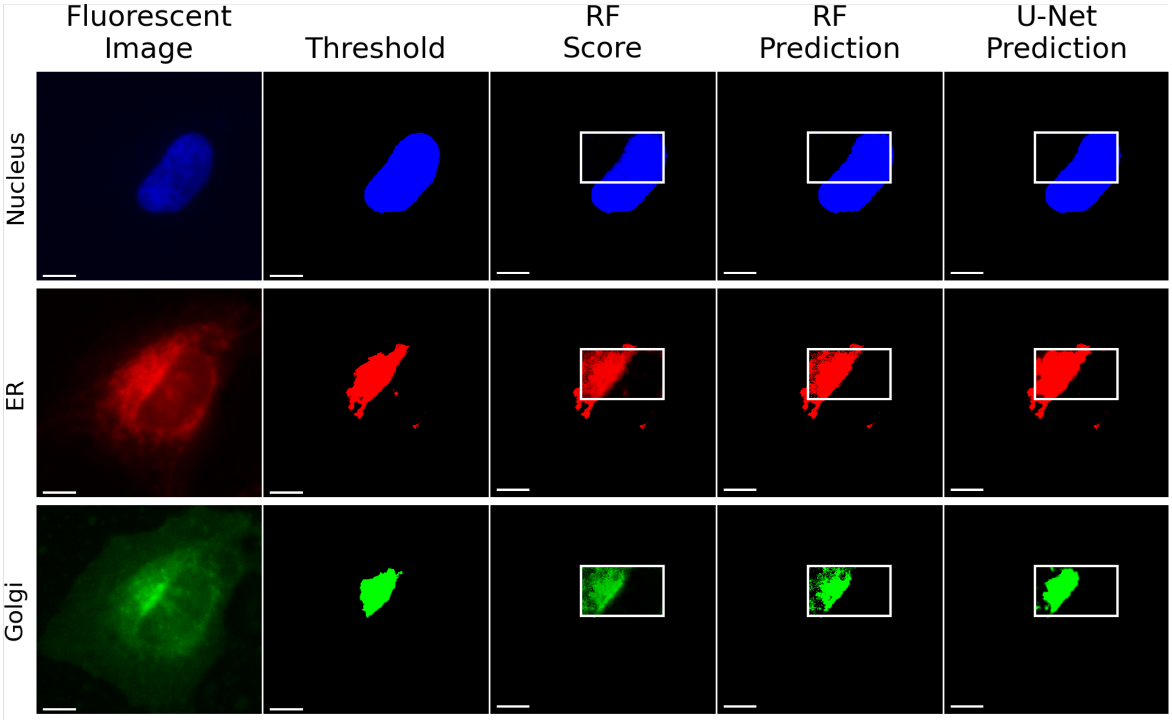


Table S7 - Performance Metrics

| Hyperspectral 5 | | | | | | | | |
|---|---|---|---|---|---|---|---|---|
| **U-Net** | | | | | | | | |
| **Organelle** | **Acc** | **TP** | **TN** | **FP** | **FN** | **IOU** | **AUC** | **F1** |
| ***Nucleus*** | 0.94 | 4890 | 4069 | 300 | 317 | 0.89 | 0.96 | 0.94 |
| ***ER*** | 0.90 | 3313 | 5319 | 763 | 181 | 0.78 | 0.96 | 0.88 |
| ***Golgi*** | 0.92 | 1853 | 6982 | 105 | 636 | 0.71 | 0.97 | 0.83 |
| **Random Forest** | | | | | | | | |
| **Organelle** | **Acc** | **TP** | **TN** | **FP** | **FN** | **IOU** | **AUC** | **F1** |
| ***Nucleus*** | 0.94 | 4842 | 4153 | 216 | 365 | 0.89 | 0.94 | 0.94 |
| ***ER*** | 0.89 | 3042 | 5515 | 567 | 452 | 0.75 | 0.89 | 0.86 |
| ***Golgi*** | 0.86 | 1654 | 6595 | 492 | 835 | 0.55 | 0.80 | 0.71 |

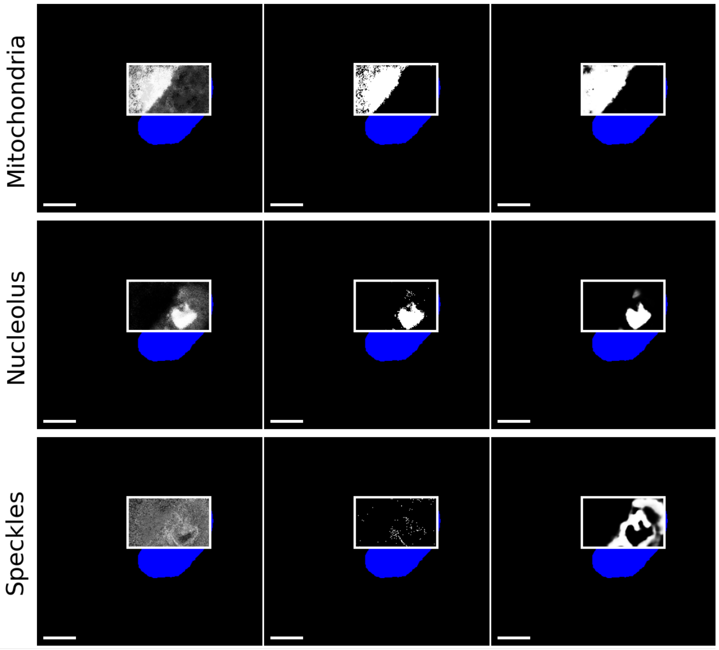


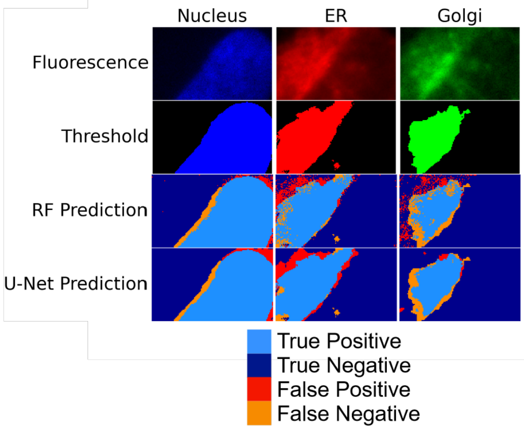


**Figure S21. Partial Cell Hyperspectral Map 5.** A. RF and U-Net predictions for stained organelles. The two leftmost columns display fluorescence images and the fluorescent threshold (scale bars: 10 μm). In columns three-five, a white box is drawn around the region where OPTIR data was collected. The third column shows the random forest confidence scores, which provide information on the certainty of the random forest model. The fourth column shows classification results generated by the random forest model. The final column presents predictions from the U-

Net model, which was trained using a combination of random forest scores and IR features. Training dataset sizes are provided in Table S2. Each row shows a different organelle. Row 1: Nucleus (Blue), Row 2: ER (Red), and Row: Golgi (green). B. Table of Accuracies of the RF and U-Net predictions for stained organelles in A. Accuracy (Acc), True Positive (TP), True Negative (TN), False Positive (FP), and False Negative (FN) are presented first. This is followed by derived performance metrics Precision (Prc), Recall (Rcl), Specificity (Spc), Intersection over Union (IoU), Area under the Curve (AuC), and F1 scores (F1). C. RF and U-Net predictions for unstained organelles in the region OPTIR data was collected. To give spatial context to the location within the cell, the nucleus fluorescence threshold is shown in blue. In line with the columns in A, the third column shows the RF scores, the fourth column shows the RF prediction, and the fifth column shows the U-Net prediction. The predictions are shown in grayscale to distinguish them from organelles with fluorescent comparison. Row 1: Mitochondria, Row 2: Nucleolus, Row 3: Nuclear Speckles. D. Mapping classification errors for the stained organelles. To show where errors in each model occur, the cellular region where OPTIR data was collected is enlarged (the regions outlined in white in panel A). The top two rows display the fluorescence and fluorescence threshold from the region. The bottom two rows show the RF and U-Net predictions, colored according to the accuracy of the prediction. Pixels correctly predicted are shown in blue, with light blue for true positives and dark blue for true negatives. Pixels predicted incorrectly are shown in red for false positives and orange for false negatives. The organelles shown here by column are the same organelles as A. Column 1: Nucleus (Blue), Column 2: ER (Red), and Column: Golgi (green).

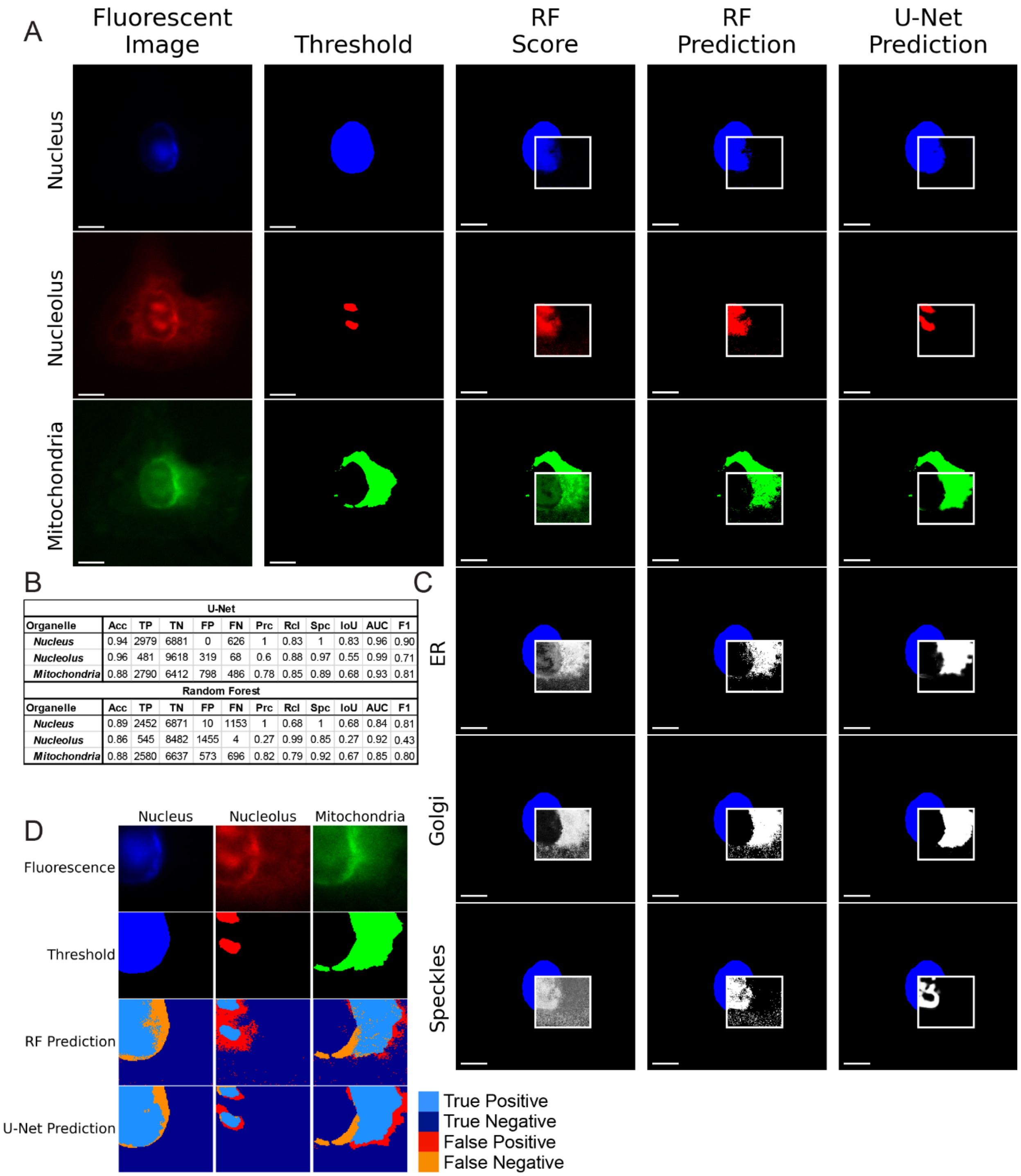


| U-Net | | | | | | | | | | | |
|---|---|---|---|---|---|---|---|---|---|---|---|
| Organelle | Acc | TP | TN | FP | FN | Prc | Rcl | Spc | IoU | AUC | F1 |
| *Nucleus* | 0.94 | 2979 | 6881 | 0 | 626 | 1 | 0.83 | 1 | 0.83 | 0.96 | 0.90 |
| *Nucleolus* | 0.96 | 481 | 9618 | 319 | 68 | 0.6 | 0.88 | 0.97 | 0.55 | 0.99 | 0.71 |
| *Mitochondria* | 0.88 | 2790 | 6412 | 798 | 486 | 0.78 | 0.85 | 0.89 | 0.68 | 0.93 | 0.81 |
| **Random Forest** | | | | | | | | | | | |
| Organelle | Acc | TP | TN | FP | FN | Prc | Rcl | Spc | IoU | AUC | F1 |
| *Nucleus* | 0.89 | 2452 | 6871 | 10 | 1153 | 1 | 0.68 | 1 | 0.68 | 0.84 | 0.81 |
| *Nucleolus* | 0.86 | 545 | 8482 | 1455 | 4 | 0.27 | 0.99 | 0.85 | 0.27 | 0.92 | 0.43 |
| *Mitochondria* | 0.88 | 2580 | 6637 | 573 | 696 | 0.82 | 0.79 | 0.92 | 0.67 | 0.85 | 0.80 |

**Figure S22. Partial Cell Hyperspectral Map 6.** A. RF and U-Net predictions for stained organelles. The two leftmost columns display fluorescence images and the fluorescent threshold (scale bars: 10 μm). In columns three-five, a white box is drawn around the region where OPTIR data was collected. The third column shows the random forest confidence scores, which provide information on the certainty of the random forest model. The fourth column shows classification results generated by the random forest model. The final column presents predictions from the U-

Net model, which was trained using a combination of random forest scores and IR features. Training dataset sizes are provided in Table S2. Each row shows a different organelle. Row 1: Nucleus (Blue), Row 2: Nucleolus (Red), and Row 3: Mitochondria (green). B. Table of accuracies of the RF and U-Net predictions for stained organelles in A. Accuracy (Acc), True Positive (TP), True Negative (TN), False Positive (FP), and False Negative (FN) are presented first. This is followed by derived performance metrics Precision (Prc), Recall (Rcl), Specificity (Spc), Intersection over Union (IoU), Area under the Curve (AuC), and F1 scores (F1). C. RF and U-Net predictions for unstained organelles in the region OPTIR data was collected. To give spatial context to the location within the cell, the nucleus fluorescence threshold is shown in blue. In line with the columns in A, the third column shows the RF scores, the fourth column shows the RF prediction, and the fifth column shows the U-Net prediction. The predictions are shown in grayscale to distinguish them from organelles with fluorescent comparison. Row 1: ER, Row 2: Golgi, Row 3: Nuclear Speckles. D. Mapping classification errors for the stained organelles. To show where errors in each model occur, the cellular region where OPTIR data was collected is enlarged (the regions outlined in white in panel A). The top two rows display the fluorescence and fluorescence threshold from the region. The bottom two rows show the RF and U-Net predictions, colored according to the accuracy of the prediction. Pixels correctly predicted are shown in blue, with light blue for true positives and dark blue for true negatives. Pixels predicted incorrectly are shown in red for false positives and orange for false negatives. The organelles shown here by column are the same organelles as A. Column 1: Nucleus (Blue), Column 2: Nucleolus (Red), and Column 3: Mitochondria (green).

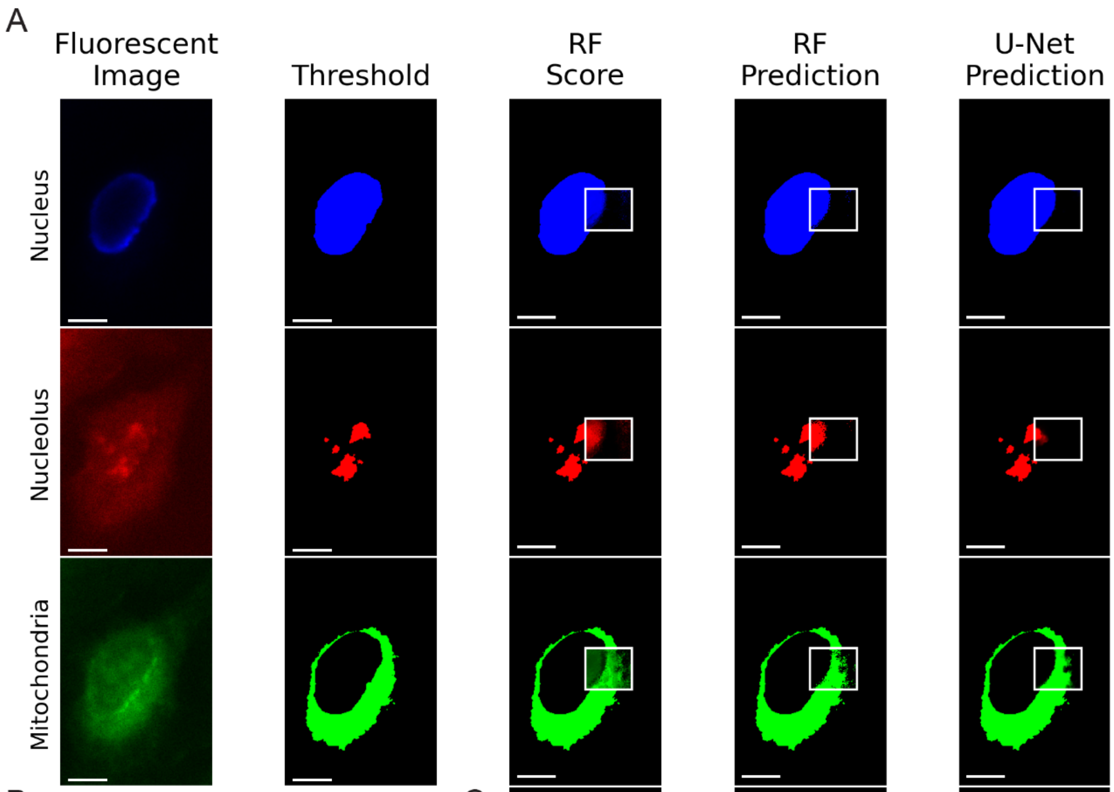


B

| U-Net | | | | | | | | | | | |
|---|---|---|---|---|---|---|---|---|---|---|---|
| Organelle | Acc | TP | TN | FP | FN | Prc | Rcl | Spc | IoU | AUC | F1 |
| ***Nucleus*** | 0.97 | 1067 | 2129 | 66 | 32 | 0.94 | 0.97 | 0.97 | 0.92 | 0.99 | 0.96 |
| ***Nucleolus*** | 0.91 | 1026 | 1966 | 123 | 179 | 0.89 | 0.85 | 0.94 | 0.77 | 0.97 | 0.87 |
| ***Mitochondria*** | 0.91 | 1026 | 1966 | 123 | 179 | 0.89 | 0.85 | 0.94 | 0.77 | 0.97 | 0.87 |
| **Random Forest** | | | | | | | | | | | |
| Organelle | Acc | TP | TN | FP | FN | Prc | Rcl | Spc | IoU | AUC | F1 |
| ***Nucleus*** | 0.96 | 1014 | 2144 | 51 | 85 | 0.95 | 0.92 | 0.98 | 0.88 | 0.95 | 0.94 |
| ***Nucleolus*** | 0.87 | 193 | 2664 | 434 | 3 | 0.31 | 0.98 | 0.86 | 0.31 | 0.92 | 0.47 |
| ***Mitochondria*** | 0.88 | 955 | 1942 | 147 | 250 | 0.87 | 0.79 | 0.93 | 0.71 | 0.86 | 0.83 |

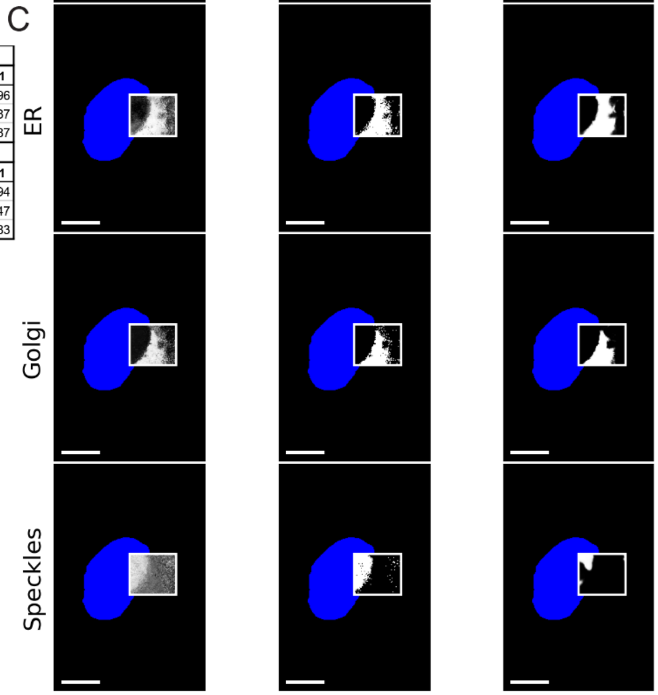


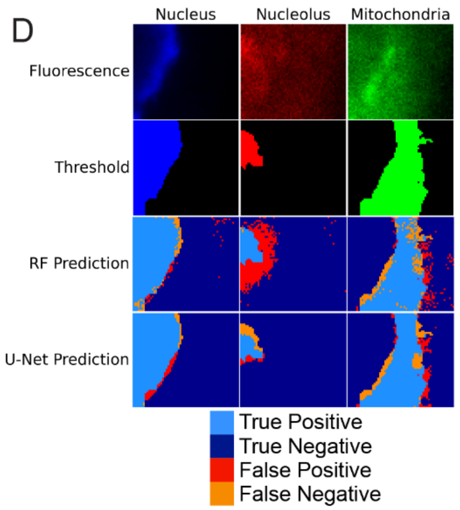


**Figure S23. Partial Cell Hyperspectral Map 7.** A. RF and U-Net predictions for stained organelles. The two leftmost columns display fluorescence images and the fluorescent threshold (scale bars: 10 μm). In columns three-five, a white box is drawn around the region where OPTIR data was collected. The third column shows the random forest confidence scores, which provide information on the certainty of the random forest model. The fourth column shows classification results generated by the random forest model. The final column presents predictions from the U-Net model, which was trained using a combination of random forest scores and IR features. Training dataset sizes are provided in Table S2. Each row shows a different organelle. Row 1: Nucleus (Blue), Row 2: Nucleolus (Red), and Row 3: Mitochondria (green). B. Table of accuracies of the RF and U-Net predictions for stained organelles in A. Accuracy (Acc), True Positive (TP), True Negative (TN), False Positive (FP), and False Negative (FN) are presented first. This is followed by derived performance metrics Precision (Prc), Recall (Rcl), Specificity (Spc), Intersection over Union (IoU), Area under the Curve (AuC), and F1 scores (F1). C. RF and U-Net predictions for unstained organelles in the region OPTIR data was collected. To give spatial context to the location within the cell, the nucleus fluorescence threshold is shown in blue. In line with the columns in A, the third column shows the RF scores, the fourth column shows the RF prediction, and the fifth column shows the U-Net prediction. The predictions are shown in grayscale to distinguish them from organelles with fluorescent comparison. Row 1: ER, Row 2: Golgi, Row 3: Nuclear Speckles. D. Mapping classification errors for the stained organelles. To show where errors in each model occur, the cellular region where OPTIR data was collected is enlarged (the regions outlined in white in panel A). The top two rows display the fluorescence and fluorescence threshold from the region. The bottom two rows show the RF and U-Net predictions, colored according to the accuracy of the prediction. Pixels correctly predicted are shown in blue, with light blue for true positives and dark blue for true negatives. Pixels predicted incorrectly are shown in red for false positives and orange for false negatives. The organelles shown here by column are the same organelles as A. Column 1: Nucleus (Blue), Column 2: Nucleolus (Red), and Column 3: Mitochondria (green).

A

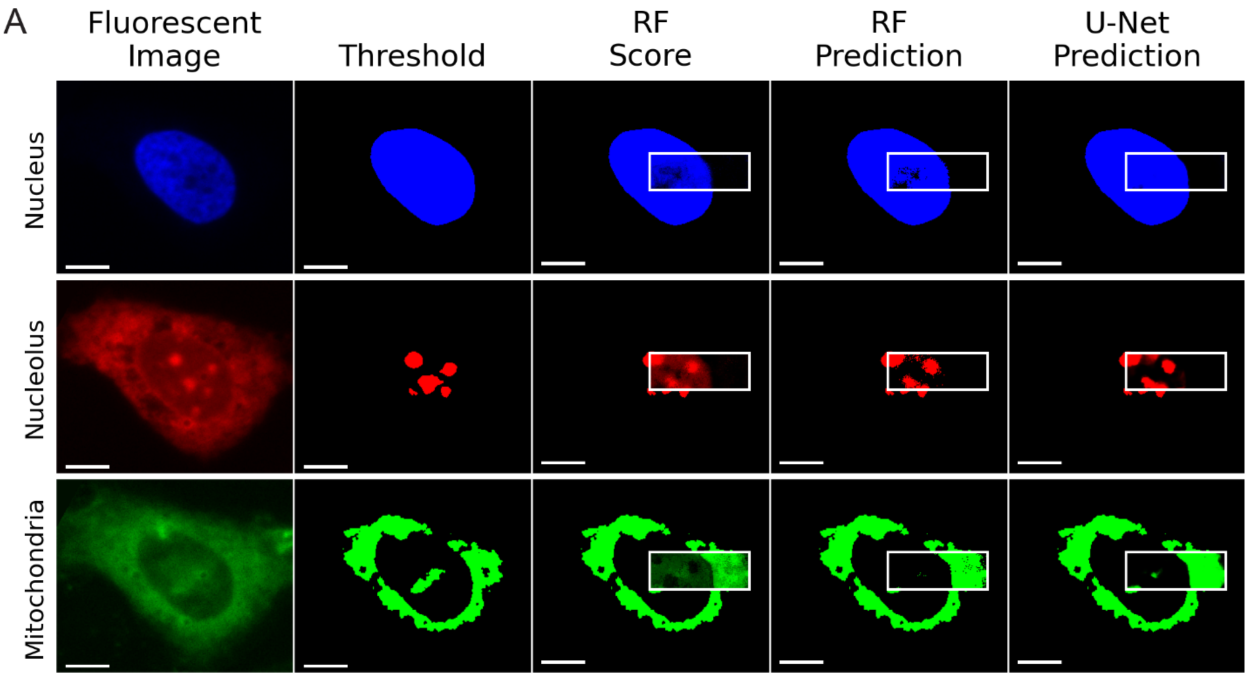


B

| U-Net | | | | | | | | | | | |
|---|---|---|---|---|---|---|---|---|---|---|---|
| Organelle | Acc | TP | TN | FP | FN | Prc | Rcl | Spc | IoU | AUC | F1 |
| *Nucleus* | 0.99 | 2803 | 1990 | 4 | 33 | 1 | 0.99 | 1 | 0.99 | 1.00 | 0.99 |
| *Nucleolus* | 0.85 | 448 | 3677 | 356 | 349 | 0.56 | 0.56 | 0.91 | 0.39 | 0.82 | 0.56 |
| *Mitochondria* | 0.76 | 1145 | 2508 | 663 | 514 | 0.63 | 0.69 | 0.79 | 0.49 | 0.80 | 0.66 |
| **Random Forest** | | | | | | | | | | | |
| Organelle | Acc | TP | TN | FP | FN | Prc | Rcl | Spc | IoU | AUC | F1 |
| *Nucleus* | 0.94 | 2531 | 1991 | 3 | 305 | 1 | 0.89 | 1 | 0.89 | 0.95 | 0.94 |
| *Nucleolus* | 0.86 | 485 | 3645 | 388 | 312 | 0.56 | 0.61 | 0.9 | 0.41 | 0.76 | 0.58 |
| *Mitochondria* | 0.75 | 1070 | 2551 | 620 | 589 | 0.63 | 0.64 | 0.8 | 0.47 | 0.72 | 0.64 |

C

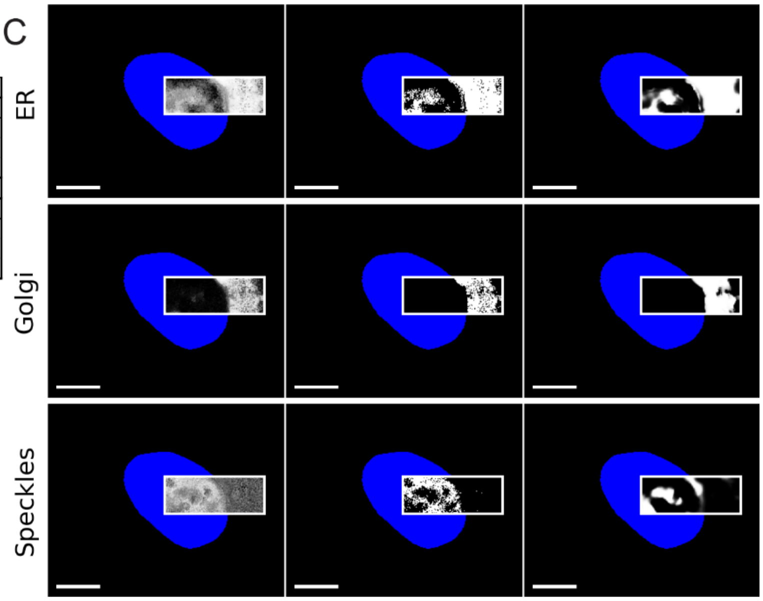


D

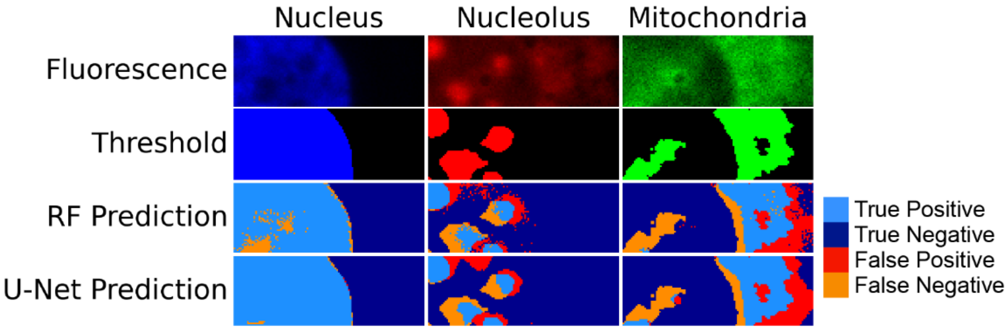


**Figure S24. Partial Cell Hyperspectral Map 8.** A. RF and U-Net predictions for stained organelles. The two leftmost columns display fluorescence images and the fluorescent threshold (scale bars: 10 μm). In columns three-five, a white box is drawn around the region where OPTIR data was collected. The third column shows the random forest confidence scores, which provide information on the certainty of the random forest model. The fourth column shows classification results generated by the random forest model. The final column presents predictions from the U-Net model, which was trained using a combination of random forest scores and IR features. Training dataset sizes are provided in Table S2. Each row shows a different organelle. Row 1: Nucleus (Blue), Row 2: Nucleolus (Red), and Row 3: Mitochondria (green). B. Table of accuracies of the RF and U-Net predictions for stained organelles in A. Accuracy (Acc), True Positive (TP), True Negative (TN), False Positive (FP), and False Negative (FN) are presented first. This is followed by derived performance metrics Precision (Prc), Recall (Rcl), Specificity (Spc), Intersection over Union (IoU), Area under the Curve (AuC), and F1 scores (F1). C. RF and U-Net predictions for unstained organelles in the region OPTIR data was collected. No fluorescence information for the organelles listed is available. To give spatial context to the location within the cell, the nucleus fluorescence threshold is shown in blue. In line with the columns in A, the third column shows the RF scores, the fourth column shows the RF prediction, and the fifth column shows the U-Net prediction. The predictions are shown in grayscale to distinguish them from organelles with fluorescent comparison. Row 1: ER, Row 2: Golgi, Row 3: Nuclear Speckles. D. Mapping classification errors for the stained organelles. To show where errors in each model occur, the cellular region where OPTIR data was collected is enlarged (the regions outlined in white in panel A). The top two rows display the fluorescence and fluorescence threshold from the region. The bottom two rows show the RF and U-Net predictions, colored according to the accuracy of the prediction. Pixels correctly predicted are shown in blue, with light blue for true positives and dark blue for true negatives. Pixels predicted incorrectly are shown in red for false positives and orange for false negatives. The organelles shown here by column are the same organelles as A. Column 1: Nucleus (Blue), Column 2: Nucleolus (Red), and Column 3: Mitochondria (green).

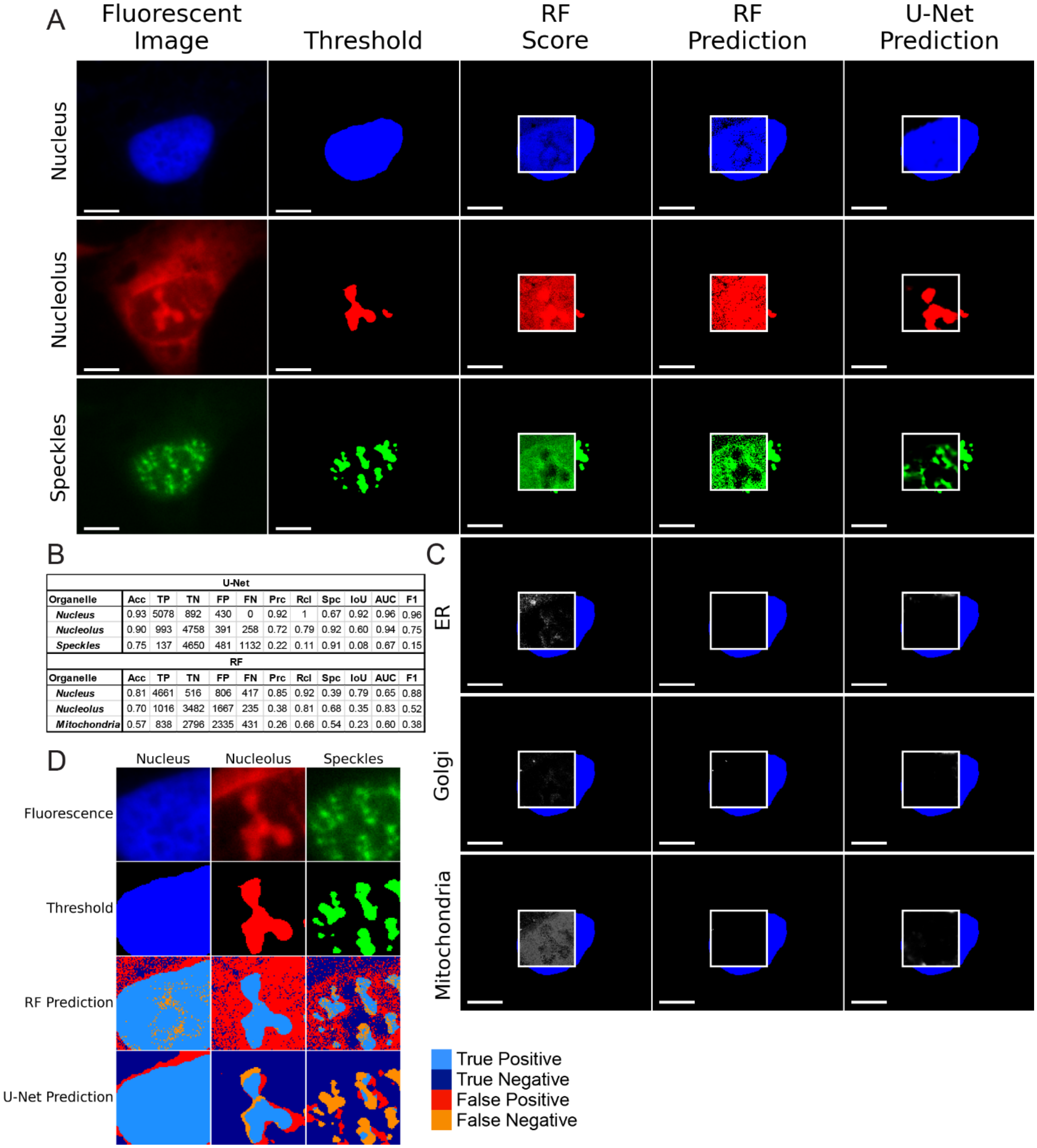


| U-Net | | | | | | | | | | | |
|---|---|---|---|---|---|---|---|---|---|---|---|
| Organelle | Acc | TP | TN | FP | FN | Prc | Rcl | Spc | IoU | AUC | F1 |
| *Nucleus* | 0.93 | 5078 | 892 | 430 | 0 | 0.92 | 1 | 0.67 | 0.92 | 0.96 | 0.96 |
| *Nucleolus* | 0.90 | 993 | 4758 | 391 | 258 | 0.72 | 0.79 | 0.92 | 0.60 | 0.94 | 0.75 |
| *Speckles* | 0.75 | 137 | 4650 | 481 | 1132 | 0.22 | 0.11 | 0.91 | 0.08 | 0.67 | 0.15 |
| **RF** | | | | | | | | | | | |
| Organelle | Acc | TP | TN | FP | FN | Prc | Rcl | Spc | IoU | AUC | F1 |
| *Nucleus* | 0.81 | 4661 | 516 | 806 | 417 | 0.85 | 0.92 | 0.39 | 0.79 | 0.65 | 0.88 |
| *Nucleolus* | 0.70 | 1016 | 3482 | 1667 | 235 | 0.38 | 0.81 | 0.68 | 0.35 | 0.83 | 0.52 |
| *Mitochondria* | 0.57 | 838 | 2796 | 2335 | 431 | 0.26 | 0.66 | 0.54 | 0.23 | 0.60 | 0.38 |

**Figure S25. Partial Cell Hyperspectral Map 9.** A. RF and U-Net predictions for stained organelles. The two leftmost columns display fluorescence images and the fluorescent threshold (scale bars: 10 µm). In columns three-five, a white box is drawn around the region where OPTIR data was collected. The third column shows the random forest confidence scores, which provide information on the certainty of the random forest model. The fourth column shows classification results generated by the random forest model. The final column presents predictions from the U-Net model, which was trained using a combination of random forest scores and IR features. Training dataset sizes are provided in Table S2. Each row shows a different organelle. Row 1:

Nucleus (Blue), Row 2: Nucleolus (Red), and Row 3: Nuclear Speckles (green). B. Table of accuracies of the RF and U-Net predictions for stained organelles in A. Determination of performance is only possible for the stained examples. Details on how these were calculated are detailed in the Model Evaluation section. Accuracy (Acc), True Positive (TP), True Negative (TN), False Positive (FP), and False Negative (FN) are presented first. This is followed by derived performance metrics Precision (Prc), Recall (Rcl), Specificity (Spc), Intersection over Union (IoU), Area under the Curve (AuC), and F1 scores (F1). C. RF and U-Net predictions for unstained organelles in the region OPTIR data was collected. To give spatial context to the location within the cell, the nucleus fluorescence threshold is shown in blue. In line with the columns in A, the third column shows the RF scores, the fourth column shows the RF prediction, and the fifth column shows the U-Net prediction. The predictions are shown in grayscale to distinguish them from organelles with fluorescent comparison. Row 1: ER, Row 2: Golgi, Row 3: Mitochondria. D. Mapping classification errors for the stained organelles. To show where errors in each model occur, the cellular region where OPTIR data was collected is enlarged (the regions outlined in white in panel A). The top two rows display the fluorescence and fluorescence threshold from the region. The bottom two rows show the RF and U-Net predictions, colored according to the accuracy of the prediction. Pixels correctly predicted are shown in blue, with light blue for true positives and dark blue for true negatives. Pixels predicted incorrectly are shown in red for false positives and orange for false negatives. The organelles shown here by column are the same organelles as A. Column 1: Nucleus (Blue), Column 2: Nucleolus (Red), and Column 3: Nuclear Speckles (green).

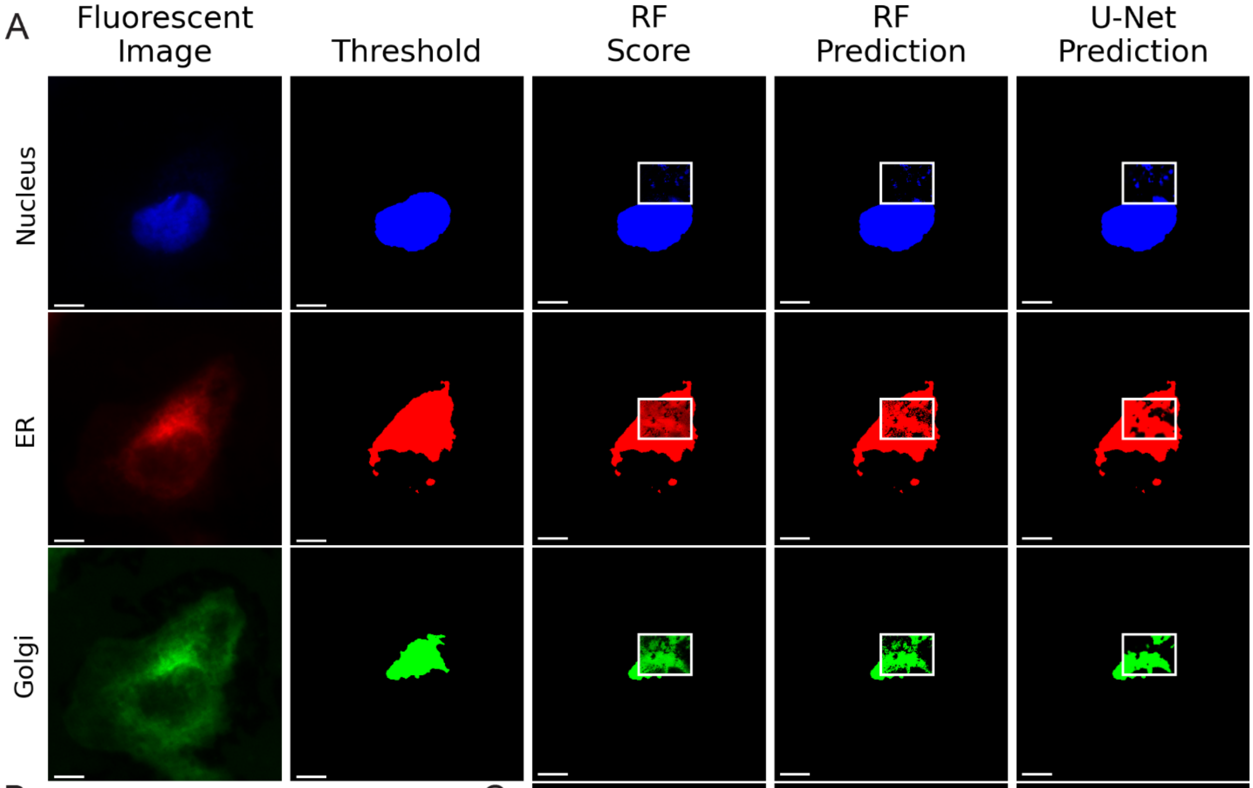


B

| U-Net | | | | | | | | | | | |
|---|---|---|---|---|---|---|---|---|---|---|---|
| Organelle | Acc | TP | TN | FP | FN | Prc | Rcl | Spc | IoU | AUC | F1 |
| ***Nucleus*** | 0.81 | 232 | 4652 | 335 | 833 | 0.41 | 0.22 | 0.93 | 0.17 | 0.54 | 0.28 |
| ***ER*** | 0.65 | 3325 | 631 | 933 | 1163 | 0.78 | 0.74 | 0.4 | 0.61 | 0.61 | 0.76 |
| ***Golgi*** | 0.78 | 1673 | 3019 | 629 | 731 | 0.73 | 0.7 | 0.83 | 0.55 | 0.81 | 0.71 |
| **Random Forest** | | | | | | | | | | | |
| Organelle | Acc | TP | TN | FP | FN | Prc | Rcl | Spc | IoU | AUC | F1 |
| ***Nucleus*** | 0.80 | 115 | 4732 | 255 | 950 | 0.31 | 0.11 | 0.95 | 0.09 | 0.53 | 0.16 |
| ***ER*** | 0.67 | 3282 | 743 | 821 | 1206 | 0.8 | 0.73 | 0.48 | 0.62 | 0.60 | 0.76 |
| ***Golgi*** | 0.74 | 1715 | 2748 | 900 | 689 | 0.66 | 0.71 | 0.75 | 0.52 | 0.73 | 0.68 |

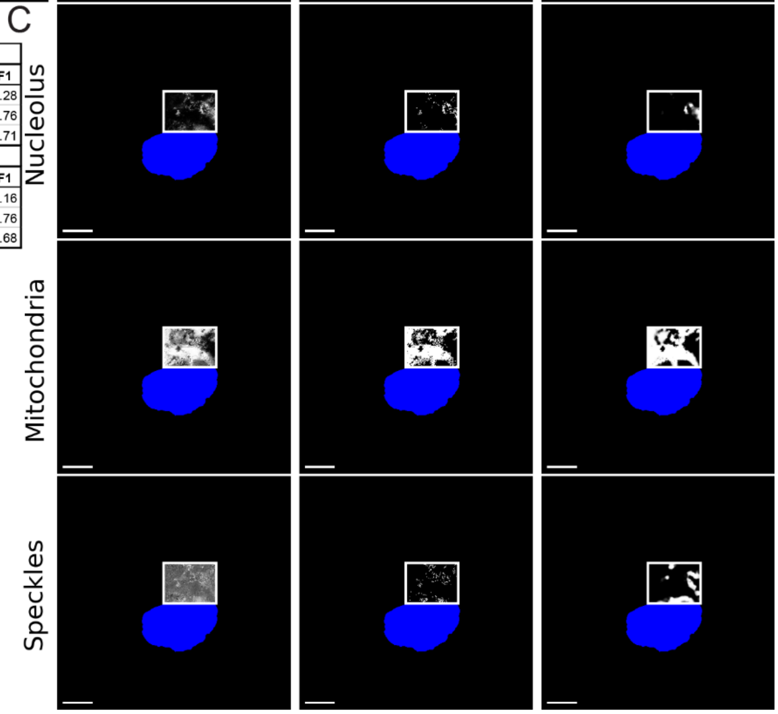


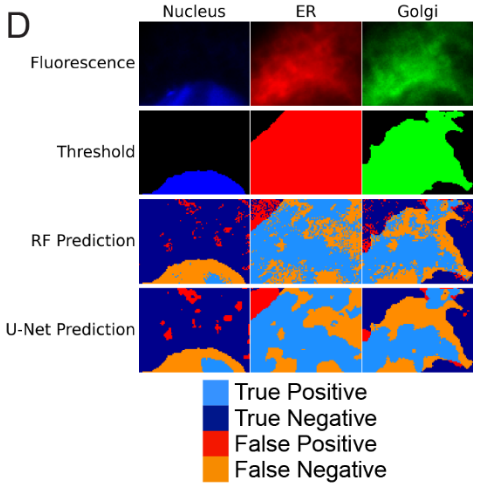


**Figure S26. Partial Cell Hyperspectral Map 10.** A. RF and U-Net predictions for stained organelles. The two leftmost columns display fluorescence images and the fluorescent threshold (scale bars: 10 μm). In columns three-five, a white box is drawn around the region where OPTIR data was collected. The third column shows the random forest confidence scores, which provide information on the certainty of the random forest model. The fourth column shows classification results generated by the random forest model. The final column presents predictions from the U-Net model, which was trained using a combination of random forest scores and IR features. Training dataset sizes are provided in Table S2. Each row shows a different organelle. Row 1: Nucleus (Blue), Row 2: ER (Red), and Row 3: Golgi (green). B. Table of accuracies of the RF and U-Net predictions for stained organelles in A. Accuracy (Acc), True Positive (TP), True Negative (TN), False Positive (FP), and False Negative (FN) are presented first. This is followed by derived performance metrics Precision (Prc), Recall (Rcl), Specificity (Spc), Intersection over Union (IoU), Area under the Curve (AuC), and F1 scores (F1). C. RF and U-Net predictions for unstained organelles in the region OPTIR data was collected. To give spatial context to the location within the cell, the nucleus fluorescence threshold is shown in blue. In line with the columns in A, the third column shows the RF scores, the fourth column shows the RF prediction, and the fifth column shows the U-Net prediction. The predictions are shown in grayscale to distinguish them from organelles with fluorescent comparison. Row 1: Mitochondria, Row 2: Nucleolus, Row 3: Nuclear Speckles. D. Mapping classification errors for the stained organelles. To show where errors in each model occur, the cellular region where OPTIR data was collected is enlarged (the regions outlined in white in panel A). The top two rows display the fluorescence and fluorescence threshold from the region. The bottom two rows show the RF and U-Net predictions, colored according to the accuracy of the prediction. Pixels correctly predicted are shown in blue, with light blue for true positives and dark blue for true negatives. Pixels predicted incorrectly are shown in red for false positives and orange for false negatives. The organelles shown here by column are the same organelles as A. Column 1: Nucleus (Blue), Column 2: ER (Red), and Column 3: Golgi (green).

A

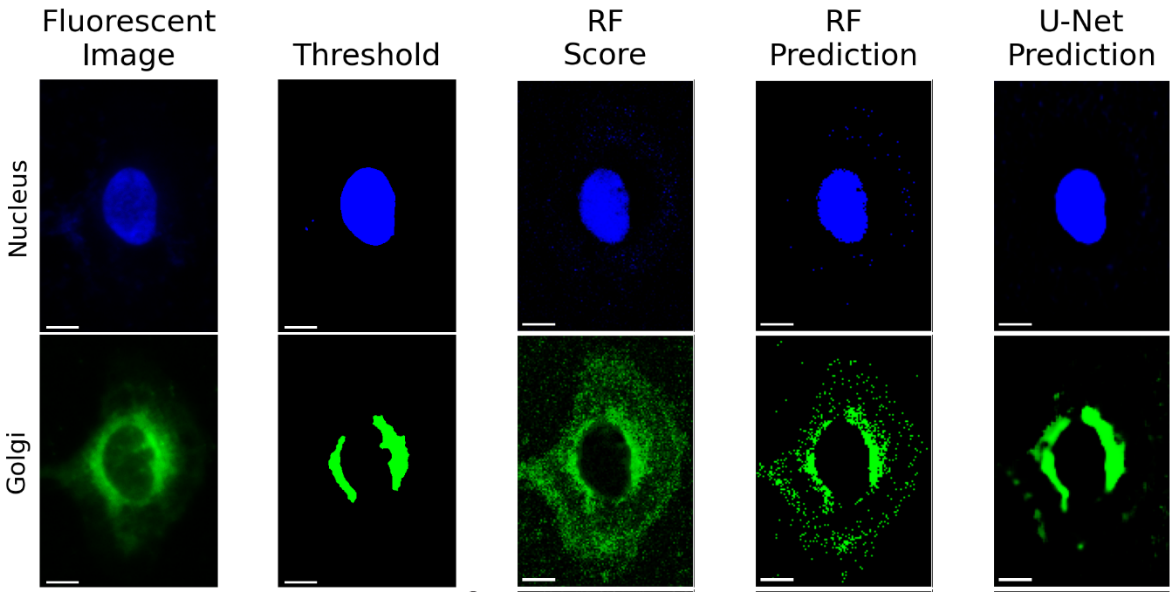


B

| U-Net | | | | | | | | | | | |
|---|---|---|---|---|---|---|---|---|---|---|---|
| **Organelle** | **Acc** | **TP** | **TN** | **FP** | **FN** | **Prc** | **Rcl** | **Spc** | **IoU** | **AUC** | **F1** |
| ***Nucleus*** | 0.99 | 1110 | 15882 | 4 | 164 | 1 | 0.87 | 1 | 0.87 | 0.98 | 0.93 |
| ***Golgi*** | 0.97 | 545 | 16126 | 166 | 323 | 0.77 | 0.63 | 0.99 | 0.53 | 0.81 | 0.69 |
| **Random Forest** | | | | | | | | | | | |
| **Organelle** | **Acc** | **TP** | **TN** | **FP** | **FN** | **Prc** | **Rcl** | **Spc** | **IoU** | **AUC** | **F1** |
| ***Nucleus*** | 0.99 | 1084 | 15837 | 49 | 190 | 0.96 | 0.85 | 1 | 0.82 | 0.92 | 0.90 |
| ***Golgi*** | 0.94 | 485 | 15700 | 592 | 383 | 0.45 | 0.56 | 0.96 | 0.33 | 0.76 | 0.50 |

C

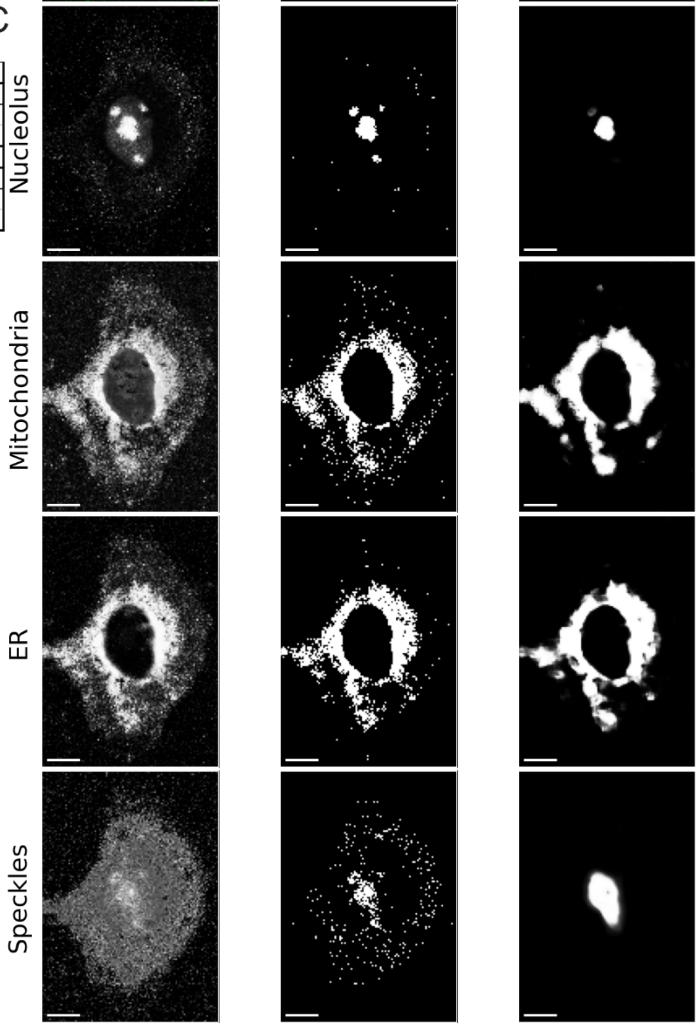


D

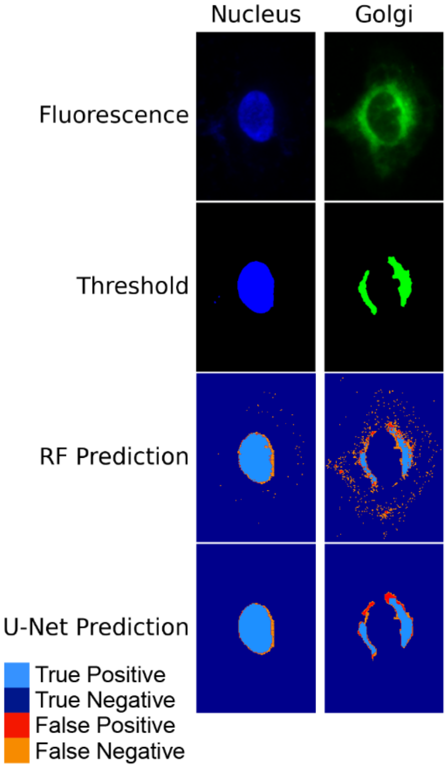

**Figure S27. Whole Cell Hyperspectral Map 1.** A. RF and U-Net predictions for stained organelles. The two leftmost columns display fluorescence images and the fluorescent threshold (scale bars: 10 μm). The third column shows the random forest confidence scores, which provide information on the certainty of the random forest model. The fourth column shows classification results generated by the random forest model. The final column presents predictions from the U-Net model, which was trained using a combination of random forest scores and IR features. Training dataset sizes are provided in Table S2. Each row shows a different organelle. Row 1: Nucleus (Blue) and Row 2: Golgi (green). B. Table of accuracies of the RF and U-Net predictions for stained organelles in A. Accuracy (Acc), True Positive (TP), True Negative (TN), False Positive (FP), and False Negative (FN) are presented first. This is followed by derived performance metrics Precision (Prc), Recall (Rcl), Specificity (Spc), Intersection over Union (IoU), Area under the Curve (AuC), and F1 scores (F1). C. RF and U-Net predictions for unstained organelles. To give spatial context to the location within the cell, the nucleus fluorescence threshold is shown in blue. In line with the columns in A, the third column shows the RF scores, the fourth column shows the RF prediction, and the fifth column shows the U-Net prediction. The predictions are shown in grayscale to distinguish them from organelles with fluorescent comparison. Row 1: Nucleolus, Row 2: ER, Row 3: Mitochondria, Row 4: Nuclear Speckles. D. Mapping classification errors for the stained organelles. To show where errors in each model occur, the cellular region where OPTIR data was collected is shown. The top two rows display the fluorescence and fluorescence threshold from the region. The bottom two rows show the RF and U-Net predictions, colored according to the accuracy of the prediction. Pixels correctly predicted are shown in blue, with light blue for true positives and dark blue for true negatives. Pixels predicted incorrectly are shown in red for false positives and orange for false negatives. The organelles shown here by column are the same organelles as A. Column 1: Nucleus (Blue) and Column 2: Golgi (Green).

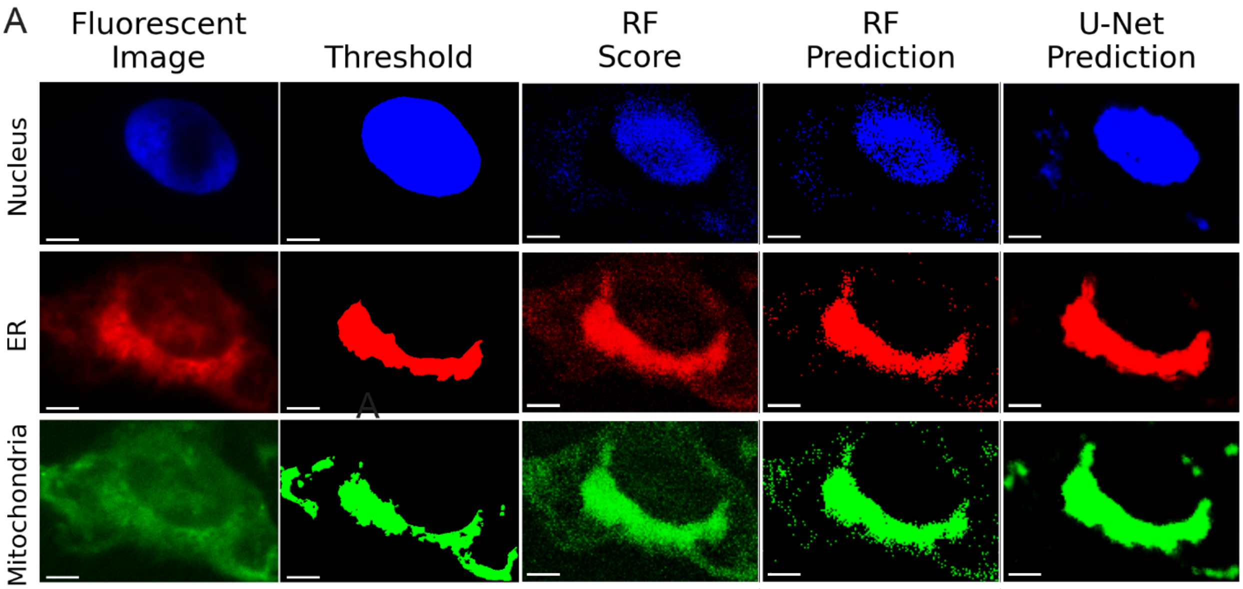


B

| U-Net | | | | | | | | | | | |
|---|---|---|---|---|---|---|---|---|---|---|---|
| Organelle | Acc | TP | TN | FP | FN | Prc | Rcl | Spc | IoU | AUC | F1 |
| *Nucleus* | 0.91 | 2052 | 11277 | 32 | 1239 | 0.98 | 0.62 | 1 | 0.62 | 0.87 | 0.76 |
| *ER* | 0.95 | 1287 | 12576 | 547 | 190 | 0.7 | 0.87 | 0.96 | 0.64 | 0.98 | 0.78 |
| *Mitochondria* | 0.89 | 1165 | 11818 | 996 | 621 | 0.54 | 0.65 | 0.92 | 0.42 | 0.91 | 0.59 |
| RF | | | | | | | | | | | |
| Organelle | Acc | TP | TN | FP | FN | Prc | Rcl | Spc | IoU | AUC | F1 |
| *Nucleus* | 0.88 | 1692 | 11111 | 198 | 1599 | 0.9 | 0.51 | 0.98 | 0.48 | 0.75 | 0.65 |
| *ER* | 0.95 | 1225 | 12595 | 528 | 252 | 0.7 | 0.83 | 0.96 | 0.61 | 0.89 | 0.76 |
| *Mitochondria* | 0.89 | 1094 | 11863 | 951 | 692 | 0.53 | 0.61 | 0.93 | 0.40 | 0.77 | 0.57 |

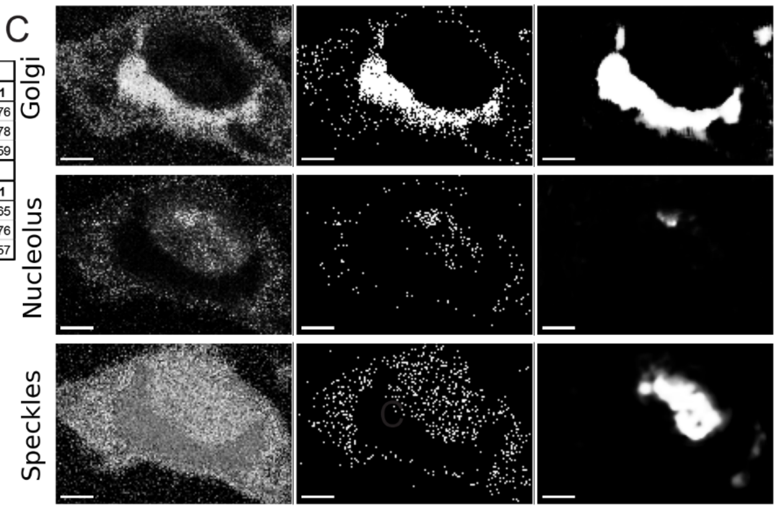


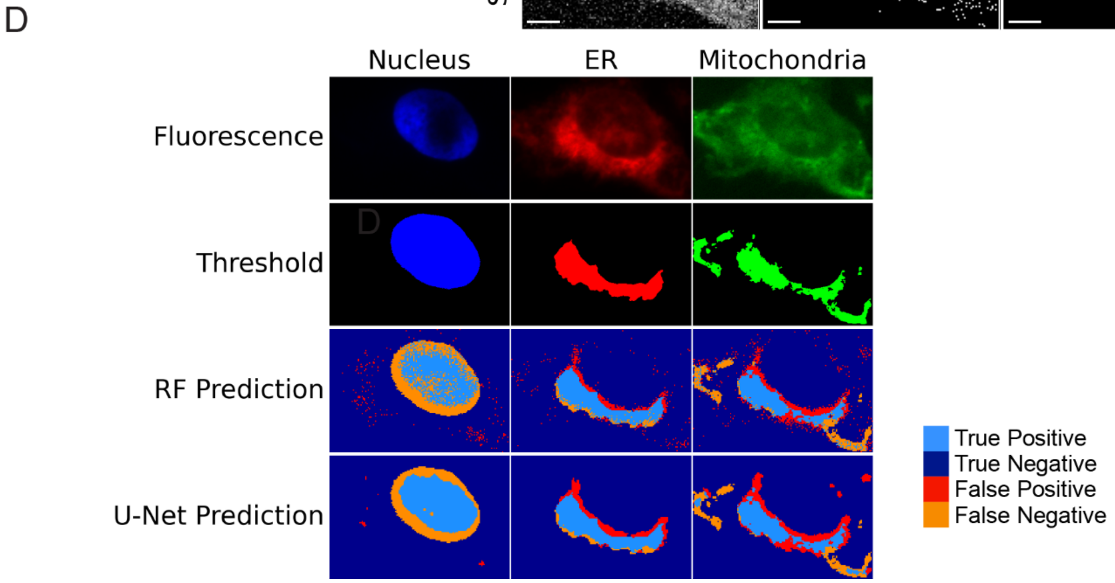

**Figure S28. Whole Cell Hyperspectral Map 2.** A. RF and U-Net predictions for stained organelles. The two leftmost columns display fluorescence images and the fluorescent threshold (scale bars: 10 μm). The third column shows the random forest confidence scores, which provide information on the certainty of the random forest model. The fourth column shows classification results generated by the random forest model. The final column presents predictions from the U-Net model, which was trained using a combination of random forest scores and IR features. Training dataset sizes are provided in Table S2. Each row shows a different organelle. Row 1: Nucleus (Blue), Row 2: ER (Red), and Row 3: Golgi (green). B. Table of accuracies of the RF and U-Net predictions for stained organelles in A. Accuracy (Acc), True Positive (TP), True Negative (TN), False Positive (FP), and False Negative (FN) are presented first. This is followed by derived performance metrics Precision (Prc), Recall (Rcl), Specificity (Spc), Intersection over Union (IoU), Area under the Curve (AuC), and F1 scores (F1). C. RF and U-Net predictions for unstained organelles. To give spatial context to the location within the cell, the nucleus fluorescence threshold is shown in blue. In line with the columns in A, the third column shows the RF scores, the fourth column shows the RF prediction, and the fifth column shows the U-Net prediction. The predictions are shown in grayscale to distinguish them from organelles with fluorescent comparison. Row 1: Mitochondria, Row 2: Nucleolus, Row 3: Nuclear Speckles. D. Mapping classification errors for the stained organelles. To show where errors in each model occur, the cellular region where OPTIR data was collected is shown. The top two rows display the fluorescence and fluorescence threshold from the region. The bottom two rows show the RF and U-Net predictions, colored according to the accuracy of the prediction. Pixels correctly predicted are shown in blue, with light blue for true positives and dark blue for true negatives. Pixels predicted incorrectly are shown in red for false positives and orange for false negatives. The organelles shown here by column are the same organelles as A. Column 1: Nucleus (Blue), Column 2: ER (Red), and Column 3: Golgi (green).

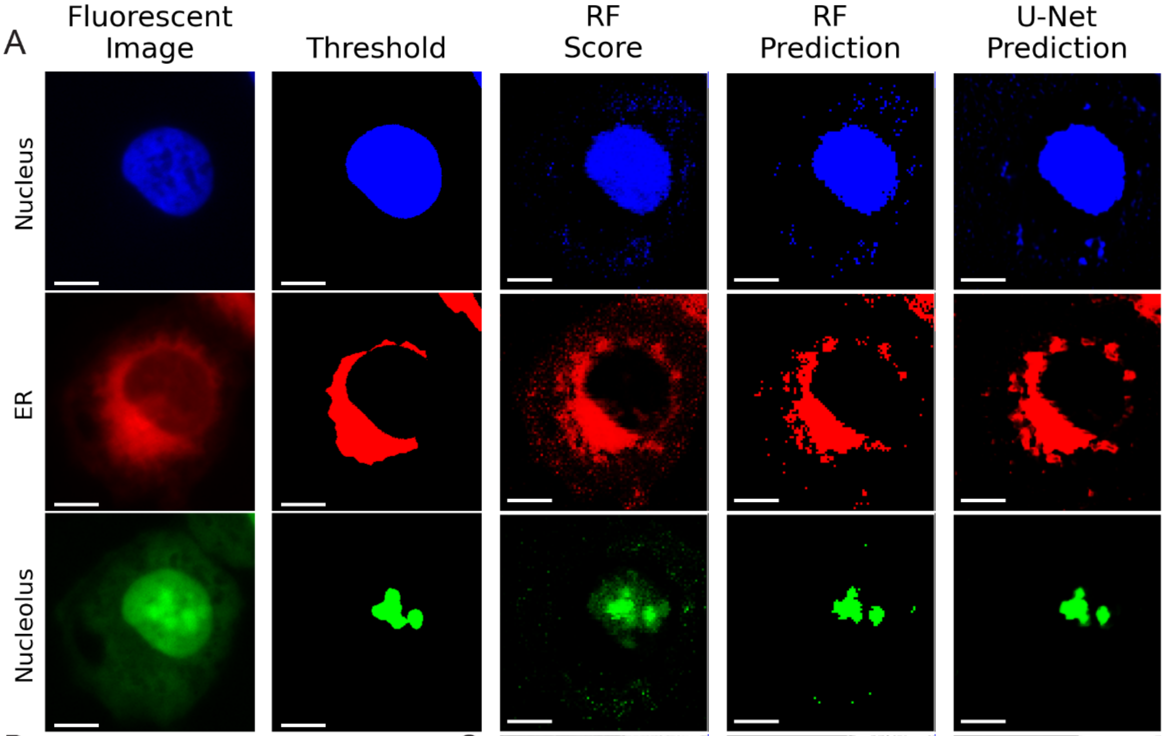


B

| U-Net | | | | | | | | | | | |
|---|---|---|---|---|---|---|---|---|---|---|---|
| Organelle | Acc | TP | TN | FP | FN | Prc | Rcl | Spc | IoU | AUC | F1 |
| ***Nucleus*** | 0.96 | 4353 | 31726 | 390 | 1151 | 0.92 | 0.79 | 0.99 | 0.74 | 0.90 | 0.85 |
| ***ER*** | 0.91 | 2214 | 32161 | 1642 | 1603 | 0.57 | 0.58 | 0.95 | 0.41 | 0.93 | 0.58 |
| ***Nucleolus*** | 0.98 | 441 | 36549 | 208 | 422 | 0.68 | 0.51 | 0.99 | 0.41 | 0.99 | 0.58 |
| **Random Forest** | | | | | | | | | | | |
| Organelle | Acc | TP | TN | FP | FN | Prc | Rcl | Spc | IoU | AUC | F1 |
| ***Nucleus*** | 0.95 | 4274 | 31538 | 578 | 1230 | 0.88 | 0.78 | 0.98 | 0.70 | 0.88 | 0.83 |
| ***ER*** | 0.91 | 2108 | 32227 | 1576 | 1709 | 0.57 | 0.55 | 0.95 | 0.39 | 0.75 | 0.56 |
| ***Nucleolus*** | 0.98 | 442 | 36475 | 282 | 421 | 0.61 | 0.51 | 0.99 | 0.39 | 0.75 | 0.56 |

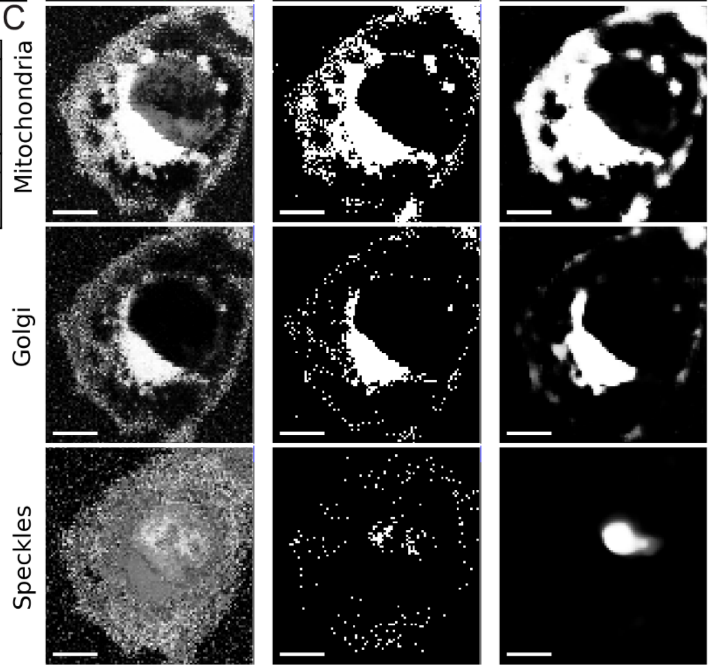


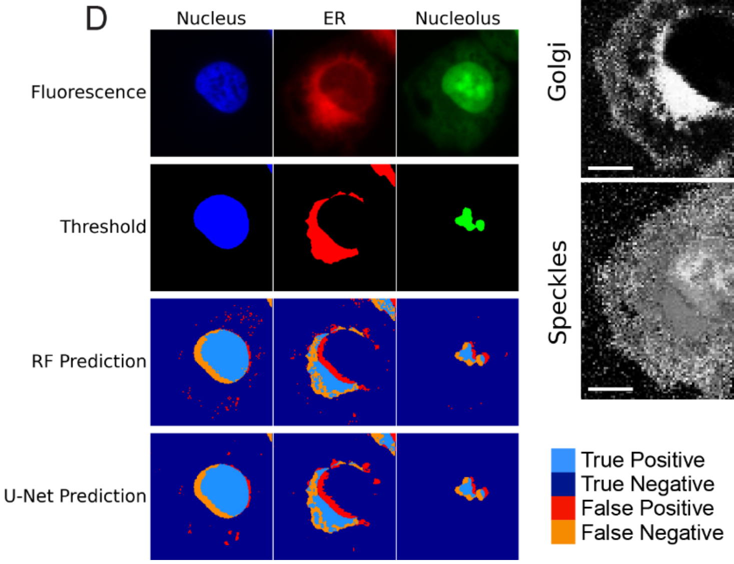


**Figure S29. Whole Cell Hyperspectral Map 3.** A. RF and U-Net predictions for stained organelles. The two leftmost columns display fluorescence images and the fluorescent threshold (scale bars: 10 μm). The third column shows the random forest confidence scores, which provide information on the certainty of the random forest model. The fourth column shows classification results generated by the random forest model. The final column presents predictions from the U-Net model, which was trained using a combination of random forest scores and IR features. Training dataset sizes are provided in Table S2. Each row shows a different organelle. Row 1: Nucleus (Blue), Row 2: ER (Red), and Row 3: Nucleolus (green). B. Table of accuracies of the RF and U-Net predictions for stained organelles in A. Accuracy (Acc), True Positive (TP), True Negative (TN), False Positive (FP), and False Negative (FN) are presented first. This is followed by derived performance metrics Precision (Prc), Recall (Rcl), Specificity (Spc), Intersection over Union (IoU), Area under the Curve (AuC), and F1 scores (F1). C. RF and U-Net predictions for unstained organelles. To give spatial context to the location within the cell, the nucleus fluorescence threshold is shown in blue. In line with the columns in A, the third column shows the RF scores, the fourth column shows the RF prediction, and the fifth column shows the U-Net prediction. The predictions are shown in grayscale to distinguish them from organelles with fluorescent comparison. Row 1: Mitochondria, Row 2: Golgi, Row 3: Nuclear Speckles. D. Mapping classification errors for the stained organelles. To show where errors in each model occur, the cellular region where OPTIR data was collected is shown. The top two rows display the fluorescence and fluorescence threshold from the region. The bottom two rows show the RF and U-Net predictions, colored according to the accuracy of the prediction. Pixels correctly predicted are shown in blue, with light blue for true positives and dark blue for true negatives. Pixels predicted incorrectly are shown in red for false positives and orange for false negatives. The organelles shown here by column are the same organelles as A. Column 1: Nucleus (Blue), Column 2: ER (Red), and Column 3: Nucleolus (green).

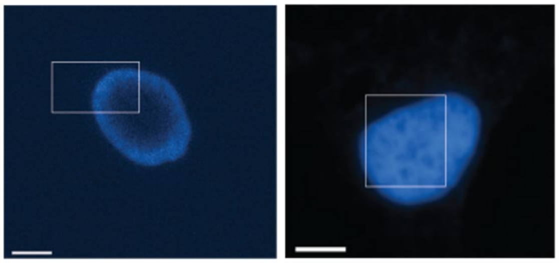

**Figure S30. Example of Different Staining Requiring Different Threshold Procedures to Establish the Ground Truth.** (L*eft*) The nuclear stain does not stain the entire nuclear region. A ring of stain surrounds the inner nucleus. A simple threshold procedure only selects the bright blue regions, which does not align with the nucleus. Additional steps to the threshold procedure are required since the stain does not fully align with the entirety of the nuclear region. The outside ring is first identified and then the region within the stained ring is filled in. (*Right)* An example of a well stained nuclear region which does not require any additional steps to the thresholding procedure.

### Table S1 – Infrared Features Selected as Parameters for Each Organelle

| Parameter Number | $v$ ($cm^{-1}$) | Nucleus | Nucleolus | Speckles | ER | Golgi | Mitochondria | Granules | Class | Assignment |
|---|---|---|---|---|---|---|---|---|---|---|
| 1 | 1040 | ■ | | | ■ | ■ | ■ | ■ | Carbohydrate | C–O stretches of primary alcohols in sugars.[2] |
| 2 | 1060 | | ■ | ■ | | | | | Nucleic Acids | C-O deoxyribose stretch[3] |
| 3 | 1070 | | ■ | ■ | | | | | Nucleic Acids | Nucleic acid bands[4,5] |
| 4 | 1063, 1087 | ■ | | | ■ | ■ | | | Nucleic Acids | C-O stretching C-O ribose[6] |
| 5 | 1155 | | | | ■ | ■ | ■ | | Carbohydrate | C-O stretches of alcohol[7] |
| 6 | 1170 | | | | ■ | ■ | ■ | ■ | Carbohydrate | C-O bands from secondary alcohols |
| 7 | 1390 | ■ | ■ | ■ | | | ■ | | Protein | Carboxylate[8] |
| 8 | 1450, 1464 | ■ | ■ | ■ | ■ | ■ | ■ | ■ | Lipid | CH2 Scissoring Mode[9,10] |
| 9 | 1541 | | ■ | ■ | | | | | Protein | Beta Sheet of Amide II[11] |
| 10 | Amide 2: Amide 1 | ■ | | | ■ | ■ | ■ | ■ | Protein | Protein Populations [11] |
| 11 | FHWM | ■ | ■ | ■ | ■ | ■ | ■ | ■ | Protein | Protein Populations[11] |
| 12 | 1643 | | | | ■ | ■ | ■ | ■ | Protein | Beta Sheet Content[11] |
| 13 | 1656 | | ■ | | | | | | Protein | Alpha Helical Content[11] |
| 14 | 1643, 1656 | | | ■ | | | | | Protein | Alpha : Beta Sheet Content Ratio[11] |
| 15 | 1690 | | | | | | ■ | | Protein | Beta Turn[11] |
| 16 | 1738 | ■ | | | ■ | ■ | ■ | ■ | Lipid | Lipids[12] |

**Table S2 – Training Data Set Sizes**

| Organelle | Spectra | Number of Cells | Positive | Negative | |
|---|---|---|---|---|---|
| | | | Organelle of Interest | Cytoplasm | Nucleoplasm |
| **Nucleus** | 7031 | 55 | 3020 | 4011 | N/A |
| **Nucleolus** | 3250 | 33 | 1400 | 724 | 1126 |
| **ER** | 4580 | 34 | 1950 | 1376 | 1254 |
| **Golgi** | 1230 | 15 | 613 | 382 | 235 |
| **Mitochondria** | 750 | 14 | 375 | 250 | 124 |
| **Speckles** | 3972 | 28 | 1225 | 941 | 1806 |
| **Granules** | 5175 | 27 | 1950 | 1488 | 1737 |

### Supplementary Information References


1. Foucart, A., Debeir, O. & Decaestecker, C. Panoptic quality should be avoided as a metric for assessing cell nuclei segmentation and classification in digital pathology. *Sci Rep* **13**, 8614 (2023).
2. Coblentz Society & Stein, S. E. D-glucose anhydrous in Evaluated Infrared Reference Spectra. in *NIST Chemistry WebBook, NIST Standard Reference Database 69* vol. 20899 (National Institute of Standards and Technology).
3. Duan, M., Li, Y., Zhang, F. & Huang, Q. Assessing B-Z DNA Transitions in Solutions via Infrared Spectroscopy. *Biomolecules* **13**, 964 (2023).
4. Stone, N., Kendall, C., Smith, J., Crow, P. & Barr, H. Raman spectroscopy for identification of epithelial cancers. *Faraday Disc.* **126**, 141 (2004).
5. Wood, B. R., Quinn, M. A., Burden, F. R. & McNaughton, D. An investigation into FTIR spectroscopy as a biodiagnostic tool for cervical cancer. *Biospectroscopy* **2**, 143–153 (1998).
6. Dovbeshko, G. FTIR spectroscopy studies of nucleic acid damage. *Talanta* **53**, 233–246 (2000).
7. Wang, H. P., Wang, H.-C. & Huang, Y.-J. Microscopic FTIR studies of lung cancer cells in pleural fluid. *Science of The Total Environment* **204**, 283–287 (1997).
8. Max, J.-J. & Chapados, C. Infrared Spectroscopy of Aqueous Carboxylic Acids: Comparison between Different Acids and Their Salts. *J. Phys. Chem. A* **108**, 3324–3337 (2004).
9. Liu, Y. & Lunter, D. J. Selective and sensitive spectral signals on confocal Raman spectroscopy for detection of ex vivo skin lipid properties. *Transl Biophotonics* **2**, e202000003 (2020).
10. Rohman, A. *et al.* The use of FTIR and Raman spectroscopy in combination with chemometrics for analysis of biomolecules in biomedical fluids: A review. *BSI* **8**, 55–71 (2020).
11. Barth, A. & Zscherp, C. What vibrations tell about proteins. *Quart. Rev. Biophys.* **35**, 369–430 (2002).
12. Derenne, A., Claessens, T., Conus, C. & Goormaghtigh, E. Infrared Spectroscopy of Membrane Lipids. in *Encyclopedia of Biophysics* (ed. Roberts, G. C. K.) 1074–1081 (Springer Berlin Heidelberg, Berlin, Heidelberg, 2013). doi:10.1007/978-3-642-16712-6_558.
